# Evaluating the informativeness of deep learning annotations for human complex diseases

**DOI:** 10.1101/784439

**Authors:** Kushal K. Dey, Bryce Van de Geijn, Samuel Sungil Kim, Farhad Hormozdiari, David R. Kelley, Alkes L. Price

**Affiliations:** Department of Epidemiology, Harvard T. H. Chan School of Public Health, Boston, MA; Department of Biostatistics, Harvard T.H. Chan School of Public Health, Boston, MA; Department of Electrical Engineering and Computer Science, Massachusetts Institute of Technology, Cambridge, MA; Calico Labs. South San Francisco, CA

## Abstract

Deep learning models have shown great promise in predicting genome-wide regulatory effects from DNA sequence, but their informativeness for human complex diseases and traits is not fully understood. Here, we evaluate the disease informativeness of allelic-effect annotations (absolute value of the predicted difference between reference and variant alleles) constructed using two previously trained deep learning models, DeepSEA and Basenji. We apply stratified LD score regression (S-LDSC) to 41 independent diseases and complex traits (average N=320K) to evaluate each annotation’s informativeness for disease heritability conditional on a broad set of coding, conserved, regulatory and LD-related annotations from the baseline-LD model and other sources; as a secondary metric, we also evaluate the accuracy of models that incorporate deep learning annotations in predicting disease-associated or fine-mapped SNPs. We aggregated annotations across all tissues (resp. blood cell types or brain tissues) in meta-analyses across all 41 traits (resp. 11 blood-related traits or 8 brain-related traits). These allelic-effect annotations were highly enriched for disease heritability, but produced only limited conditionally significant results – only Basenji-H3K4me3 in meta-analyses across all 41 traits and brain-specific Basenji-H3K4me3 in meta-analyses across 8 brain-related traits. We conclude that deep learning models are yet to achieve their full potential to provide considerable amount of unique information for complex disease, and that the informativeness of deep learning models for disease beyond established functional annotations cannot be inferred from metrics based on their accuracy in predicting regulatory annotations.

## Introduction

Disease risk variants identified by genome-wide association studies (GWAS) lie predominantly in non-coding regions of the genome^1,2,3,4,5,6,7^, motivating broad efforts to generate genome-wide maps of regulatory marks across tissues and cell types^8,9,10,11^. Recently, deep learning models trained using these genome-wide maps have shown considerable promise in predicting regulatory marks directly from DNA sequence^12,13,14,15,16,17,18^. In particular, these studies showed that variant-level deep learning annotations (predictive annotations based on the reference allele) attained high accuracy in predicting the underlying chromatin marks^13,14,15,16^, and that models incorporating allelic-effect deep learning annotations (absolute value of the predicted difference between reference and variant alleles) attained high accuracy in predicting disease-associated SNPs^13,14,15,16^. Additional applications of deep learning models, including analyses of signed allelic-effect annotations, are discussed in the Discussion section. However, it is unclear whether deep learning annotations at commonly varying SNPs contain unique information for complex disease that is not present in other annotations.

Here, we evaluate the informativeness for complex disease of allelic-effect annotations at commonly varying SNPs constructed using two deep learning models previously trained on tissue-specific regulatory features (DeepSEA^13, 15^ and Basenji^16^). We apply stratified LD score regression^5, 19^ (S-LDSC) to 41 independent diseases and complex traits (average *N* =320K) to evaluate each annotation’s informativeness for disease heritability conditional on the underlying variant-level annotations as well as a broad set of coding, conserved, regulatory and LD-related annotations from the baseline-LD model^19^ and other sources (imputed Roadmap and ChromHMM annotations^11, 20,21,22^). As a secondary metric, we also evaluate the accuracy of models that incorporate deep learning annotations in predicting disease-associated or fine-mapped SNPs^23, 24^. We aggregate DeepSEA and Basenji annotations across all tissues in meta-analyses across all 41 traits, across blood cell types in meta-analyses across 11 blood-related traits, and across brain tissues in meta-analyses across 8 brain-related traits.

## Results

### Overview of methods

We define a genomic annotation as an assignment of a numeric value (either binary or continuous-valued) to each SNP (Methods). Our focus is on continuous-valued annotations (with values between 0 and 1) trained by deep learning models to predict biological function from DNA sequence. Annotation values are defined for each SNP with minor allele count ≥ 5 in a 1000 Genomes Project European reference panel^25^, as in our previous work^5^. We have publicly released all new annotations analyzed in this study (see URLs).

In our analysis of allelic-effect (Δ) deep learning annotations across 41 traits, we analyzed 16 non-tissue-specific deep learning annotations: 8 DeepSEA annotations^13, 15^ previously trained to predict 4 tissue-specific chromatin marks (DNase, H3K27ac, H3K4me1, H3K4me3) known to be associated with active promoter and enhancer regions across 127 Roadmap tissues^11, 26^, aggregated using the average (Avg) or maximum (Max) across tissues, and 8 analogous Basenji annotations^16^, quantile-matched with DeepSEA annotations to lie between 0 and 1 (Table 1 and Methods). To assess whether the allelic-effect annotations provided unique information for disease, we conservatively included the underlying variant-level (V) annotations (Table S1) as well as a broad set of coding, conserved, regulatory and LD-related annotations in our analyses: 86 annotations from the baseline-LD (v2.1) model^19^, which has been shown to effectively model LD-dependent architectures^27^; 8 Roadmap annotations^11^ (for same chromatin marks as DeepSEA and Basenji annotations), imputed using ChromImpute^20^; and 40 ChromHMM annotations^21, 22^ based on 20 ChromHMM states across 127 Roadmap tissues^11^ (Table S2). When comparing pairs of annotations that differed only in their aggregation strategy (Avg/Max), chromatin mark (DNase/H3K27ac/H3K4me1/H3K4me3), model (DeepSEA/Basenji) or type (variant-level/allelic-effect), respectively, we observed large correlations across aggregation strategies (average *r* = 0.71), chromatin marks (average *r* = 0.58), models (average *r* = 0.54) and types (average *r* = 0.48) (Figure S1).

**Table 1.**
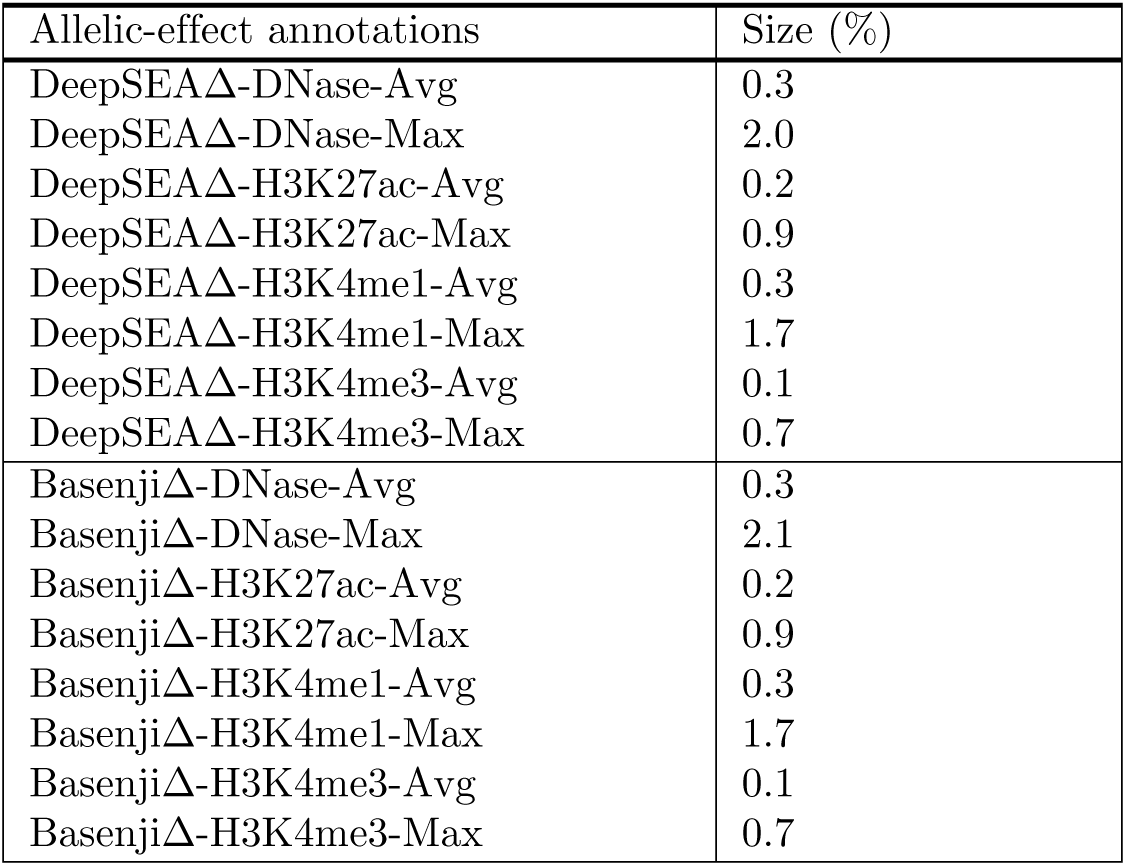
**List of non-tissue-specific allelic-effect analyzed.** We list the 16 allelic-effect deep learning annotations (8 DeepSEAΔ, 8 BasenjiΔ) and their annotation sizes (average annotation value across SNPs). A list of non-tissue-specific variant-level annotations is provided in Table S1.

In our analysis of 11 blood-related traits (resp. 8 brain-related traits), we analyzed 8 DeepSEA annotations and 8 Basenji annotations that were aggregated across 27 blood cell types (resp. 13 brain tissues), instead of all 127 tissues. Details of other annotations included in these analyses are provided below.

We assessed the informativeness of these annotations for disease heritability using stratified LD score regression (S-LDSC) with the baseline-LD model^5, 19^. We considered two metrics, enrichment and standardized effect size (*τ\**). Enrichment is defined as the proportion of heritability explained by SNPs in an annotation divided by the proportion of SNPs in the annotation^5^, and generalizes to continuous-valued annotations with values between 0 and 1^28^. Standardized effect size (*τ\**) is defined as the proportionate change in per-SNP heritability associated with a 1 standard deviation increase in the value of the annotation, conditional on other annotations included in the model^19^; unlike enrichment, *τ^*^*quantifies effects that are unique to the focal annotation. In our “marginal” analyses, we estimated *τ\** for each focal annotation conditional on annotations from the baseline-LD model. In our “joint” analyses, we merged baseline-LD model annotations with focal annotations that were marginally significant after Bonferroni correction and performed forward stepwise elimination to iteratively remove focal annotations that had conditionally non-significant *τ\** values after Bonferroni correction, as in ref.^19^. All analyses of allelic-effect annotations were further conditioned on jointly significant annotations from a variant-level analysis, if any. Distinct from evaluating deep learning annotations using S-LDSC, we also evaluated the accuracy of models that incorporate deep learning annotations in predicting disease-associated or fine-mapped SNPs^23, 24^ (Methods).

### A Basenji all-tissues allelic-effect annotation is conditionally informative for disease

We evaluated the informativeness of allelic-effect deep learning annotations for disease heritability by applying S-LDSC with the baseline-LD model^5, 19^ to summary association statistics for 41 independent diseases and complex traits (average *N* =320K); for 6 traits we analyzed two different data sets, leading to a total of 47 data sets analyzed (Table S3). We meta-analyzed results across these 47 data sets, which were chosen to be independent^28^. The 41 traits include 27 UK Biobank traits^29^ for which summary association statistics are publicly available (see URLs).

Although our main focus is on allelic-effect deep learning annotations, analysis of variant-level deep learning annotations was a necessary prerequisite step, for two reasons: (i) allelic-effect annotations are computed as differences between variant-level annotations for each allele, and (ii) we wished to condition analyses of allelic-effect annotations on jointly significant variant-level annotations, if any. We thus constructed 8 variant-level DeepSEAV annotations by applying previously trained DeepSEA models^15^ (see URLs) for each of 4 tissue-specific chromatin marks (DNase, H3K27ac, H3K4me1, H3K4me3) across 127 Roadmap tissues^11^ to 1kb of human reference sequence around each SNP; for each chromatin mark, we aggregated variant-level DeepSEAV annotations across the 127 tissues using either the average (Avg) or maximum (Max) across tissues (Table 1 and Methods). The DeepSEA model was highly predictive of the corresponding tissue-specific chromatin marks, with AUROC values reported by ref.^15^ ranging from 0.77 − 0.97 (Table S4). We also constructed 8 variant-level BasenjiV annotations by applying previously trained Basenji models^16^ (see URLs) and aggregating across tissues in analogous fashion (Table 1 and Methods); Basenji uses a Poisson likelihood model, unlike the binary classification approach of DeepSEA, and analyzes 130kb of human reference sequence around each SNP using dilated convolutional layers. The constituent tissue-specific BasenjiV annotations do not lie between 0 and 1; so we transformed these annotations to lie between 0 and 1 via quantile matching with corresponding DeepSEAV annotations, to ensure a fair comparison of the two approaches (Methods). Although the variant-level DeepSEAV and BasenjiV annotations were highly enriched for heritability, we determined that none of them were conditionally informative across the 41 traits (Figures S2, S3, S4, S5, S6 and Supplementary Note). This is an expected result, because the variant-level deep learning annotations simply predict measured variant-level annotations from Roadmap that are also included in the model.

Our main focus is on allelic-effect annotations (absolute value of the predicted difference between reference and variant alleles), which have been the focus of recent work^13,14,15,16^. We evaluated the informativeness of 8 non-tissue-specific DeepSEAΔ and 8 non-tissue-specific BasenjiΔ allelic-effect annotations (Table 1) for disease heritability by applying S-LDSC to the 41 traits. Analyses of allelic-effect annotations were conditioned on the baseline-LD model plus 7 annotations from Figure S6. For ease of comparison, allelic-effect Basenji annotations were quantile-matched with corresponding allelic-effect DeepSEA annotations, analogous to analyses of variant-level annotations.

A summary of the results is provided in Figure 1 (All tissues, All traits column; numerical results in Table S5), which reports the number of allelic-effect annotations of various types with significant heritability enrichment, marginal conditional signal, and joint conditional signal, respectively. In our marginal analysis of disease heritability, all allelic-effect annotations from DeepSEA and Basenji models were significantly enriched for heritability across 41 traits; the allelic-effect BasenjiΔ annotations were more enriched for disease heritability (2.40x) than allelic-effect DeepSEAΔ annotations (1.91x) (Table S6). However, only 0 DeepSEAΔ annotations and 1 BasenjiΔ annotation, BasenjiΔ-H3K4me3-Max, attained a Bonferroni-significant standardized effect size (*τ\**) (Figure 2 and Table S6); results were similar when conditioned on just the baseline-LD model (Table S7). Despite the high correlation between variant-level and allelic-effect annotations (*r* = 0.48; Figure S1), the corresponding variant-level annotation (BasenjiV-H3K4me3-Max) did not produce significant conditional signal (Figure 2 and Table S8), consistent with Figure S2). We note that since BasenjiΔ-H3K4me3-Max was the only marginally significant annotation in the non-tissue-specific allelic-effect analysis, it is automatically jointly significant.

**Figure 1.**
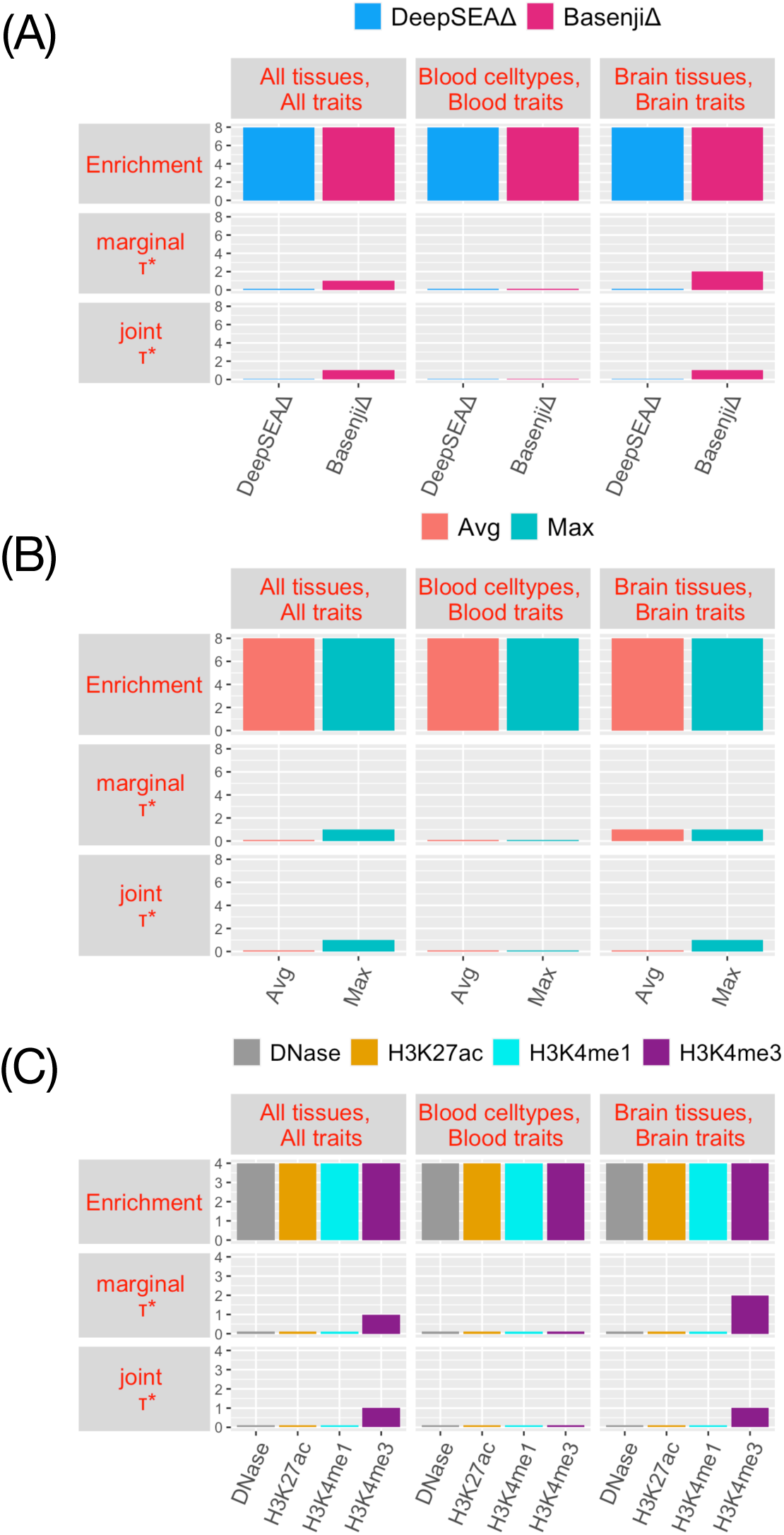
**Summary of disease informativeness of allelic-effect deep learning annotations**: We report the number of allelic-effect annotations with significant heritability enrichment, marginal conditional *τ\**, and joint conditional *τ\**, across (A) different deep learning models (DeepSEA/Basenji), (B) different aggregation strategies (Avg/Max) and (C) different chromatin marks (DNase/H3K27ac/H3K4me1/H3K4me3). Numerical results are reported in Table S5 (numerical summary of results), Table S6 (enrichment and marginal *τ\** for all tissues, all traits analysis), Table S15 (enrichment and marginal *τ\** of blood cell types, blood traits analysis), Table S21 (enrichment and marginal *τ\** of brain tissues, brain traits analysis) and Table S27 (joint *τ\** of brain tissues, brain traits analysis). No Supplementary Table is needed for joint *τ\** of all tissues, all traits (1 marginally significant annotation) or blood cell types, blood traits (0 marginally significant annotations)

**Figure 2.**
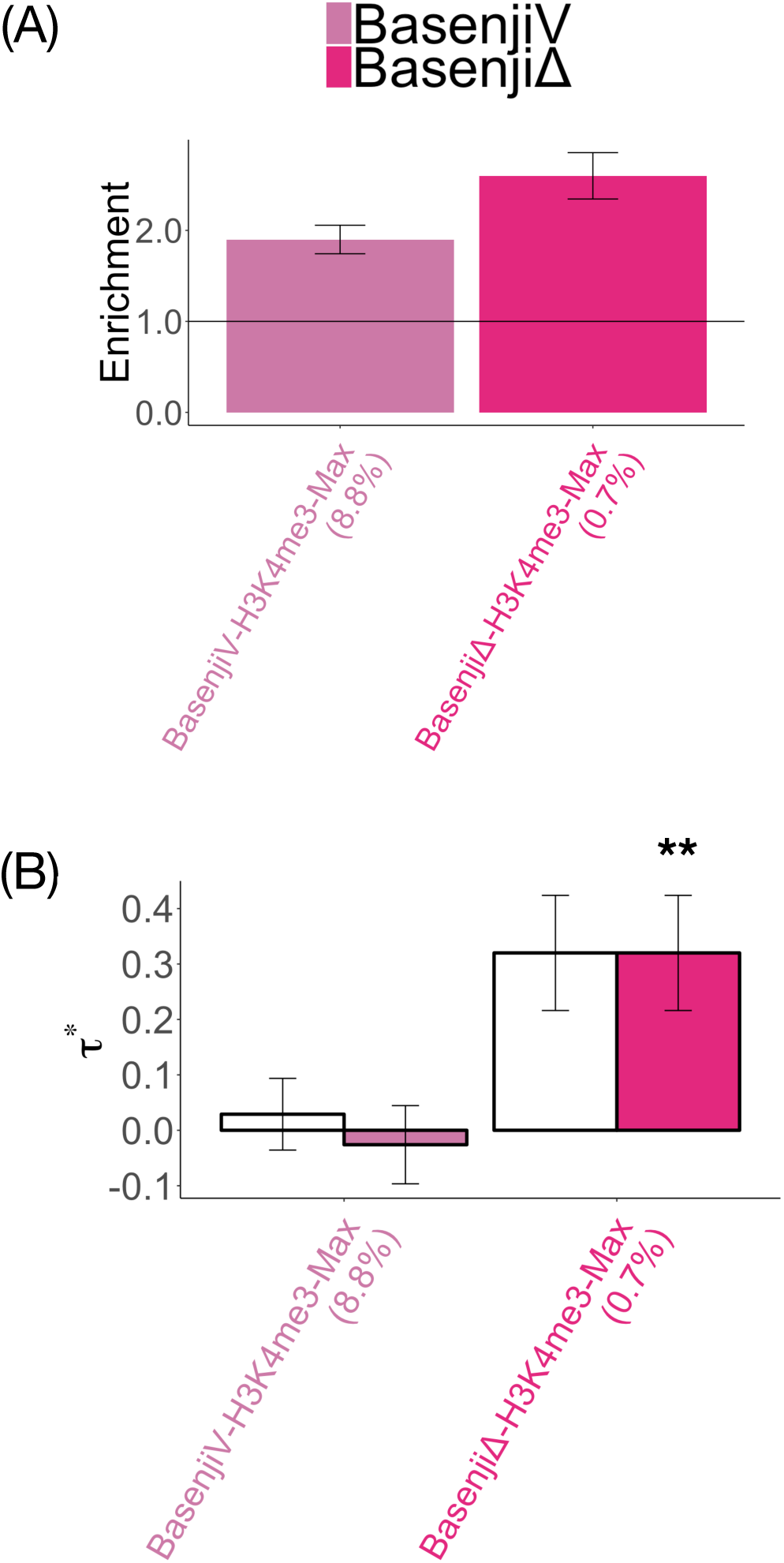
**Disease informativeness of non-tissue-specific allelic-effect deep learning annotations**: (A) Heritability enrichment, conditioned on the non-tissue-specific variant-level joint model. Horizontal line denotes no enrichment. (B) Standardized effect size *τ\** conditioned on either the non-tissue-specific variant-level joint model (marginal analysis: left column, white) or the variant-level joint model plus 1 non-tissue-specific allelic-effect Basenji annotation (BasenjiΔ-H3K4me3-Max) (non-tissue-specific final joint model: right column, dark shading. Results are meta-analyzed across 41 traits. Results are displayed only for the allelic-effect annotation (BasenjiΔ-H3K4me3-Max) with significant *τ\** in marginal analyses after correcting for 106 (variant-level + allelic-effect) non-tissue-specific annotations tested (*P <* 0.05*/*106), along with the corresponding variant-level annotation; the correlation between the two annotations is 0.43. For non-tissue-specific final joint model (right column), ** denotes *P <* 0.05*/*106. Error bars denote 95% confidence intervals. Numerical results are reported in Table S6 and Table S8.

To assess the impact of conditioning on conservation-related annotations, we performed a marginal analysis in which we no longer conditioned on the 11 conservation-related annotations of the baseline-LD model (e.g. GERP++^19, 30^, PhastCons^31^, conservation across 29 mammals^32^, Background selection statistic^33^; Table S9). In this analysis, 6 DeepSEAΔ and 4 BasenjiΔ produced Bonferroni-significant conditional signals (Table S10). This implies that conditioning on conservation-related annotations had a major impact on our primary analysis. Consistent with this finding, we observed substantial correlations (up to *r* = 0.24) between allelic-effect annotations and conservation-related annotations (Figure S7). These results can be viewed as a proof-of-concept that allelic-effect annotations can uncover biological signals.

We investigated the *k*-mer composition of regions proximal to the BasenjiΔ-H3K4me3-Max annotation. For each of all 682 possible *k*-mers with 1≤*k*≤5 (merged with their reverse complements), we assessed the weighted *k*-mer enrichment in 1kb regions around each SNP in the annotation (Methods). Many CpG-related *k*-mers (*k≥*3) attained Bonferroni-significant enrichments, with the largest and most significant enrichments attained by CGCGC (4.1x and P=3.5-e10) and CGGCG (4.1x and P=3.6e-10) (Table S11); these were far larger and more statistically significant than enrichments for simple GC-rich motifs such as the 2-mer CpG (1.2x and P=0.3), ruling out a systematic GC artifact as an explanation for our findings. We note that the CGCG motif is known to correlate with nucleosome occupancy^34, 35^, which may potentially be expected since active promoters tend to have well-positioned nucleosomes marked by H3K4me3. Although the 5-mers CGCGC and CGGCG are too small to associate to known transcription factor binding motifs, we determined that the 9-mer GCGGTGGCT, which was enriched for heritability of blood-related traits in a previous study^36^ and is associated with the ZNF33A transcription factor binding motif, was enriched in the BasenjiΔ-H3K4me3-Max annotation (Table S12).

As an alternative to conditional analysis using S-LDSC, we analyzed various sets of annotations by training a gradient boosting model to classify 12, 296 SNPs from the NIH GWAS catalog^23^ and assessing the AUROC, as in ref.^13, 16^ (Methods); although this is not a formal conditional analysis, comparing the AUROC achieved by different sets of annotations can provide an indication of which annotations provide unique information for disease. Results are reported in Table S13. We reached three main conclusions. First, the aggregated DeepSEAΔ and BasenjiΔ annotations were informative for disease (AUROC = 0.584 and 0.592, respectively, consistent with enrichments of these annotations (DeepSEAΔ: 1.50x, BasenjiΔ: 1.75x) for NIH GWAS SNPs; Table S14). Second, including tissue-specific DeepSEAΔ and BasenjiΔ annotations for all 127 tissues slightly improved the results (AUROC = 0.602 and 0.611, respectively; lower than AUROC = 0.657 and 0.666 reported in ref.^16^ because our analysis was restricted to chromatin marks and did not consider transcription factor binding site (TFBS) or cap analysis of gene expression (CAGE) data). Third, the disease informativeness of the baseline-LD model plus 7 non-tissue-specific annotations from Figure S6) (AUROC = 0.762) was not substantially impacted by adding the aggregated DeepSEAΔ and BasenjiΔ annotations (AUROC = 0.766 and 0.769, respectively). These findings were consistent with our S-LDSC analyses; in particular, the slightly higher AUROC for Basenji and DeepSEA allelic-effect annotations (across all analyses) was consistent with our S-LDSC results showing higher enrichments and a conditionally significant signal for Basenji annotations. Although a key limitation of the NIH GWAS catalog is that it consists predominantly of marginally associated variants that have not been fine-mapped, which thus form a noisy SNP set, these analyses show that it does contain useful signal.

We conclude that allelic-effect DeepSEA and Basenji annotations that were aggregated across tissues were enriched for heritability across the 41 traits (with higher enrichments for Basenji), and that one Basenji allelic-effect annotation was conditionally informative.

### A Basenji brain-specific allelic-effect annotation is conditionally informative for disease

We evaluated the informativeness of blood-specific allelic-effect annotations across 11 blood-related traits (Table S3), and the informativeness of brain-specific allelic-effect annotations across 8 brain-related traits (Table S3).

As in the all-tissues analysis, we first evaluated tissue-specific variant-level annotations. The blood-specific variant-level DeepSEAV and BasenjiV annotations were highly enriched for heritability across 11 blood-related traits, but we determined that none of them were conditionally informative (Figures S8, S9, S10, S11 and Supplementary Note). The brain-specific variant-level DeepSEAV and BasenjiV annotations were also highly enriched for heritability across 8 brain-related traits; surprisingly, two of these annotations (DeepSEAV-H3K4me3-brain-Max and BasenjiV-H3K27ac-brain-Max) were conditionally informative (Figures S12, S13, S14, S15 and Supplementary Note). This is a surprising result, because the brain-specific variant-level deep learning annotations simply predict measured brain-specific variant-level annotations from Roadmap that were also included in the model and suggests new information can be retrieved for brain tissues from de-noising of epigenomic signal using deep learning models. A possible reason for this may be poorer representation of brain tissues in the Roadmap data compared to the blood cell types.

We evaluated the informativeness of 8 blood-specific DeepSEAΔ and 8 blood-specific BasenjiΔ annotations (Table 1) for disease heritability by applying S-LDSC to the 11 blood-related traits. These analyses were conditioned on the the the baseline model plus 7 non-tissue-specific annotations from Figure S6, 6 blood-specific Roadmap and ChromHMM annotations from Figure S11 and BasenjiΔ-H3K4me3-Max (the 1 significant non-tissue-specific allelic-effect annotation; Figure 2 and Table S6).

A summary of the results is provided in Figure 1 (Blood cell types, Blood traits column); numerical results in Table S5. In our marginal analysis of disease heritability, all blood-specific allelic-effect annotations were enriched for disease heritability. Furthermore, blood-specific BasenjiΔ annotations were much more enriched for disease heritability (4.57x) than blood-specific DeepSEAΔ annotations (2.20x), despite similar annotation sizes (Table S15). However, none of the blood-specific allelic-effect annotations attained a Bonferroni-significant standardized effect size (*τ\**) (Table S15). (When we did not condition on the 11 conservation-related annotations of the baseline-LD model (Table S9), this remained the case (Table S16). In contrast, when we did not condition on BasenjiΔ-H3K4me3-Max, 0 blood-specific DeepSEAΔ annotations and 1 BasenjiΔ annotation attained a Bonferroni-significant *τ^*^*(Table S17); when we did not condition on BasenjiΔ-H3K4me3-Max or the 6 blood-specific annotations from S11, 0 blood-specific DeepSEAΔ annotations and 6 blood-specific BasenjiΔ annotations attained a Bonferroni-significant *τ\** (Table S18)).

We also analyzed various sets of blood-specific allelic-effect annotations by training a gradient boosting model to classify 8,741 fine-mapped autoimmune disease SNPs^24^ (relevant to blood-specific annotations only) and assessing the AUROC (analogous to Table S13). Results are reported in Table S19. We reached three main conclusions. First, the aggregated blood-specific DeepSEAΔ and BasenjiΔ annotations were informative for disease, with Basenji being more informative (AUROC = 0.613 and 0.672, respectively, consistent with moderate enrichments (DeepSEAΔ: 1.71x, BasenjiΔ: 2.37x) of these annotations for the fine-mapped SNPs; Table S20). Second, including cell-type-specific allelic-effect DeepSEAΔ and BasenjiΔ annotations for all 27 blood cell types slightly improved the results (AUROC = 0.633 and 0.684, respectively). Third, the disease informativeness of the blood-specific variant-level joint model plus BasenjiΔ-H3K4me3-Max (AUROC = 0.848) was not substantially impacted by adding the aggregated blood-specific DeepSEAΔ and BasenjiΔ annotations (AUROC = 0.847 and 0.851, respectively). These findings were consistent with our S-LDSC analysis.

We evaluated the informativeness of 8 brain-specific DeepSEAΔ and 8 brain-specific BasenjiΔ annotations (Table 1) for disease heritability by applying S-LDSC to the 8 brain-related traits. These analyses were conditioned on the baseline-LD model plus 7 non-tissue-specific annotations from Figure S6, DeepSEAV-H3K4me3-brain-Max and BasenjiV-H3K27ac-brain-Max (the 2 significant brain-specific variant-level annotations; Figure S12) plus 4 additional brain-specific annotations from Figure S15 plus BasenjiΔ-H3K4me3-Max (the 1 significant non-tissue-specific allelic-effect annotation; Figure 2 and Table S6).

A summary of the results is provided in Figure 1 (Brain tissues, Brain traits column); numerical results in Table S5. In our marginal S-LDSC analysis, brain-specific BasenjiΔ annotations were more enriched for disease heritability (2.53x) than brain-specific DeepSEAΔ annotations (1.94x), despite similar annotation sizes. Two brain-specific BasenjiΔ annotations (BasenjiΔ-H3K4me3-brain-Max and BasenjiΔ-H3K4me3-brain-Avg) attained a Bonferroni-significant standardized effect size (*τ\**) (Figure 3 and Table S21). (When we did not condition on the 11 conservation-related annotations of the baseline-LD model (Table S9), 8 brain-specific DeepSEAΔ and 6 brain-specific BasenjiΔ annotations attained a Bonferroni-significant *τ\** (Table S22). In addition, when we did not condition on BasenjiΔ-H3K4me3-Max, 0 brain-specific DeepSEAΔ annotations and 3 brain-specific BasenjiΔ annotations attained a Bonferroni-significant *τ\** (Table S23); when we did not condition on BasenjiΔ-H3K4me3-Max or the 6 brain-specific annotations from Figure S12 and Figure S15, 7 brain-specific DeepSEAΔ annotations and 7 brain-specific BasenjiΔ annotations attained a Bonferroni-significant *τ\** (Table S24)).

**Figure 3.**
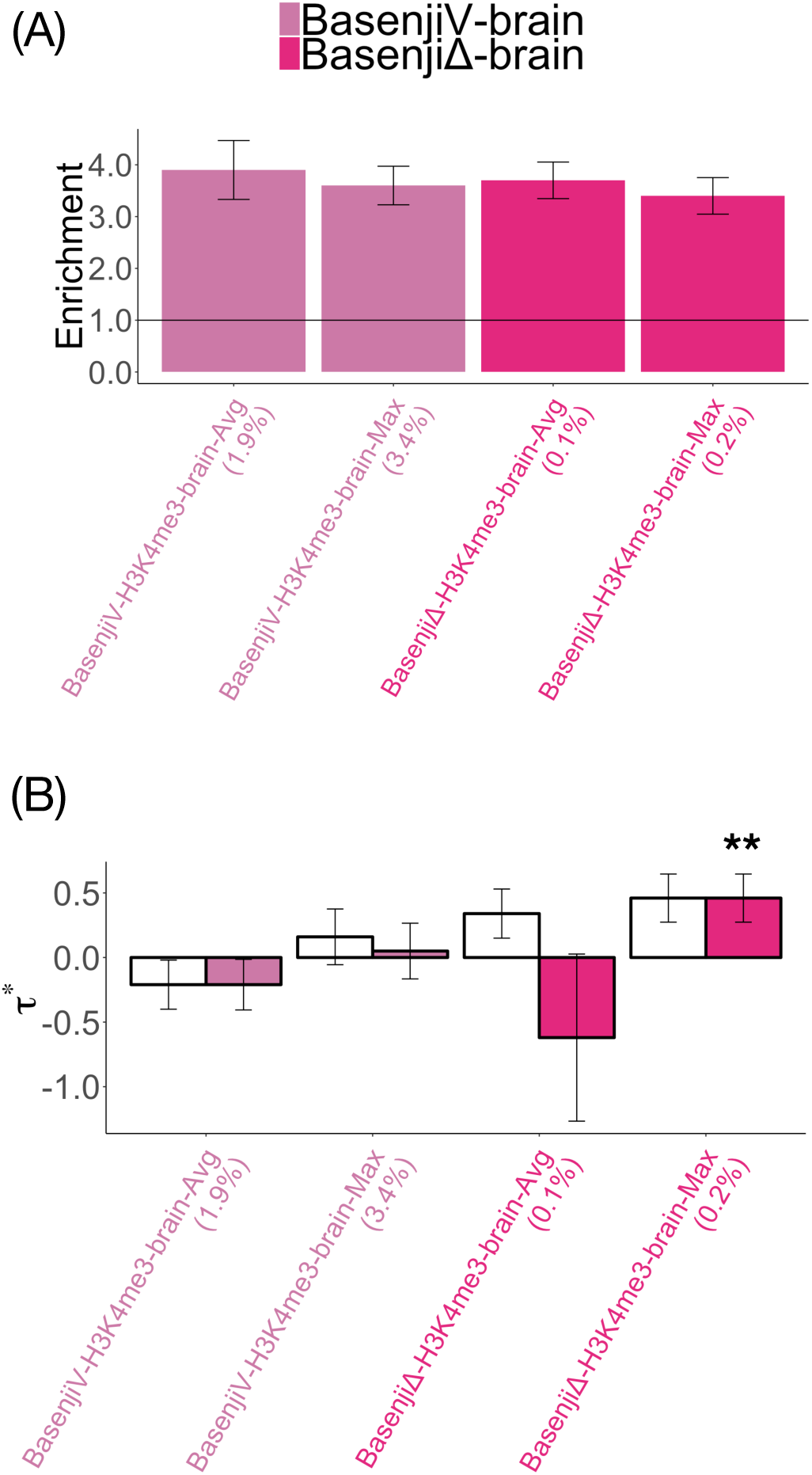
**Disease informativeness of brain-specific allelic-effect deep learning annotations**: (A) Heritability enrichment, conditioned on the brain-specific variant-level joint model and the 1 significant non-tissue-specific allelic-effect annotation (BasenjiΔ-H3K4me3-Max). Horizontal line denotes no enrichment. (B) Standardized effect size *τ\** conditioned on either the brain-specific variant-level joint model and BasenjiΔ-H3K4me3-Max (marginal analysis: left column, white) or the same model plus 1 brain-specific allelic-effect annotation (BasenjiΔ-H3K4me3-brain-Max) (brain-specific final joint model: right column, dark shading). Results are meta-analyzed across 8 brain-related traits. Results are displayed only for the 2 allelic-effect annotations with significant *τ^∗^*in marginal analyses after correcting for 80 (variant-level + allelic-effect) brain-specific annotations tested (*P <* 0.05*/*80), along with the corresponding variant-level annotations; the correlation between the two allelic-effect annotations is 0.78, and the average correlation between the two pairs of variant-level (Basenji) and allelic-effect (BasenjiΔ) annotations is 0.44. For brain-specific final joint model (right column), ** denotes *P <* 0.05*/*80. Error bars denote 95% confidence intervals. Numerical results are reported in Table S21 and Table S27.

Despite the high correlation between variant-level and allelic-effect annotations (*r* = 0.48; Figure S1), the corresponding variant-level annotations (BasenjiV-H3K4me3-brain-Max and BasenjiV-H3K4me3-brain-Avg) did not produce significant signal (Figure 3 and Table S25), consistent with our variant-level analysis (Figure S12). However, when we did not condition on these two variant-level annotations, 4 brain-specific DeepSEAΔ annotations and 6 brain-specific BasenjiΔ annotations attained a Bonferroni-significant *τ\** (Table S26).

We jointly analyzed the two annotations, BasenjiΔ-H3K4me3-brain-Max and BasenjiΔ-H3K4me3-brain-Avg, that were Bonferroni-significant in marginal analyses (Figure 3) by performing forward stepwise elimination to iteratively remove annotations that had conditionally non-significant *τ\** values after Bonferroni correction (based on the 80 variant-level and allelic-effect brain-specific annotations tested in marginal analyses). Of these, only BasenjiΔ-H3K4me3-brain-Max was jointly significant in the resulting brain-specific final joint model, with *τ\** very close to 0.5 (Figure 3, Table S21 and Table S27); annotations with *τ\** 0.5 are unusual, and considered to be important^36^. A *k*-mer enrichment analysis (analogous to above) indicated that BasenjiΔ-H3K4me3-brain-Max was enriched for the *k*-mers CGCGC (6.2x and p=1.1e-25) and CGGCG (6.1x and p=4.9e-25) (far larger and more statistically significant than enrichments for simple GC-rich motifs such as the 2-mer CpG (1.4x and P=0.32)), analogous to BasenjiΔ-H3K4me3-Max (Table S11). The 9-mer GCGGTGGCT (which was enriched for heritability of blood-related traits in a previous study^36^, is associated with the ZNF33A transcription factor binding motif, and was enriched in the BasenjiΔ-H3K4me3-Max annotation; see above) was not enriched in the BasenjiΔ-H3K4me3-brain-Max annotation (Table S12).

We did not consider secondary analyses of fine-mapped SNPs for brain-related traits, due to the lack of a suitable resource analogous to ref.^24^.

We conclude that blood-specific allelic-effect annotations were very highly enriched for heritability but not uniquely informative for blood-related traits, whereas one brain-specific allelic-effect annotation was uniquely informative for brain-related traits. Blood-specific and brain-specific allelic-effect Basenji annotations generally outperformed DeepSEA annotations, yielding higher enrichments and the sole conditionally significant annotation, similar to our non-tissue-specific allelic-effect analyses.

## Discussion

We have evaluated the informativeness for disease of (variant-level and) allelic-effect annotations constructed using two previously trained deep learning models, DeepSEA^13, 15^ and Basenji^16^. We evaluated each annotation’s informativeness using S-LDSC^5, 19^; as a secondary metric, we also evaluated the accuracy of gradient boosting models incorporating deep learning annotations in predicting disease-associated or fine-mapped SNPs^23, 24^, as in previous work^13, 16^. In non-tissue-specific analyses, we identified one allelic-effect Basenji annotation that was uniquely informative for 41 diseases and complex traits. In blood-specific analyses, we identified no deep learning annotations that were uniquely informative for 11 blood-related traits. In brain-specific analyses, we identified brain-specific variant-level DeepSEA and Basenji annotations and a brain-specific allelic-effect Basenji annotation that were uniquely informative for 8 brain-related traits. We caution that—because we conditioned on a broad set of known functional annotations, in contrast to previous studies—the improvements provided by deep learning annotations were very small in magnitude, implying that further work is required to achieve the full potential of deep learning models for complex disease.

Our results imply that the informativeness of deep learning annotations for disease cannot be inferred from metrics such as AUROC that evaluate their accuracy in predicting underlying regulatory annotations derived from experimental assays. Instead, deep learning annotations must be evaluated using methods that specifically assess their informativeness for disease, conditional on a broad set of other functional annotations. The S-LDSC method that we applied here is one such method, and the accuracy of gradient boosting models incorporating both deep learning annotations and other functional annotations can also be a useful metric. We emphasize the importance of conditioning on a broad set of functional annotations, in order to assess whether deep learning models leveraging DNA sequence provide unique (as opposed to redundant) information. Previous work has robustly linked deep learning annotations to disease^12,13,14,15,16^, but those analyses did not condition on a broad set of other functional annotations.

Our work has several limitations, representing important directions for future research. First, our analyses of deep learning annotations using S-LDSC are inherently focused on common variants, but deep learning models have also shown promise in prioritizing rare pathogenic variants^15, 37, 38^. The value of deep learning models for prioritizing rare pathogenic variants has been questioned in a recent analysis focusing on Human Gene Mutation Database (HGMD) variants^39^, meriting further investigation. Second, our analyses of allelic-effect annotations are restricted to unsigned analyses, but signed analyses have also proven valuable in linking deep learning annotations to molecular traits and complex disease^16, 40, 41^. However, genome-wide signed relationships are unlikely to hold for the regulatory marks (DNase and histone marks) that we focus on here, which do not correspond to specific genes or pathways. Third, we focused here on deep learning models trained to predict specific regulatory marks, but deep learning models have also been used to predict a broader set of regulatory features, including gene expression levels and cryptic splicing^15, 16, 38^, that may be informative for complex disease. We have also not considered the application of deep learning models to TFBS, CAGE and ATAC-seq data^16, 41^, which is a promising future research direction. Fourth, we focused here on deep learning models trained using human data, but models trained using data from other species may also be informative for human disease^41, 42^. Fifth, the forward stepwise elimination procedure that we use to identify jointly significant annotations^19^ is a heuristic procedure whose choice of prioritized annotations may be close to arbitrary in the case of highly correlated annotations. Nonetheless, our framework does impose rigorous criteria for conditional informativeness. Finally, beyond deep learning models, it is of high interest to evaluate other machine learning methods for predicting regulatory effects^43,44,45,46,47^.

## URLs

- Annotations analyzed in this study: https://data.broadinstitute.org/alkesgroup/LDSCORE/DeepLearning/
- 1000 Genomes Project Phase 3 data: ftp://ftp.1000genomes.ebi.ac.uk/vol1/ftp/release/20130502
- UK Biobank summary statistics: https://data.broadinstitute.org/alkesgroup/UKBB/
- Baseline-LD annotations: https://data.broadinstitute.org/alkesgroup/LDSCORE/
- BOLT-LMM software: https://data.broadinstitute.org/alkesgroup/BOLT-LMM
- DeepSEA pre-trained model: https://github.com/FunctionLab/ExPecto
- Basenji pre-trained model: https://github.com/calico/basenji

## Acknowledgments

We thank Huwenbo Shi and Steven Gazal for helpful discussions. This research was funded by NIH grants U01 HG009379, R01 MH101244, and R37 MH107649. This research was conducted using the UK Biobank Resource under application 16549.

## Methods

### Genomic annotations and the baseline-LD model

We define a functional annotation as an assignment of a numeric value to each SNP; annotations can be either binary or continuous-valued (Methods). Our focus is on continuous-valued annotations (with values between 0 and 1) trained by deep learning models to predict biological function from DNA sequence. We define a genomic annotation as an assignment of a numeric value to each SNP in a predefined reference panel (e.g., 1000 Genomes Project^25^; see URLs). Continuous-valued annotations can have any real value; our focus is on continuous-valued annotations with values between 0 and 1. Annotations that correspond to known or predicted function are referred to as functional annotations. The baseline-LD model (v.2.1) contains 86 functional annotations (see URLs). These annotations include binary coding, conserved, and regulatory annotations (e.g., promoter, enhancer, histone marks, TFBS) and continuous-valued linkage disequilibrium (LD)-related annotations.

### DeepSEA and Basenji annotations

Tissue-specific deep learning annotations were derived using two pre-trained Convolutional Neural Net (CNN) models: DeepSEA^13, 15^ (architecture from ref.^15^) and Basenji^16^ (see URLs). DeepSEA is a classification based model trained on binary peak call data from 2, 002 cell-type specific TFBS, histone mark and chromatin accessibility annotations from the ENCODE^21^ and Roadmap Epigenomics^11^ projects. Basenji is a Poisson likelihood model trained on original count data from 4, 229 cell-type specific histone mark, chromatin accessibility and FANTOM5 CAGE^48, 49^ annotations. Additionally, Basenji uses dilated convolutional layers that allow scanning much larger contiguous sequence around a variant (≈130kb) compared to DeepSEA (1kb). We restricted our analyses to DNase-I Hypersensitivity Sites (DHS) and 3 histone marks (H3K27ac, H3K4me1 and H3K4me3) that are known to be associated with active enhancers and promoters^50^.

For each SNP with minor allele count ≥ 5 in 1000 Genomes, we applied the pretrained DeepSEA and Basenji models to the surrounding DNA sequence (based on the reference allele) to compute the predicted probability of a tissue-specific chromatin mark (DNase, H3K27ac, H3K4me1, H3K4me3) to generate the corresponding variant-level annotation. To generate the corresponding allelic-effect annotation, we compute the predicted difference in probability between the reference and the alternate alleles. The Basenji annotations were quantile-matched to corresponding DeepSEA annotations to ensure a fair comparison of the two approaches. We aggregated these probabilistic annotations across all 127 Roadmap tissues by taking either the average (Avg) or maximum (Max) to generate non-tissue specific annotations, yielding 8 DeepSEA annotations and 8 Basenji annotations. Similarly, we aggregated over 27 blood cell types (resp. 13 brain tissues) to generate blood (resp. brain) specific annotations for each chromatin mark.

### BiClassCNN annotations

We trained a new deep learning model, BiClassCNN, to prioritize SNPs within non-tissue-specific annotations; analyses of BiClassCNN annotations are described in the Supplementary Note. BiClassCNN analyzes 1kb of human reference sequence around each SNP (analogous to DeepSEA). The positive training set for BiClassCNN consists of 1kb of reference sequence around SNPs that are known to have the functionality of interest (e.g. coding); we included all such sequences in the positive training set. The negative training set consists of 1kb of reference sequence around SNPs that are 1kb away from all SNPs with the functionality of interest; we included a subset of such sequences in the negative training set, so as to match the overall size, GC content and repeat element content of the positive set (as in ref.^43, 51^). We used a shallow Convolutional Neural Net architecture for training (see Figure S16).

We ran two training models, one for the even chromosomes and one for odd chromosomes, and used the trained model on even (resp. odd) chromosomes to assign a predicted probability of functionality (e.g. coding), based on sequence context, to each SNP on odd (resp. even) chromosomes. Unlike DeepSEA and Basenji, BiClassCNN annotations were restricted to regions of known functionality (e.g. coding) by setting annotation values to 0 outside those regions; thus, BiClassCNN prioritizes SNPs within regions of known functionality (e.g. coding). (BiClassCNN annotations that were not restricted in this fashion were far less informative for disease.)

We restricted S-LDSC analyses of BiClassCNN annotations to annotations for which the BiClassCNN AUROC value was at least 0.6 (Table 1 and Table S4). This eliminated three annotations (Intron, H3K27ac and UTR-3’), leaving a total of 12 BiClassCNN annotations.

### Other annotations

We also considered:

- (Table S32) 8 Roadmap annotations^11^ (analogous to DeepSEA and Basenji annotations) imputed using ChromImpute^20^.
- (Table S32) 40 ChromHMM annotations^21, 22^ based on 20 ChromHMM states across 127 Roadmap tissues^11^, again aggregated using the average (Avg) or maximum (Max) across tissues.
- (Table S33) 12 annotations consisting of CpG-island, local CpG-content and local GC-content annotations, as well as these annotations restricted to coding, repressed and TSS regions (for which BiClassCNN produced conditionally significant signals). The CpG-island annotation was retrieved from the UCSC genome browser^52^. Local CpG-content and local GC-content denote the proportion of *CpG* and *G* + *C* dinuclotides in 1kb regions around each variant of the genome, computed using the hg19 reference genome fasta file. By definition, the LocalGCcontent annotation is of larger size than the LocalCpGcontent annotation.
- (Table S33) 3 annotations consisting of a pLI annotation, as well as this annotation restricted to coding and TSS regions. The pLI annotation was defined by annotating each SNP in a 5kb window around a gene with the pLI score of that gene^53^. We did not consider the pLI annotation restricted to repressed regions because unlike TSS and coding, repressed regions are not directly linked to a gene.
- (Table S33) 2 coding annotations, SIFT^54^ and Polyphen^55, 56^, which have been analyzed in previous work^57, 58^.

### Stratified LD score regression

Stratified LD score regression (S-LDSC) is a method that assesses the contribution of a genomic annotation to disease and complex trait heritability^5, 19^. Let *a_cj_* be the value of annotation *c* for SNP *j*, where *a_cj_* may be binary (0/1), continuous or probabilistic. S-LDSC assumes a linear model for *Y* on the normalized genotype matrix *X*:

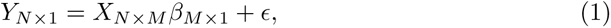

where *β* = (*β*_1_*, β*_2_*,…, β_M_*) is the genotype effect size and *E* denotes environmental noise. S-LDSC assumes that the per-SNP heritability for each SNP *j* can be decomposed as

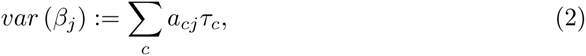

where *τ_c_* is the per-SNP contribution of one unit of annotation *a_c_* to heritability. Under this model assumption, the GWAS summary *χ*^2^ statistics can be linked to *τ_c_* as follows:

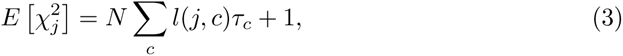

where 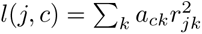 is the *stratified LD score* of SNP *j* with respect to annotation *c* and *r_jk_* is the genotypic correlation between SNPs *j* and *k*.

We assess the informativeness of an annotation *c* using two metrics. The first metric is enrichment (*E*), defined as follows (for binary and probabilistic annotations only):

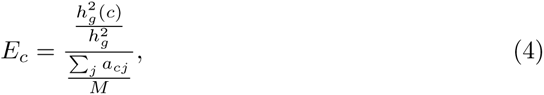

where 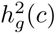 is the heritability explained by the SNPs in annotation *c*, weighted by the annotation values.

The second metric is standardized effect size (*τ\**) defined as follows (for binary, probabilistic, and continuous-valued annotations):

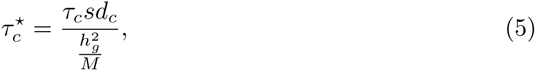

where *sd_c_* is the standard error of annotation *c*, 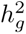 the total SNP heritability and *M* is the total number of SNPs on which this heritability is computed (equal to 5, 961, 159 in our analyses). 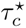 represents the proportionate change in per-SNP heritability associated to a 1 standard deviation increase in the value of the annotation. The main difference between enrichment and *τ\** is that *τ\** quantifies effects that are unique to the focal annotation *c* (after conditioning on all other annotations), whereas enrichment quantifies effects that are unique and/or non-unique to the focal annotation. We computed the statistical significance (p-values) of the enrichment and *τ\** of each annotation via block-jackknife, as previously described^5^; for *τ\**, we assumed that 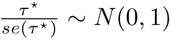.

### Weighted *k*-mer enrichment Analysis

We performed weighted *k*-mer enrichment analyses of the deep learning annotations that were conditionally informative for disease heritability, for all 682 possible *k*-mers with 1 *k* 5 (merged with their reverse complements). Results of these analyses are reported in Table S11 and Table S50.

For each *k*-mer *i*, we computed *k*-mer counts 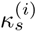 in the 1kb regions around each SNP *s* in the genome.

For each deep learning annotation *D*, for each *k*-mer *i*, we computed the weighted average 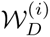 of *k*-mer counts *κ*^(i)^, weighted by values of the probabilistic annotation:

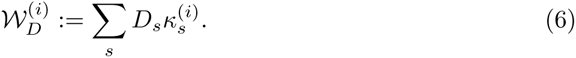

We compared 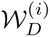 with 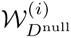, where *D*^null^ is defined as the probabilistic annotation with all values uniformly equal to *D*, the average value (annotation size) of annotation *D*.

We computed the *weighted k-mer enrichment* of annotation *D* with respect to *k*-mer *i* as

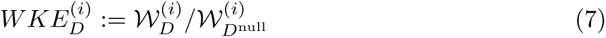

We assessed the statistical significance of the weighted *k*-mer enrichment via a permutation test in which we randomly permuted the values of the deep learning annotation *D* across SNPs and compared 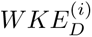 to values of 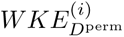 for each permuted annotation *D*^perm^. We computed p-values by fitting a Gaussian distribution to the values of 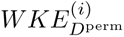 across 10,000 such permutations.

### Classification of disease-associated or fine-mapped SNPs

As an alternative to conditional analysis using S-LDSC, we evaluated the efficacy of various sets of annotations for classifying 12,296 disease-associated SNPs from the NIH GWAS catalog^23^ (as in ref.^13, 16^) or 8,741 fine-mapped autoimmune disease SNPs^24^ against the same number of control SNPs, matched for minor allele frequency. We used XGBoost, a machine learning technique based on gradient tree boosting^59, 60^. To optimize classification performance, we selected XGBoost parameter settings to minimize overfitting, as in ref.^61^.

## Supplementary Note

### Overview of analysis of variant-level annotations

In this supplementary note, we perform disease heritability analysis of 25 variant-level deep learning annotations from three models (8 DeepSEAV, 8 BasenjiV and 9 BiClassCNN), as well as other variant-level epigenomic annotations such as 8 Roadmap annotations^11^ (analogous to DeepSEAV and BasenjiV annotations), imputed using ChromImpute^20^; and 40 ChromHMM annotations^21, 22^ based on 20 ChromHMM states across 127 Roadmap tissues^11^ (Table S2). The variant-level deep learning annotations can be treated as a de-noised representation of Roadmap data learned from sequence architecture. All these annotations contributed to the conditional model in the analysis of the allelic-effect deep learning annotations in the main text. All analyses of variant-level annotations were conditional on a broad set of coding, conserved, regulatory and LD-related annotations in our analyses: 86 annotations from the baseline-LD (v2.1) model^19^. In our analysis of 11 blood-related traits (resp. 8 brain-related traits), for variant-level annotations, we analyzed 16 variant-level annotations (8 DeepSEAV and 8 BasenjiV) that were aggregated across 27 blood cell types (resp. 13 brain tissues), instead of all 127 tissues.

We calculated the variant-level DeepSEAV annotations by applying previously trained DeepSEA models^15^ (see URLs) for each of 4 tissue-specific chromatin marks (DNase, H3K27ac, H3K4me1, H3K4me3) across 127 Roadmap tissues^11^ to 1kb of human reference sequence around each SNP ad calculating the predicted occurrence of the mark based on the sequence (with respect to the reference allele); for each chromatin mark, we aggregated allelic-effect DeepSEA annotations across the 127 tissues using either the average (Avg) or maximum (Max) across tissues (Table 1 and Methods). Similarly, we calculated the variant-level BasenjiV annotations by applying previously trained Basenji models^16^ (see URLs) and aggregating across tissues in analogous fashion (1 and Methods). The constituent tissue-specific BasenjiV annotations do not lie between 0 and 1, and ref.^16^ did not report AUROC values; we transformed these annotations to lie between 0 and 1 via quantile matching with corresponding DeepSEAV annotations, to ensure a fair comparison of the two approaches (Methods). When comparing pairs of annotations that differed only in their aggregation strategy (Avg/Max), chromatin mark (DNase/H3K27ac/H3K4me1/H3K4me3), model (DeepSEA/Basenji) or type (variant-level/allelic-effect), respectively, we observed large correlations across aggregation strategies (average *r* = 0.71), chromatin marks (average *r* = 0.58), models (average *r* = 0.54) and types (average *r* = 0.48) (Figure S1).

Both DeepSEA and Basenji models were trained on tissue-specific features and these features were then aggregated across tissues; as an alternative, we performed a new Convolutional Neural Net training of sequences that separates out non-tissue-specific functional features from putatively non-functional features (Figure S16). We constructed 9 annotations from a new CNN model called BiClassCNN that separates non-tissue-specific functional regions from potentially non-functional regions for different functional categories (Table 1 and Methods); unlike variant-level DeepSEAV and BasenjiV annotations, the BiClassCNN model was only applied to non-tissue-specific annotations (e.g. coding, promoter, or TSS), and was used to prioritize SNPs within these annotations by restricting BiClassCNN predictions to the underlying annotations; prior to this restriction, the BiClassCNN model was highly predictive of the underlying annotations, with AUROC values ranging from 0.67 *−* 0.84 (Table S4).

### No all-tissues variant-level annotation is conditionally informative for disease

In our marginal analysis of disease heritability using S-LDSC, all DeepSEAV annotations were highly enriched for disease heritability (average enrichment 2.2x), but only one DeepSEAV annotation (DeepSEAV-H3K4me3-Max) attained a Bonferroni-significant standardized effect size (*τ\**) conditional on the baseline-LD model (Figure S2 and Table S29). This implies that high AUROC values for variant-level annotations do not necessarily translate into conditional informativeness for human disease. A summary of the results is provided in Figure S17 (All tissues, All traits column; numerical results in Table S28), which reports the number of variant-level annotations of various types with significant heritability enrichment, marginal conditional signal, and joint conditional signal, respectively. In our marginal analysis of disease heritability using S-LDSC, all BasenjiV annotations were highly enriched for disease heritability (average enrichment 2.2x), but no BasenjiV annotation attained a Bonferroni-significant *τ\** conditional on the baseline-LD model (Table S29). In our marginal analysis of disease heritability using S-LDSC, all BiClassCNN annotations were highly enriched for disease heritability (average enrichment 2.9x) except for BiClassCNN-Repressed (which was depleted, 0.68x), and 4 BiClassCNN annotations attained a Bonferroni-significant *τ\** conditional on the baseline-LD model (Figure S2 and Table S30). 3 of these 4 annotations (BiClassCNN-Coding, BiClassCNN-Repressed, and BiClassCNN-TSS) produced independent signals (Table S31).

To ensure a conservative assessment of which annotations provide unique information for disease, we included a broader set of annotations in our analyses. First, we included 8 Roadmap annotations^11^ (analogous to DeepSEAV and BasenjiV annotations) imputed using ChromImpute^20^, and 40 ChromHMM annotations^21, 22^ based on 20 ChromHMM states across 127 Roadmap tissues^11^, again aggregated using the average (Avg) or maximum (Max) across tissues (Table S2 and Methods). Of these, 4 Roadmap annotations and 17 ChromHMM annotations attained a Bonferroni-significant *τ\** conditional on the baseline-LD model (Figure S3, Figure S4 and Table S32). Second, motivated by the BiClassCNN results (see above), we investigated additional annotations related to coding, repressed and TSS regions. We determined that BiClassCNN-TSS was highly correlated with CpG-island, local CpG-content (±1kb) and local GC-content (±1kb) annotations (Figure S5). We thus incorporated CpG-island, local CpG-content and local GC-content annotations, as well as these annotations restricted to coding, repressed and TSS regions. We also considered a gene-level annotation (±5kb) based on probability of loss-of-function intolerance (pLI)^53^, the pLI annotation restricted to coding and TSS regions, and PolyPhen^55, 56^ and SIFT^54^ coding annotations (12 + 5 = 17 additional annotations; Table S33 and Methods). 11 of these 17 annotations attained a Bonferroni-significant *τ^*^*conditional on the baseline-LD model (Table S33).

We jointly analyzed all 38 annotations that were Bonferroni-significant in our marginal analyses (Figure S2, Figure S3, Figure S4, Table S29, Table S30, Table S32, Table S33) by performing forward stepwise elimination to iteratively remove annotations that had conditionally non-significant *τ^∗^*values after Bonferroni correction (based on a total of 106 (variant-level + allelic-effect) non-tissue-specific annotations tested in marginal analyses). Of these, 0 DeepSEAV, 0 BiClassCNN, 1 Roadmap, 1 ChromHMM, 3 local GC-content and 2 pLI annotations were jointly significant in the resulting joint model (Figure S2, Figure S6 and Table S34); these annotations were included in all of the conditional analyses below. We determined that the surviving Roadmap-H3K4me1-Avg annotation was sufficient to eliminate the DeepSEAV-H3K4me3-Max annotation that was significant in our marginal analysis (Table S35), and that the local GC-content annotations were sufficient to eliminate the 4 BiClassCNN annotations that were significant in our marginal analysis (Table S36).

We also analyzed various sets of allelic-effect annotations by training a gradient boosting model to classify 12, 296 SNPs from the NIH GWAS catalog^23^ and assessing the AUROC (analogous to Table S13), as in ref.^13, 16^ (Methods). Results are reported in Table S37. We reached three main conclusions. First, the aggregated variant-level DeepSEAV and BasenjiV annotations were only moderately informative for disease (AUROC = 0.582 and 0.612, respectively, consistent with moderate enrichments (DeepSEAV: 1.49x, BasenjiV: 1.49x) of these annotations for NIH GWAS SNPs; Table S38). Second, including tissue-specific variant-level DeepSEAV and BasenjiV annotations for all 127 tissues had little impact on the results (AUROC = 0.591 and 0.620, respectively). Third, the disease informativeness of the baseline-LD model (AUROC = 0.758) was not substantially impacted by adding the aggregated variant-level DeepSEAV and BasenjiV annotations (AUROC = 0.759 and 0.766, respectively); results were similar for the non-tissue-specific joint model (baseline-LD model + 7 annotations). These findings were consistent with our S-LDSC analyses.

We conclude that variant-level DeepSEAV and BasenjiV annotations that were aggregated across tissues were highly enriched for heritability but not conditionally informative across the 41 traits, and that marginally significantly informative BiClassCNN annotations were explained away by local GC-content.

### DeepSEA and Basenji brain-specific variant-level annotations are conditionally informative for disease

We next evaluated the informativeness of blood-specific variant-level annotations for blood-related diseases and traits. We analyzed 11 independent blood-related traits (6 autoimmune diseases and 5 blood cell traits; ref.^28^ and Table S3) by running S-LDSC conditioned on the non-tissue-specific variant-level joint model (baseline-LD model + 7 annotations; Figure S6).

We analyzed 8 blood-specific variant-level DeepSEAV and 8 blood-specific variant-level BasenjiV annotations (Table S39), representing the blood-specific analogues of the non-tissue-specific variant-level DeepSEAV and BasenjiV annotations from Table 1; in each case we computed the Average (Avg) or Maximum (Max) of cell-type-specific DeepSEAV and BasenjiV variant-level annotations across blood cell types. A summary of the results is provided in Figure S17 (Blood cell types, Blood traits column); numerical results in Table S28. In our marginal S-LDSC analysis, both blood-specific BasenjiV annotations and blood-specific DeepSEAV annotations were very highly enriched (average enrichment 4.15x and 3.75x, respectively). However, only 0 blood-specific DeepSEAV annotations and 4 blood-specific BasenjiV annotations attained a Bonferroni-significant standardized effect size (*τ\**) conditional on the non-tissue-specific variant-level joint model (Figure S8 and Table S39).

We also analyzed 8 blood-specific Roadmap and 40 blood-specific ChromHMM annotations (Table S40), representing the blood-specific analogues of the non-tissue-specific Roadmap and ChromHMM annotations from Table S2. In our marginal S-LDSC analysis, all 8 blood-specific Roadmap annotations and 23 blood-specific ChromHMM annotations attained a Bonferroni-significant *τ\** (Figure S9, Figure S10 and Table S40).

We jointly analyzed the 35 blood-specific annotations that were Bonferroni-significant in marginal analyses (4 BasenjiV, 8 Roadmap, 23 ChromHMM; Figure S8, Figure S9 and Figure S10) by performing forward stepwise elimination to iteratively remove annotations that had conditionally non-significant *τ\** values after Bonferroni correction (based on the 80 (variant-level + allelic effect) blood-specific annotations tested in marginal analyses). Of these, 0 BasenjiV, 2 Roadmap and 4 ChromHMM annotations were jointly significant in the resulting joint model (Figure S8, Figure S11 and Table S41). We determined that the 2 surviving blood-specific Roadmap annotations were sufficient to eliminate 3 of the 4 blood-specific BasenjiV annotations that were significant in marginal analyses (Table S42). We note that several of the blood-specific Roadmap and ChromHMM annotations had *τ* >* 0.5 in the blood-specific joint model (Figure S11) (annotations with *τ* >* 0.5 are unusual, and considered to be important^36^), consistent with the well-documented importance of tissue-specific annotations, particularly for blood-related traits^5, 11^.

We also analyzed various sets of blood-specific annotations by training a gradient boosting model to classify 8,741 fine-mapped autoimmune disease SNPs^24^ (relevant to blood-specific annotations only) and assessing the AUROC (analogous to Table S37). Results are reported in Table S43. We reached three main conclusions. First, the aggregated blood-specific variant-level DeepSEAV and BasenjiV annotations were informative for disease, with BasenjiV being more informative (AUROC = 0.621 and 0.664, respectively, consistent with moderate enrichments (DeepSEAV: 1.75x, BasenjiV: 1.97x) of these annotations for the fine-mapped SNPs; Table S44). Second, including cell-type-specific variant-level DeepSEAV and BasenjiV annotations for all 27 blood cell types had little impact on the results (AUROC = 0.631 and 0.671, respectively). Third, the disease informativeness of the non-tissue-specific joint model (AUROC = 0.845) was not substantially impacted by adding aggregated blood-specific DeepSEAV and BasenjiV variant-level annotations (AUROC = 0.849 and 0.853, respectively); results were similar for the the blood-specific joint model (non-tissue-specific joint model + 6 blood annotations). These findings were consistent with our S-LDSC analyses.

We next evaluated the informativeness of brain-specific variant-level annotations for brain-related diseases and traits, analogous to blood. We analyzed 8 independent brain-related traits (ref.^28^ and Table S3) by running S-LDSC conditioned on the non-tissue-specific variant-level joint model (baseline-LD model + 7 annotations; Figure S6).

We analyzed 8 brain-specific variant-level DeepSEAV and 8 brain-specific variant-level BasenjiV annotations (Table S45), representing the brain-specific analogues of the non-tissue-specific variant-level DeepSEAV and BasenjiV annotations from Table 1; in each case we computed the Average (Avg) or Maximum (Max) of tissue-type-specific DeepSEAV and BasenjiV variant-level annotations across brain tissues. A summary of the results is provided in Figure S17 (Brain tissues, Brain traits column); numerical results in Table S28. In our marginal S-LDSC analysis, both brain-specific DeepSEAV and BasenjiV annotations were highly enriched (average enrichment 2.71x and 2.73x, respectively; Table S45). Furthermore, all 8 brain-specific DeepSEAV annotations and all 8 brain-specific BasenjiV annotations attained a Bonferroni-significant standardized effect size (*τ\**) conditional on the non-tissue-specific variant-level joint model (Figure S12 and Table S45).

We also analyzed 8 brain-specific Roadmap and 40 brain-specific ChromHMM annotations (Table S46), representing the brain-specific analogues of the non-tissue-specific Roadmap and ChromHMM annotations from Table S2. In our marginal S-LDSC analysis, all 8 brain-specific Roadmap annotations and 13 brain-specific ChromHMM annotations attained a Bonferroni-significant *τ^∗^*(Figure S13, Figure S14 and Table S46).

We jointly analyzed the 37 brain-specific annotations that were Bonferroni-significant in marginal analyses (8 DeepSEAV, 8 BasenjiV, 8 Roadmap, 13 ChromHMM; Figure S12, Figure S13 and Figure S14) by performing forward stepwise elimination to iteratively remove annotations that had conditionally non-significant *τ\** values after Bonferroni correction (based on the 80 (variant-level + allelic-effect) brain-specific annotations tested in marginal analyses). Of these, 1 DeepSEAV, 1 BasenjiV, 1 Roadmap and 3 ChromHMM annotations were jointly significant in the resulting joint model (Figure S12, Figure S15 and Table S47). We determined that neither the 1 surviving brain-specific Roadmap annotation nor the 3 surviving brain-specific ChromHMM annotations were sufficient to eliminate any of the 8 DeepSEAV and 8 BasenjiV brain annotations that were significant in marginal analyses (Table S48 and Table S49). We note that none of the brain-specific DeepSEAV, BasenjiV, Roadmap and ChromHMM annotations had *τ* >* 0.5 in the brain-specific joint model (Figure S12 and Figure S15). We did not consider secondary analyses of fine-mapped SNPs for brain-related traits, due to the lack of a suitable resource analogous to ref.^24^.

We conclude that blood-specific variant-level DeepSEAV and BasenjiV annotations that were aggregated across blood cell types were very highly enriched for heritability but not uniquely informative for blood-related traits. On the other hand, brain-specific variant-level DeepSEAV and BasenjiV annotations that were aggregated across brain tissues were uniquely informative for brain-related traits. This may be because brain tissues are not as well-represented in Roadmap data as blood cell types, leaving more room for new information to be retrieved by deep learning models. This also justifies that though allelic-effect annotations should definitely be the annotations of primary interest, variant-level annotations may also be of some value, and should be conditioned on.

---

**Table S1.** **List of non-tissue-specific variant-level deep learning annotations analyzed.** We list the 25 variant-level deep learning annotations (8 DeepSEAV, 8 BasenjiV, 9 BiClassCNN) and their annotation sizes.

| Variant-level annotations | Size (%) |
| --- | --- |
| DeepSEAV-DNase-Avg | 1.6 |
| DeepSEAV-DNase-Max | 13.2 |
| DeepSEAV-H3K27ac-Avg | 3.2 |
| DeepSEAV-H3K27ac-Max | 13.7 |
| DeepSEAV-H3K4me1-Avg | 5.5 |
| DeepSEAV-H3K4me1-Max | 23.2 |
| DeepSEAV-H3K4me3-Avg | 1.7 |
| DeepSEAV-H3K4me3-Max | 8.8 |
| BasenjiV-DNase-Avg | 1.6 |
| BasenjiV-DNase-Max | 13.1 |
| BasenjiV-H3K27ac-Avg | 3.2 |
| BasenjiV-H3K27ac-Max | 13.7 |
| BasenjiV-H3K4me1-Avg | 5.5 |
| BasenjiV-H3K4me1-Max | 23.2 |
| BasenjiV-H3K4me3-Avg | 1.7 |
| BasenjiV-H3K4me3-Max | 8.8 |
| BiClassCNN-Coding | 1.0 |
| BiClassCNN-Enhancer | 2.6 |
| BiClassCNN-Promoter | 2.7 |
| BiClassCNN-Repressed | 36.0 |
| BiClassCNN-SuperEnhancer | 6.7 |
| BiClassCNN-TFBS | 7.9 |
| BiClassCNN-TSS | 0.8 |
| BiClassCNN-UTR-5' | 0.4 |
| BiClassCNN-WeakEnhancer | 1.1 |

**Table S2.** **List of non-tissue-specific Roadmap and ChromHMM annotations analyzed**. We list the 8 Roadmap and 40 ChromHMM annotations that we analyzed, together with the size of the annotation. These annotations are probabilistic, thus the size of the annotation is defined as the average annotation across all reference SNPs.

| Annotation | Size (in %) |
| --- | --- |
| Roadmap-DNase-Avg | 2.2% |
| Roadmap-H3K27ac-Avg | 3.1% |
| Roadmap-H3K4me1-Avg | 4.4% |
| Roadmap-H3K4me3-Avg | 1.7% |
| Roadmap-DNase-Max | 13.4% |
| Roadmap-H3K27ac-Max | 21.4% |
| Roadmap-H3K4me1-Max | 25.4% |
| Roadmap-H3K4me3-Max | 6.6% |
| ChromHMM-DNase-Avg | 0.7% |
| ChromHMM-Active_Enhancer_1-Avg | 0.3% |
| ChromHMM-Active_Enhancer_2-Avg | 0.4% |
| ChromHMM-Enhancer_acetylation-Avg | 0.3% |
| ChromHMM-Active_Enhancer_Flanking-Avg | 0.5% |
| ChromHMM-Weak_Enhancer_1-Avg | 0.3% |
| ChromHMM-Weak_Enhancer_2-Avg | 1.2% |
| ChromHMM-Heterochromatin-Avg | 1.4% |
| ChromHMM-Promoter_Bivalent-Avg | 0.3% |
| ChromHMM-Promoter_Downstream_1-Avg | 0.4% |
| ChromHMM-Promoter_Downstream_2-Avg | 0.2% |
| ChromHMM-Promoter_Poised-Avg | 0.2% |
| ChromHMM-Promoter_Upstream-Avg | 0.4% |
| ChromHMM-Quiescent-Avg | 78.4% |
| ChromHMM-Repressed_Polycomb-Avg | 1.7% |
| ChromHMM-TSS-Avg | 0.2% |
| ChromHMM-Transcription_Enhancer_3'-Avg | 0.2% |
| ChromHMM-Transcription_Enhancer_5'-Avg | 0.4% |
| ChromHMM-Transcription_Weak_Enhancer-Avg | 0.5% |
| ChromHMM-Transcriptional_Regulator-Avg | 0.3% |
| ChromHMM-DNase-Max | 9.2% |
| ChromHMM-Active_Enhancer_1-Max | 7.1% |
| ChromHMM-Active_Enhancer_2-Max | 9.9% |
| ChromHMM-Enhancer_acetylation-Max | 11.3% |
| ChromHMM-Active_Enhancer_Flanking-Max | 11.1% |
| ChromHMM-Weak_Enhancer_1-Max | 7.7% |
| ChromHMM-Weak_Enhancer_2-Max | 19.9% |
| ChromHMM-Heterochromatin-Max | 7.1% |
| ChromHMM-Promoter_Bivalent-Max | 1.9% |
| ChromHMM-Promoter_Downstream_1-Max | 1.4% |
| ChromHMM-Promoter_Downstream_2-Max | 1.5% |
| ChromHMM-Promoter_Poised-Max | 6.7% |
| ChromHMM-Promoter_Upstream-Max | 2.5% |
| ChromHMM-Quiescent-Max | 91.0% |
| ChromHMM-Repressed_Polycomb-Max | 9.9% |
| ChromHMM-TSS-Max | 0.9% |
| ChromHMM-Transcription_Enhancer_3'-Max | 3.1% |
| ChromHMM-Transcription_Enhancer_5'-Max | 4.8% |
| ChromHMM-Transcription_Weak_Enhancer-Max | 4.7% |
| ChromHMM-Transcriptional_Regulator-Max | 3.3% |

**Table S3.** **List of 41 diseases and complex traits analyzed**. We first list the 11 blood-related traits (5 blood cell traits and 6 autoimmune diseases), followed by 10 brain traits (with 2 traits analyzed by two different datasets). Overall, for 6 traits we analyzed two different data sets, leading to a total of 47 data sets.

| Trait | Source | N |
| --- | --- | --- |
| Platelet Count | UKBiobank<sup>29</sup> | 444382 |
| Red Blood Cell Count | UKBiobank<sup>29</sup> | 445174 |
| Red Blood Cell Distribution Width | UKBiobank<sup>29</sup> | 442700 |
| Eosinophil Count | UKBiobank<sup>29</sup> | 439938 |
| White Blood Cell Count | UKBiobank<sup>29</sup> | 444502 |
| Auto Immune Traits (Sure) | UKBiobank<sup>29</sup> | 459324 |
| Crohn's Disease | Jostins et al., 2012 Nature<sup>62</sup> | 20883 |
| Rheumatoid Arthritis | Okada et al., 2014 Nature<sup>63</sup> | 37681 |
| Ulcerative Colitis | Jostins et al., 2012 Nature<sup>62</sup> | 27432 |
| Lupus | Bentham et al., 2015<sup>64</sup> | 14267 |
| Celiac | Dubois et al., 2010<sup>65</sup> | 15283 |
| Age at Menarche | UKBiobank<sup>29</sup> | 242278 |
| BMI | Speliotes et al., 2010 Nat Genet<sup>66</sup> | 122033 |
| BMI | UKBiobank<sup>29</sup> | 457824 |
| Depressive symptoms | Okbay et al., 2016 Nat Genet<sup>67</sup> | 161460 |
| Neuroticism | UKBiobank<sup>29</sup> | 372066 |
| Schizophrenia | SCZ Working Group of the PGC, 2014 Nature<sup>4</sup> | 70100 |
| Years of Education | Okbay et al., 2016 Nature<sup>67</sup> | 328917 |
| Ever Smoked | TAG Consortium, 2010 Nat Genet<sup>68</sup> | 74035 |
| Smoking Status | UKBiobank<sup>29</sup> | 457683 |
| Bipolar Disorder | PGC Bipolar Disorder Group<sup>69</sup> | 16731 |
| Age at Menopause | UKBiobank<sup>29</sup> | 143025 |
| Age first birth | Barban et al., 2016 Nat Genet<sup>70</sup> | 222037 |
| Anorexia | Boraska et al., 2014 Mol Psych<sup>71</sup> | 32143 |
| Autism Spectrum | PGC Cross-Disorder Group, 2013 Lancet<sup>72</sup> | 10263 |
| College Education | UKBiobank<sup>29</sup> | 454813 |
| Coronary Artery Disease | Schunkert et al., 2011 Nat Genet<sup>73</sup> | 77210 |
| Dermatologic Diseases | UKBiobank<sup>29</sup> | 459324 |
| Eczema | UKBiobank<sup>29</sup> | 458699 |
| FEV1-FVC Ratio | UKBiobank<sup>29</sup> | 371949 |
| Forced Vital Capacity (FVC) | UKBiobank<sup>29</sup> | 371949 |
| Hair Color | UKBiobank<sup>29</sup> | 452720 |
| HDL | Teslovich et al., 2010 Nature<sup>74</sup> | 97749 |
| Heel T Score | UKBiobank<sup>29</sup> | 445921 |
| Height | Lango Allen et al., 2010 Nature<sup>75</sup> | 131547 |
| Height | UKBiobank<sup>29</sup> | 458303 |
| High Cholesterol | UKBiobank<sup>29</sup> | 459324 |
| Hypothyroidism | UKBiobank<sup>29</sup> | 459324 |
| LDL | Teslovich et al., 2010 Nature<sup>74</sup> | 93354 |
| Morning Person | UKBiobank<sup>29</sup> | 410520 |
| Number children ever born | Barban et al., 2016 Nat Genet<sup>70</sup> | 318863 |
| Respiratory and Ear-nose-throat Diseases | UKBiobank<sup>29</sup> | 459324 |
| Skin Color | UKBiobank<sup>29</sup> | 453609 |
| Sunburn Occasion | UKBiobank<sup>29</sup> | 344229 |
| Tanning | UKBiobank<sup>29</sup> | 449984 |
| Type 2 Diabetes | Morris et al., 2012 Nat Genet<sup>76</sup> | 60786 |
| Type 2 Diabetes | UKBiobank<sup>29</sup> | 459324 |

**Table S4.** **AUROC values of deep learning annotations**. AUROC (Area under the ROC curve) for DeepSEAV (reported by ref. ^15^) and BiClassCNN annotations. For the DeepSEA model, the Minimum-Maximum (Average) AUROC across all tissues is reported. The constituent tissue-specific Basenji annotations do not lie on a probabilistic scale, and ref.^16^ did not report AUROC values

| Annotation | Area under ROC curve (AUROC) |
| --- | --- |
| DeepSEAV-DNase | All: 0.83-0.95 (0.91)<br>brain: 0.84-0.92 (0.88)<br>blood: 0.86-0.94 (0.89) |
| DeepSEAV-H3K27ac | All: 0.78-0.88 (0.84)<br>brain: 0.80-0.85 (0.82)<br>blood: 0.78-0.87 (0.83) |
| DeepSEAV-H3K4me1 | All: 0.77-0.86 (0.81)<br>brain: 0.79-0.83 (0.80)<br>blood: 0.78-0.83 (0.81) |
| DeepSEAV-H3K4me3 | All: 0.88-0.97 (0.92)<br>brain: 0.89-0.95 (0.91)<br>blood: 0.88-0.95 (0.92) |
| BiClassCNN-Coding | 0.79 |
| BiClassCNN-DNase | 0.69 |
| BiClassCNN-Enhancer | 0.72 |
| BiClassCNN-H3K4me1 | 0.70 |
| BiClassCNN-H3K4me3 | 0.73 |
| BiClassCNN-Promoter | 0.76 |
| BiClassCNN-Repressed | 0.70 |
| BiClassCNN-SuperEnhancer | 0.67 |
| BiClassCNN-TFBS | 0.75 |
| BiClassCNN-TSS | 0.84 |
| BiClassCNN-UTR-5' | 0.75 |
| BiClassCNN-WeakEnhancer | 0.74 |

**Table S5.** **Number of significantly disease informative non-tissue-specific allelic-effect deep learning annotations**. Number of significantly disease informative non-tissue-specific allelic-effect annotations across different aggregation strategies (A) different deep learning models (DeepSEA and Basenji), (B) different aggregation strategies (Average and Maximum) and (C) different types of epigenomic marks, in terms of marginal enrichment in heritability, conditional *τ\** and joint *τ\** for all traits, blood traits and brain traits respectively. See Figure 1 for visual illustration.

| **PANEL A** |  |  |
| --- | --- | --- |
| Enrichment |  |  |
| Model | Aggr. Strategy | Number of significant annotations |
| DeepSEAΔ | All tissues. All traits | 8 |
|  | Brain tissues, Brain traits | 8 |
|  | Blood cell types, Blood traits | 8 |
| BasenjiΔ | All tissues. All traits | 8 |
|  | Brain tissues, Brain traits | 8 |
|  | Blood cell types, Blood traits | 8 |
| Marginal $\tau^*$ | | |
| Model | Aggregation Strategy | Number of significant annotations |
| DeepSEAΔ | All tissues. All traits | 0 |
|  | Brain tissues, Brain traits | 0 |
|  | Blood cell types, Blood traits | 0 |
| BasenjiΔ | All tissues. All traits | 1 |
|  | Brain tissues, Brain traits | 2 |
|  | Blood cell types, Blood traits | 0 |
| Joint $\tau^*$ | | |
| Model | Aggregation Strategy | Number of significant annotations |
| DeepSEAΔ | All tissues. All traits | 0 |
|  | Brain tissues, Brain traits | 0 |
|  | Blood cell types, Blood traits | 0 |
| BasenjiΔ | All tissues. All traits | 1 |
|  | Brain tissues, Brain traits | 1 |
|  | Blood cell types, Blood traits | 0 |
| **PANEL B** |  |  |
| Enrichment |  |  |
| Model | Aggr. Strategy | Number of significant annotations |
| Avg | All tissues. All traits | 8 |
|  | Brain tissues, Brain traits | 8 |
|  | Blood cell types, Blood traits | 8 |
| Max | All tissues. All traits | 8 |
|  | Brain tissues, Brain traits | 8 |
|  | Blood cell types, Blood traits | 8 |
| Marginal $\tau^*$ | | |
| Model | Aggregation Strategy | Number of significant annotations |
| Avg | All tissues. All traits | 0 |
|  | Brain tissues, Brain traits | 1 |
|  | Blood cell types, Blood traits | 0 |
| Max | All tissues. All traits | 1 |
|  | Brain tissues, Brain traits | 1 |
|  | Blood cell types, Blood traits | 0 |
| Joint $\tau^*$ | | |

| Model | Aggregation Strategy | Number of significant annotations |
| --- | --- | --- |
| Avg | All tissues. All traits | 0 |
|  | Brain tissues, Brain traits | 0 |
|  | Blood cell types, Blood traits | 0 |
| Max | All tissues. All traits | 1 |
|  | Brain tissues, Brain traits | 1 |
|  | Blood cell types, Blood traits | 0 |

**PANEL C** Enrichment
| Model | Aggr. Strategy | Number of significant annotations |
| --- | --- | --- |
| DNase | All tissues. All traits | 4 |
|  | Brain tissues, Brain traits | 4 |
|  | Blood cell types, Blood traits | 4 |
| H3K27ac | All tissues. All traits | 4 |
|  | Brain tissues, Brain traits | 4 |
|  | Blood cell types, Blood traits | 4 |
| H3K4me1 | All tissues. All traits | 4 |
|  | Brain tissues, Brain traits | 4 |
|  | Blood cell types, Blood traits | 4 |
| H3K4me3 | All tissues. All traits | 4 |
|  | Brain tissues, Brain traits | 4 |
|  | Blood cell types, Blood traits | 4 |

Marginal $\tau^*$
| Model | Aggregation Strategy | Number of significant annotations |
| --- | --- | --- |
| DNase | All tissues. All traits | 0 |
|  | Brain tissues, Brain traits | 0 |
|  | Blood cell types, Blood traits | 0 |
| H3K27ac | All tissues. All traits | 0 |
|  | Brain tissues, Brain traits | 0 |
|  | Blood cell types, Blood traits | 0 |
| H3K4me1 | All tissues. All traits | 0 |
|  | Brain tissues, Brain traits | 0 |
|  | Blood cell types, Blood traits | 0 |
| H3K4me3 | All tissues. All traits | 1 |
|  | Brain tissues, Brain traits | 2 |
|  | Blood cell types, Blood traits | 0 |

Joint $\tau^*$
| Model | Aggregation Strategy | Number of significant annotations |
| --- | --- | --- |
| DNase | All tissues. All traits | 0 |
|  | Brain tissues, Brain traits | 0 |
|  | Blood cell types, Blood traits | 0 |
| H3K27ac | All tissues. All traits | 0 |
|  | Brain tissues, Brain traits | 0 |
|  | Blood cell types, Blood traits | 0 |
| H3K4me1 | All tissues. All traits | 0 |
|  | Brain tissues, Brain traits | 0 |
|  | Blood cell types, Blood traits | 0 |
| H3K4me3 | All tissues. All traits | 1 |
|  | Brain tissues, Brain traits | 1 |
|  | Blood cell types, Blood traits | 0 |

**Table S6.** **S-LDSC results for marginal analysis of non-tissue-specific allelic-effect DeepSEA**Δ **and Basenji**Δ **annotations conditioned on non-tissue-specific variant-level joint model**. Standardized Effect sizes (*τ\**) and Enrichment (E) of 8 non-tissue-specific DeepSEAΔ and 8 non-tissue-specific BasenjiΔ annotations, when conditioned on the non-tissue-specific variant-level joint model (baseline-LD model + 7 annotations from Figure S6). Results are meta-analyzed across 41 traits.

| Annotation | $\tau^*$ | se($\tau^*$) | p($\tau^*$) | $E$ | se ($E$) | p($E$) |
| --- | --- | --- | --- | --- | --- | --- |
| DeepSEAΔ-DNase-Avg (0.3%) | 0.07 | 0.07 | 0.32 | 2.3 | 0.21 | 3.3e-07 |
| DeepSEAΔ-DNase-Max (2.0%) | -0.034 | 0.066 | 0.61 | 1.6 | 0.1 | 1.5e-05 |
| DeepSEAΔ-H3K27ac-Avg (0.2%) | 0.056 | 0.075 | 0.45 | 2 | 0.12 | 9.7e-10 |
| DeepSEAΔ-H3K27ac-Max (0.9%) | 0.02 | 0.064 | 0.75 | 1.7 | 0.095 | 3.3e-10 |
| DeepSEAΔ-H3K4me1-Avg (0.3%) | 0.014 | 0.057 | 0.81 | 1.7 | 0.066 | 1.9e-12 |
| DeepSEAΔ-H3K4me1-Max (1.7%) | -0.051 | 0.06 | 0.4 | 1.4 | 0.061 | 2.7e-07 |
| DeepSEAΔ-H3K4me3-Avg (0.1%) | 0.078 | 0.07 | 0.27 | 2.6 | 0.16 | 0.00042 |
| DeepSEAΔ-H3K4me3-Max (0.7%) | 0.069 | 0.071 | 0.33 | 2 | 0.087 | 2.6e-13 |
| BasenjiΔ-DNase-Avg (0.3%) | -0.001 | 0.13 | 0.99 | 2.7 | 0.36 | 0.00038 |
| BasenjiΔ-DNase-Max (2.0%) | 0.14 | 0.075 | 0.071 | 2.3 | 0.17 | 1.1e-10 |
| BasenjiΔ-H3K27ac-Avg (0.3%) | 0.19 | 0.11 | 0.075 | 2.4 | 0.18 | 6.3e-11 |
| BasenjiΔ-H3K27ac-Max (0.9%) | 0.078 | 0.036 | 0.032 | 2.1 | 0.11 | 1.3e-18 |
| BasenjiΔ-H3K4me1-Avg (0.3%) | 0.14 | 0.063 | 0.027 | 2 | 0.1 | 5.6e-16 |
| BasenjiΔ-H3K4me1-Max (1.7%) | 0.13 | 0.05 | 0.0093 | 1.9 | 0.091 | 4.2e-16 |
| BasenjiΔ-H3K4me3-Avg (0.1%) | 0.16 | 0.071 | 0.023 | 3.2 | 0.19 | 6.8e-14 |
| BasenjiΔ-H3K4me3-Max (0.7%) | 0.32 | 0.053 | 7.7e-10 | 2.6 | 0.13 | 3.3e-18 |

**Table S7.**
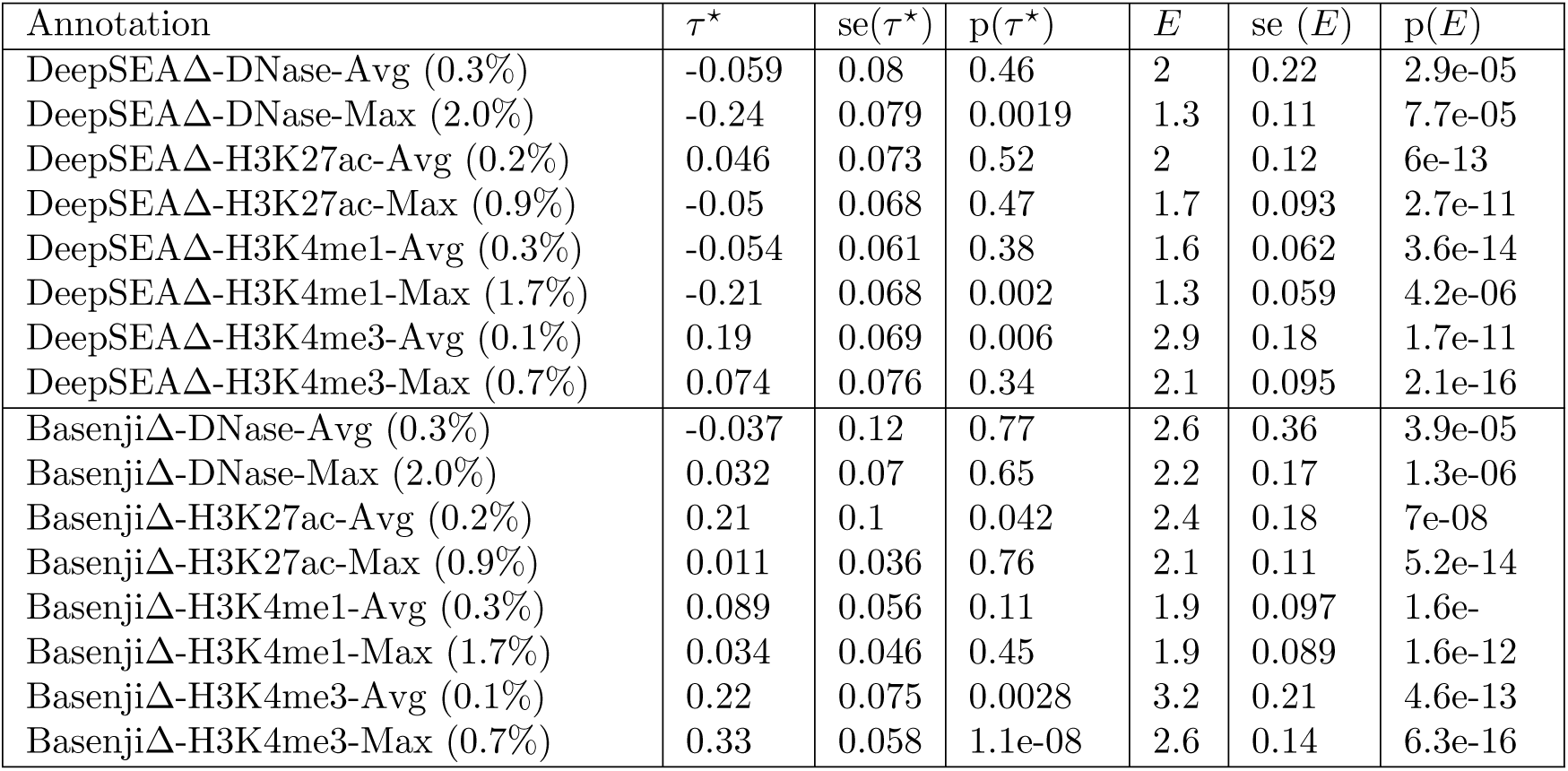
**S-LDSC results for marginal analysis of non-tissue-specific allelic-effect DeepSEA**Δ **and Basenji**Δ **annotations conditioned on baseline-LD model**. Standardized Effect sizes (*τ\**) and Enrichment (E) of 8 non-tissue-specific DeepSEAΔ and 8 non-tissue-specific BasenjiΔ annotations, when conditioned on the baseline-LD model. Results are meta-analyzed across 41 independent traits.

**Table S8.** **S-LDSC results for marginal analysis of non-tissue-specific variant-level BasenjiV annotations conditioned on the non-tissue-specific variant-level joint model and the non-tissue-specific final joint model**. Standardized Effect sizes (*τ\**) and Enrichment (E) of 8 non-tissue-specific Basenji variant level annotations, when conditioned on (Top) non-tissue-specific variant-level joint model (baseline-LD model + 7 annotations from Figure S6) and (Bottom) non-tissue-specific final joint model comprising of the non-tissue-specific variant-level joint model plus 1 significant allelic-effect annotation (BasenjiΔ-H34me3-Max) from Figure 2. Results are meta-analyzed across 41 independent traits.

| Conditional on non-tissue-specific variant-level joint model |  |  |  |  |  |  |
| --- | --- | --- | --- | --- | --- | --- |
| Annotation | $\tau^*$ | se($\tau^*$) | p($\tau^*$) | $E$ | se ($E$) | p($E$) |
| BasenjiV-DNase-Avg (1.6%) | -0.085 | 0.064 | 0.18 | 2.5 | 0.19 | 1.4e-13 |
| BasenjiV-DNase-Max (13.2%) | -0.052 | 0.06 | 0.38 | 1.8 | 0.075 | 2.3e-19 |
| BasenjiV-H3K27ac-Avg (3.2%) | 0.047 | 0.055 | 0.39 | 2.5 | 0.14 | 1.2e-17 |
| BasenjiV-H3K27ac-Max (13.7%) | -0.018 | 0.03 | 0.55 | 1.8 | 0.073 | 1e-17 |
| BasenjiV-H3K4me1-Avg (5.5%) | 0.041 | 0.04 | 0.31 | 2 | 0.075 | 1.6e-18 |
| BasenjiV-H3K4me1-Max (23.2%) | -0.046 | 0.025 | 0.067 | 1.6 | 0.046 | 1.6e-19 |
| BasenjiV-H3K4me3-Avg (1.7%) | 0.017 | 0.058 | 0.77 | 3.7 | 0.24 | 5.1e-17 |
| BasenjiV-H3K4me3-Max (8.8%) | 0.029 | 0.033 | 0.38 | 2 | 0.083 | 1.6e-17 |
| Conditional on non-tissue-specific final joint model |  |  |  |  |  |  |
| Annotation | $\tau^*$ | se($\tau^*$) | p($\tau^*$) | $E$ | se ($E$) | p($E$) |
| BasenjiV-DNase-Avg (1.6%) | -0.11 | 0.065 | 0.08 | 1.8 | 0.074 | 1.4e-38 |
| BasenjiV-DNase-Max (13.2%) | -0.11 | 0.065 | 0.099 | 2.5 | 0.19 | 8.6e-32 |
| BasenjiV-H3K27ac-Avg (3.2%) | 0.0051 | 0.055 | 0.93 | 2.5 | 0.13 | 1.3e-33 |
| BasenjiV-H3K27ac-Max (13.7%) | -0.075 | 0.03 | 0.012 | 1.8 | 0.072 | 1.5e-31 |
| BasenjiV-H3K4me1-Avg (5.5%) | -0.011 | 0.04 | 0.79 | 1.9 | 0.073 | 7.9e-32 |
| BasenjiV-H3K4me1-Max (23.2%) | -0.11 | 0.036 | 0.76 | 1.6 | 0.046 | 3.5e-33 |
| BasenjiV-H3K4me3-Avg (1.7%) | -0.031 | 0.061 | 0.61 | 3.5 | 0.23 | 5.3e-34 |
| BasenjiV-H3K4me3-Max (8.8%) | -0.026 | 0.036 | 0.47 | 1.9 | 0.081 | 3.4e-32 |

**Table S9.** **The 11 conservation related terms in the baseline-LD model**. The names of the 11 annotations related to conservation that were removed in the analysis with no conservation in Tables S10, S16 and S22. See URLs for the full list of baseline-LD annotations.

| Conservation related Annotations |
| --- |
| Conserved_LindbladToh<sup>32</sup> |
| Conserved_LindbladToh.extend.500<sup>32</sup> |
| Conserved_Vertebrate_phastCons46way<sup>31</sup> |
| Conserved_Vertebrate_phastCons46way.extend.500<sup>31</sup> |
| Conserved_Mammal_phastCons46way<sup>31</sup> |
| Conserved_Mammal_phastCons46way.extend.500<sup>31</sup> |
| Conserved_Primate_phastCons46way<sup>31</sup> |
| Conserved_Primate_phastCons46way.extend.500<sup>31</sup> |
| GERP.NS<sup>30,19</sup> |
| GERP.RSsup4<sup>30,19</sup> |
| Backgrd_Selection_Stat<sup>33</sup> |

**Table S10.** **S-LDSC results for marginal analysis of non-tissue-specific allelic-effect DeepSEA**Δ **and Basenji**Δ **annotations conditioned on the non-tissue-specific variant-level model without any conservation related annotations**. Standardized Effect sizes (*τ\**) and Enrichment (E) of 8 non-tissue-specific DeepSEAΔ and 8 non-tissue-specific BasenjiΔ annotations, when conditioned on the non-tissue-specific variant-level model (baseline-LD model + 7 annotations from Figure S6) without any conservation related annotations. Results are meta-analyzed across 41 independent traits.

| Annotation | $\tau^*$ | se($\tau^*$) | p($\tau^*$) | $E$ | se ($E$) | p($E$) |
| --- | --- | --- | --- | --- | --- | --- |
| DeepSEAΔ-DNase-Avg (0.3%) | 0.34 | 0.083 | 3.4e-05 | 3 | 0.19 | 4.7e-13 |
| DeepSEAΔ-DNase-Max (2.0%) | 0.32 | 0.093 | 0.00057 | 2.1 | 0.12 | 7.6e-13 |
| DeepSEAΔ-H3K27ac-Avg (0.2%) | 0.44 | 0.088 | 6.3e-07 | 2.6 | 0.12 | 1.1e-24 |
| DeepSEAΔ-H3K27ac-Max (0.9%) | 0.4 | 0.077 | 2.4e-07 | 2.2 | 0.094 | 4.6e-22 |
| DeepSEAΔ-H3K4me1-Avg (0.3%) | 0.38 | 0.074 | 2e-07 | 2.1 | 0.067 | 2.4e-24 |
| DeepSEAΔ-H3K4me1-Max (1.7%) | 0.35 | 0.082 | 2.1e-05 | 1.9 | 0.068 | 7.1e-18 |
| DeepSEAΔ-H3K4me3-Avg (0.1%) | 0.34 | 0.087 | 7.3e-05 | 3.5 | 0.19 | 2.7e-18 |
| DeepSEAΔ-H3K4me3-Max (0.7%) | 0.3 | 0.085 | 0.00043 | 2.4 | 0.098 | 3.7e-19 |
| BasenjiΔ-DNase-Avg (0.3%) | 0.21 | 0.13 | 0.12 | 3.1 | 0.36 | 1.6e-14 |
| BasenjiΔ-DNase-Max (2.0%) | 0.33 | 0.07 | 3e-06 | 2.4 | 0.16 | 8.7e-22 |
| BasenjiΔ-H3K27ac-Avg (0.2%) | 0.23 | 0.12 | 0.049 | 2.4 | 0.19 | 9e-26 |
| BasenjiΔ-H3K27ac-Max (0.9%) | 0.11 | 0.039 | 0.0071 | 2 | 0.11 | 2.2e-24 |
| BasenjiΔ-H3K4me1-Avg (0.3%) | 0.28 | 0.066 | 1.5e-05 | 2 | 0.1 | 8.9e-27 |
| BasenjiΔ-H3K4me1-Max (1.7%) | 0.22 | 0.04 | 7.1e-08 | 1.9 | 0.088 | 2.3e-24 |
| BasenjiΔ-H3K4me3-Avg (0.1%) | 0.18 | 0.076 | 0.015 | 3.4 | 0.21 | 7.6e-25 |
| BasenjiΔ-H3K4me3-Max (0.7%) | 0.3 | 0.058 | 1.7e-07 | 2.6 | 0.15 | 2.1e-28 |

**Table S11.**
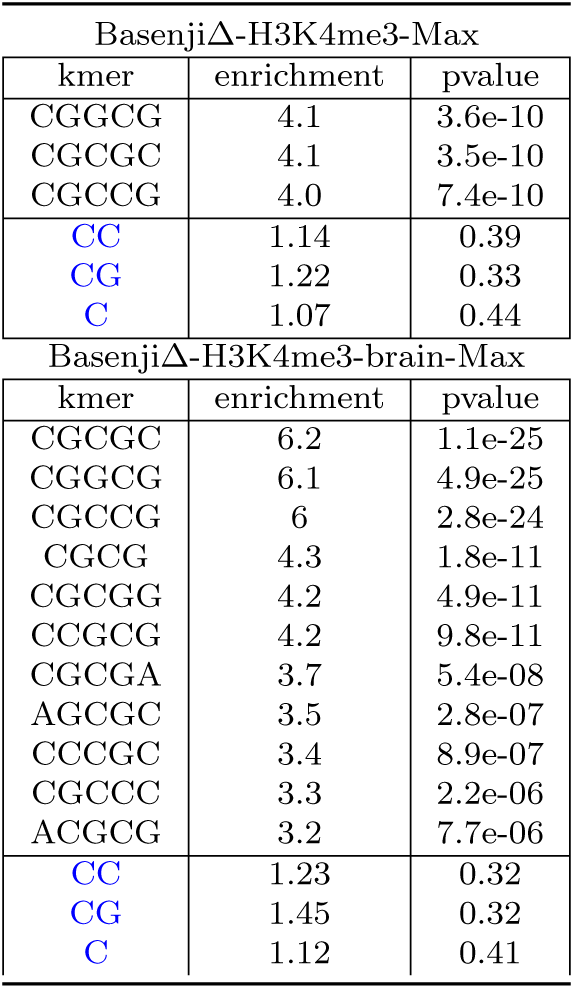
**Weighted** k**-mer enrichments for significant allelic-effect Basenji annotations**. We report the weighted *k*-mer enrichment and enrichment p-value for the top significantly enriched *k*-mers (1≤*k*≤5) for the two allelic effect annotations found significant: BasenjiΔ-H3K4me3-Max and BasenjiΔ-H3K4me3-brain-Max annotations. The Bonferonni corrcection threshold is 0.05*/*(4 682) where 682 is the number of *k*-mers analyzed for the 2 significant variant-level annotations analyzed in Table S50 and the 2 significant allelic-effect annotations analyzed here. We also report results for simple GC-rich motifs (blue font) for comparison purposes.

| BasenjiΔ-H3K4me3-Max | | |
| --- | --- | --- |
| kmer | enrichment | pvalue |
| CGGCG | 4.1 | 3.6e-10 |
| CGCGC | 4.1 | 3.5e-10 |
| CGCCG | 4.0 | 7.4e-10 |
| CC | 1.14 | 0.39 |
| CG | 1.22 | 0.33 |
| C | 1.07 | 0.44 |
| BasenjiΔ-H3K4me3-brain-Max | | |
| kmer | enrichment | pvalue |
| CGCGC | 6.2 | 1.1e-25 |
| CGGCG | 6.1 | 4.9e-25 |
| CGCCG | 6 | 2.8e-24 |
| CGCG | 4.3 | 1.8e-11 |
| CGCGG | 4.2 | 4.9e-11 |
| CCGCG | 4.2 | 9.8e-11 |
| CGCGA | 3.7 | 5.4e-08 |
| AGCGC | 3.5 | 2.8e-07 |
| CCCGC | 3.4 | 8.9e-07 |
| CGCCC | 3.3 | 2.2e-06 |
| ACGCG | 3.2 | 7.7e-06 |
| CC | 1.23 | 0.32 |
| CG | 1.45 | 0.32 |
| C | 1.12 | 0.41 |

**Table S12.** **Enrichment of two 9-mers of interest for significant allelic-effect Basenji**Δ **annotations**. We report the weighted *k*-mer enrichment and enrichment p-value of two 9-mers, GCGGTGGCT and GTGGTGGCT, with previously reported evidence of being connected to trait architecture^36^, with respect to the two allelic effect annotations found significant: BasenjiΔ-H3K4me3-Max and BasenjiΔ-H3K4me3-brain-Max annotations. We additionally also compare against the same annotation for blood (BasenjiΔ-H3K4me3-blood-Max) for a better comparison.

| BasenjiΔ-H3K4me3-Max | | |
| --- | --- | --- |
| kmer | enrichment | pvalue |
| GCGGTGGCT | 1.12 | 9.7e-08 |
| GTGGTGGCT | 0.96 | 1.00 |
| BasenjiΔ-H3K4me3-brain-Max | | |
| kmer | enrichment | pvalue |
| GCGGTGGCT | 1.03 | 0.003 |
| GTGGTGGCT | 0.85 | 1.00 |
| BasenjiΔ-H3K4me3-blood-Max | | |
| kmer | enrichment | pvalue |
| GCGGTGGCT | 1.19 | 3.3e-19 |
| GTGGTGGCT | 0.93 | 1.00 |

**Table S13.** **AUROC of various sets of annotations incorporating non-tissue-specific allelic-effect deep learning annotations in predicting** 12, 296 **NIH GWAS SNPs**. We report the AUROC for a gradient boosting model trained using each respective set of annotations, either without including baseline-LD model annotations; including baseline-LD model annotations; or including annotations from the non-tissue-specific variant-level joint model (baseline-LD model + 7 annotations; baseline-LD+7).

| Feature | GWAS SNPs |
| --- | --- |
| DeepSEAΔ-Avg/Max | 0.584 |
| BasenjiΔ-Avg/Max | 0.592 |
| DeepSEAΔ-All | 0.602 |
| BasenjiΔ-All | 0.611 |
| baseline-LD | 0.758 |
| baseline-LD + DeepSEAΔ-Avg/Max | 0.767 |
| baseline-LD + BasenjiΔ-Avg/Max | 0.767 |
| baseline-LD + DeepSEAΔ-All | 0.770 |
| baseline-LD + BasenjiΔ-All | 0.772 |
| baseline-LD+7 | 0.762 |
| baseline-LD+7 + DeepSEAΔ-Avg/Max | 0.766 |
| baseline-LD+7 + BasenjiΔ-Avg/Max | 0.769 |
| baseline-LD+7 + DeepSEAΔ-All | 0.771 |
| baseline-LD+7 + BasenjiΔ-All | 0.773 |

**Table S14.** **Enrichment of NIH GWAS SNPs for non-tissue-specific allelic-effect annotations**. We report the enrichment in deep learning annotations (along with Jackknife standard error) at top 12, 296 known disease-associated SNPs from NIH GWAS catalog^23^.

| Feature | GWAS SNPs |
| --- | --- |
| BasenjiΔ-DNase-Avg | 1.93 (0.020) |
| BasenjiΔ-DNase-Max | 1.70 (0.012) |
| BasenjiΔ-H3K27ac-Avg | 1.76 (0.01) |
| BasenjiΔ-H3K27ac-Max | 1.63 (0.009) |
| BasenjiΔ-H3K4me1-Avg | 1.54 (0.008) |
| BasenjiΔ-H3K4me1-Max | 1.53 (0.008) |
| BasenjiΔ-H3K4me3-Avg | 2.08 (0.022) |
| BasenjiΔ-H3K4me3-Max | 1.79 (0.011) |
| DeepSEAΔ-DNase-Avg | 1.64 (0.01) |
| DeepSEAΔ-DNase-Max | 1.41 (0.007) |
| DeepSEAΔ-H3K27ac-Avg | 1.57 (0.009) |
| DeepSEAΔ-H3K27ac-Max | 1.46 (0.008) |
| DeepSEAΔ-H3K4me1-Avg | 1.43 (0.007) |
| DeepSEAΔ-H3K4me1-Max | 1.33 (0.006) |
| DeepSEAΔ-H3K4me3-Avg | 1.67 (0.013) |
| DeepSEAΔ-H3K4me3-Max | 1.50 (0.007) |

**Table S15.** **S-LDSC results for marginal analysis of blood-specific allelic-effect deep learning annotations against blood-specific variant-level joint model and 1 significant non-tissue-specific Basenji**Δ **annotation**. Standardized Effect sizes (*τ\**) and Enrichment (E) of 8 blood-specific DeepSEAΔ and 8 blood-specific BasenjiΔ annotations, when conditioned on the blood-specific joint model annotations (baseline-LD model + 7 non-tissue-specific annotations from Figure S6 + 6 blood-specific Roadmap and ChromHMM annotations from Figure S11) and 1 significant allelic-effect annotation (BasenjiΔ-H3K4me3-Max) from Table S6. Results are meta-analyzed across 11 blood-related traits.

| Annotation | $\tau^*$ | se($\tau^*$) | p($\tau^*$) | $E$ | se ($E$) | p($E$) |
| --- | --- | --- | --- | --- | --- | --- |
| DeepSEAΔ-DNase-blood-Avg (0.2%) | -0.4 | 0.12 | 0.00074 | 2.3 | 0.38 | 1.8e-05 |
| DeepSEAΔ-DNase-blood-Max (0.6%) | -0.35 | 0.11 | 0.0011 | 2 | 0.26 | 5.1e-11 |
| DeepSEAΔ-H3K27ac-blood-Avg (0.2%) | -0.23 | 0.11 | 0.028 | 2.6 | 0.23 | 1.6e-08 |
| DeepSEAΔ-H3K27ac-blood-Max (0.5%) | -0.34 | 0.11 | 0.002 | 2.1 | 0.16 | 1.8e-08 |
| DeepSEAΔ-H3K4me1-blood-Avg (0.3%) | -0.35 | 0.13 | 0.0035 | 1.9 | 0.16 | 1.3e-08 |
| DeepSEAΔ-H3K4me1-blood-Max (0.8%) | -0.34 | 0.12 | 0.0023 | 1.7 | 0.14 | 6.4e-07 |
| DeepSEAΔ-H3K4me3-blood-Avg (0.1%) | -0.3 | 0.11 | 0.0063 | 2.7 | 0.35 | 4.3e-08 |
| DeepSEAΔ-H3K4me3-blood-Max (0.4%) | -0.38 | 0.14 | 0.0056 | 2.2 | 0.24 | 7.9e-10 |
| BasenjiΔ-DNase-blood-Avg (0.2%) | -0.31 | 0.2 | 0.12 | 5 | 0.41 | 9.5e-11 |
| BasenjiΔ-DNase-blood-Max (0.6%) | -0.012 | 0.22 | 0.96 | 4.7 | 0.28 | 4e-08 |
| BasenjiΔ-H3K27ac-blood-Avg (0.2%) | 0.34 | 0.17 | 0.046 | 4.8 | 0.29 | 1.5e-08 |
| BasenjiΔ-H3K27ac-blood-Max (0.5%) | 0.23 | 0.17 | 0.17 | 4 | 0.19 | 2.9e-08 |
| BasenjiΔ-H3K4me1-blood-Avg (0.3%) | 0.033 | 0.1 | 0.75 | 3.6 | 0.2 | 4.9e-09 |
| BasenjiΔ-H3K4me1-blood-Max (0.8%) | -0.006 | 0.15 | 0.97 | 3.3 | 0.17 | 1.3e-08 |
| BasenjiΔ-H3K4me3-blood-Avg (0.1%) | 0.57 | 0.33 | 0.083 | 6.5 | 0.55 | 2.6e-09 |
| BasenjiΔ-H3K4me3-blood-Max (0.4%) | 0.39 | 0.3 | 0.21 | 4.7 | 0.3 | 7.8e-09 |

**Table S16.** **S-LDSC results for marginal analysis of blood-specific allelic-effect deep learning annotations against blood-specific variant-level joint model and 1 significant non-tissue-specific Basenji**Δ **annotation but without any conservation related annotation**. Standardized Effect sizes (*τ\**) and Enrichment (E) of 8 blood-specific DeepSEAΔ and 8 blood-specific BasenjiΔ annotations, when conditioned on the blood-specific variant-level joint model (baseline-LD model + 7 non-tissue-specific annotations from Figure S6 + 6 blood-specific Roadmap and ChromHMM annotations from Figure S11) and BasenjiΔ-H3K4me3-Max but without the conservation related annotation. Results are meta-analyzed across 11 blood-related traits.

| Annotation | $\tau^*$ | se($\tau^*$) | p($\tau^*$) | $E$ | se ($E$) | p($E$) |
| --- | --- | --- | --- | --- | --- | --- |
| DeepSEAΔ-DNase-blood-Avg (0.2%) | -0.17 | 0.13 | 0.18 | 2.6 | 0.38 | 0.0089 |
| DeepSEAΔ-DNase-blood-Max (0.6%) | -0.067 | 0.1 | 0.51 | 2.3 | 0.26 | 0.0035 |
| DeepSEAΔ-H3K27ac-blood-Avg (0.2%) | 0.078 | 0.1 | 0.45 | 3.1 | 0.23 | 1.9e-05 |
| DeepSEAΔ-H3K27ac-blood-Max (0.5%) | -0.035 | 0.1 | 0.74 | 2.4 | 0.16 | 0.00016 |
| DeepSEAΔ-H3K4me1-blood-Avg (0.3%) | -0.026 | 0.087 | 0.76 | 2.2 | 0.16 | 0.00011 |
| DeepSEAΔ-H3K4me1-blood-Max (0.8%) | -0.007 | 0.085 | 0.93 | 2 | 0.14 | 0.00054 |
| DeepSEAΔ-H3K4me3-blood-Avg (0.1%) | -0.031 | 0.1 | 0.77 | 3.3 | 0.35 | 0.00039 |
| DeepSEAΔ-H3K4me3-blood-Max (0.4%) | -0.11 | 0.14 | 0.43 | 2.6 | 0.23 | 0.00079 |
| BasenjiΔ-DNase-blood-Avg (0.2%) | -0.042 | 0.17 | 0.8 | 5 | 0.41 | 2.2e-05 |
| BasenjiΔ-DNase-blood-Max (0.6%) | 0.13 | 0.18 | 0.47 | 4.6 | 0.29 | 6.2e-07 |
| BasenjiΔ-H3K27ac-blood-Avg (0.2%) | 0.42 | 0.12 | 0.00039 | 4.8 | 0.29 | 5.1e-09 |
| BasenjiΔ-H3K27ac-blood-Max (0.5%) | 0.32 | 0.11 | 0.0047 | 4 | 0.2 | 3.5e-09 |
| BasenjiΔ-H3K4me1-blood-Avg (0.3%) | 0.19 | 0.094 | 0.044 | 3.5 | 0.2 | 9.8e-09 |
| BasenjiΔ-H3K4me1-blood-Max (0.8%) | 0.14 | 0.11 | 0.17 | 3.2 | 0.17 | 3.8e-08 |
| BasenjiΔ-H3K4me3-blood-Avg (0.1%) | 0.6 | 0.2 | 0.0021 | 6.5 | 0.49 | 1.9e-07 |
| BasenjiΔ-H3K4me3-blood-Max (0.4%) | 0.4 | 0.14 | 0.0049 | 4.7 | 0.3 | 4.9e-09 |

**Table S17.**
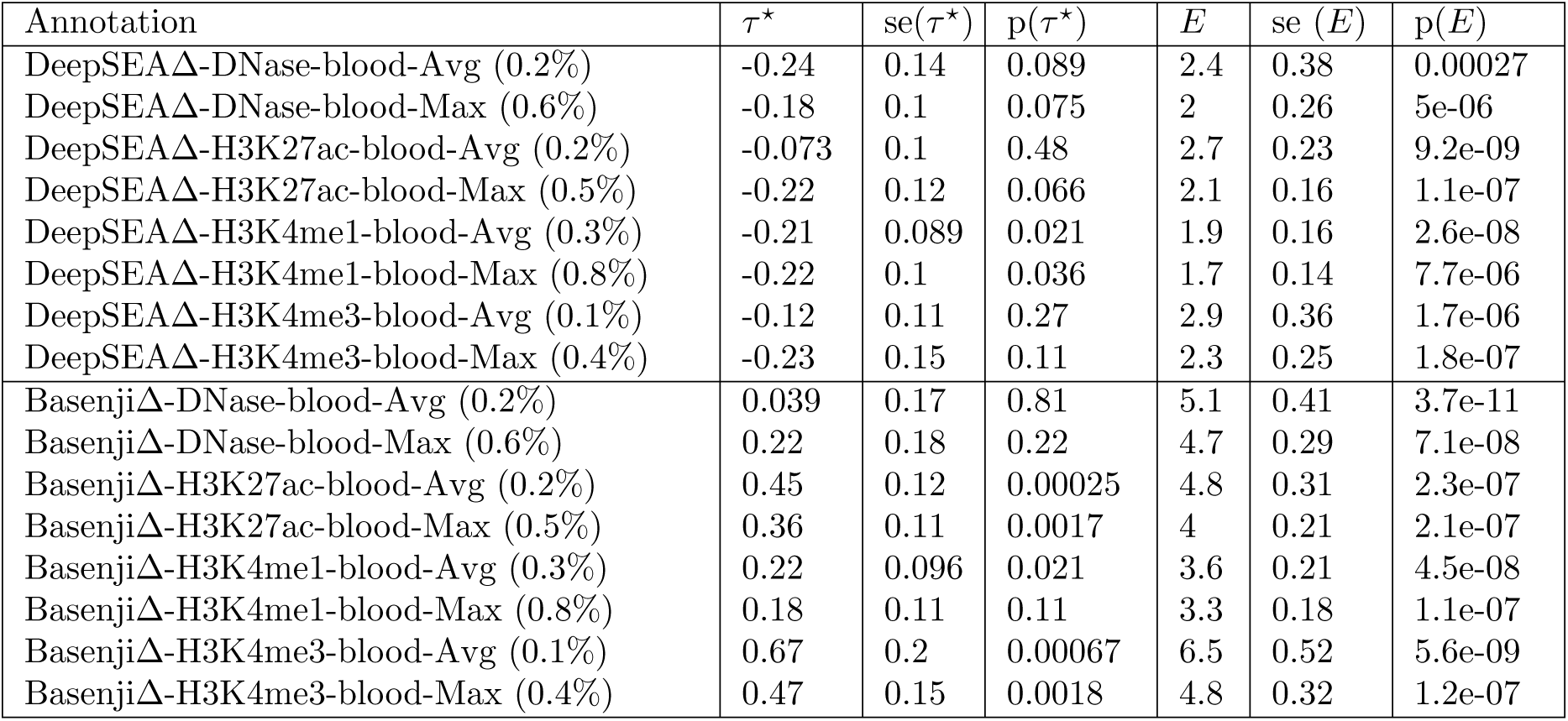
**S-LDSC results for marginal analysis of blood-specific allelic-effect deep learning annotations against blood-specific variant-level joint model**. Standardized Effect sizes (*τ\**) and Enrichment (E) of 8 blood-specific DeepSEAΔ and 8 blood-specific BasenjiΔ annotations, when conditioned on the blood-specific variant-level joint model annotations (baseline-LD model + 7 non-tissue-specific annotations from Figure S6 + 6 blood-specific Roadmap and ChromHMM annotations from Figure S11). Results are meta-analyzed across 11 blood-related traits.

| Annotation | $\tau^*$ | se($\tau^*$) | p($\tau^*$) | $E$ | se ($E$) | p($E$) |
| --- | --- | --- | --- | --- | --- | --- |
| DeepSEAΔ-DNase-blood-Avg (0.2%) | -0.24 | 0.14 | 0.089 | 2.4 | 0.38 | 0.00027 |
| DeepSEAΔ-DNase-blood-Max (0.6%) | -0.18 | 0.1 | 0.075 | 2 | 0.26 | 5e-06 |
| DeepSEAΔ-H3K27ac-blood-Avg (0.2%) | -0.073 | 0.1 | 0.48 | 2.7 | 0.23 | 9.2e-09 |
| DeepSEAΔ-H3K27ac-blood-Max (0.5%) | -0.22 | 0.12 | 0.066 | 2.1 | 0.16 | 1.1e-07 |
| DeepSEAΔ-H3K4me1-blood-Avg (0.3%) | -0.21 | 0.089 | 0.021 | 1.9 | 0.16 | 2.6e-08 |
| DeepSEAΔ-H3K4me1-blood-Max (0.8%) | -0.22 | 0.1 | 0.036 | 1.7 | 0.14 | 7.7e-06 |
| DeepSEAΔ-H3K4me3-blood-Avg (0.1%) | -0.12 | 0.11 | 0.27 | 2.9 | 0.36 | 1.7e-06 |
| DeepSEAΔ-H3K4me3-blood-Max (0.4%) | -0.23 | 0.15 | 0.11 | 2.3 | 0.25 | 1.8e-07 |
| BasenjiΔ-DNase-blood-Avg (0.2%) | 0.039 | 0.17 | 0.81 | 5.1 | 0.41 | 3.7e-11 |
| BasenjiΔ-DNase-blood-Max (0.6%) | 0.22 | 0.18 | 0.22 | 4.7 | 0.29 | 7.1e-08 |
| BasenjiΔ-H3K27ac-blood-Avg (0.2%) | 0.45 | 0.12 | 0.00025 | 4.8 | 0.31 | 2.3e-07 |
| BasenjiΔ-H3K27ac-blood-Max (0.5%) | 0.36 | 0.11 | 0.0017 | 4 | 0.21 | 2.1e-07 |
| BasenjiΔ-H3K4me1-blood-Avg (0.3%) | 0.22 | 0.096 | 0.021 | 3.6 | 0.21 | 4.5e-08 |
| BasenjiΔ-H3K4me1-blood-Max (0.8%) | 0.18 | 0.11 | 0.11 | 3.3 | 0.18 | 1.1e-07 |
| BasenjiΔ-H3K4me3-blood-Avg (0.1%) | 0.67 | 0.2 | 0.00067 | 6.5 | 0.52 | 5.6e-09 |
| BasenjiΔ-H3K4me3-blood-Max (0.4%) | 0.47 | 0.15 | 0.0018 | 4.8 | 0.32 | 1.2e-07 |

**Table S18.** **S-LDSC results for marginal analysis of blood-specific allelic-effect deep learning annotations conditioned on the non-tissue-specific final joint model**. Standardized Effect sizes (*τ\**) and Enrichment (E) of 8 blood-specific DeepSEAΔ and 8 blood-specific BasenjiΔ annotations, when conditioned on the non-tissue-specific final joint model. Results are meta-analyzed across 11 blood-related traits.

| Annotation | $\tau^*$ | se($\tau^*$) | p($\tau^*$) | $E$ | se ($E$) | p($E$) |
| --- | --- | --- | --- | --- | --- | --- |
| DeepSEAΔ-DNase-blood-Avg (0.2%) | -0.37 | 0.14 | 0.0064 | 2.9 | 0.52 | 1.8e-06 |
| DeepSEAΔ-DNase-blood-Max (0.6%) | -0.49 | 0.12 | 2.4e-05 | 2 | 0.28 | 1.6e-06 |
| DeepSEAΔ-H3K27ac-blood-Avg (0.2%) | 0.024 | 0.11 | 0.83 | 3.3 | 0.29 | 1.2e-18 |
| DeepSEAΔ-H3K27ac-blood-Max (0.5%) | -0.23 | 0.11 | 0.038 | 2.4 | 0.2 | 1.2e-16 |
| DeepSEAΔ-H3K4me1-blood-Avg (0.3%) | -0.33 | 0.099 | 0.0011 | 2 | 0.18 | 2e-10 |
| DeepSEAΔ-H3K4me1-blood-Max (0.8%) | -0.47 | 0.098 | 1.4e-06 | 1.7 | 0.15 | 4.8e-08 |
| DeepSEAΔ-H3K4me3-blood-Avg (0.1%) | -0.071 | 0.12 | 0.56 | 3.8 | 0.41 | 1.4e-08 |
| DeepSEAΔ-H3K4me3-blood-Max (0.4%) | -0.23 | 0.13 | 0.078 | 2.7 | 0.28 | 5.7e-09 |
| BasenjiΔ-DNase-blood-Avg (0.2%) | 0.56 | 0.2 | 0.0049 | 7.2 | 0.57 | 1.6e-09 |
| BasenjiΔ-DNase-blood-Max (0.6%) | 0.69 | 0.21 | 0.00081 | 5.8 | 0.37 | 5.1e-09 |
| BasenjiΔ-H3K27ac-blood-Avg (0.2%) | 1.8 | 0.26 | 1.5e-12 | 6.2 | 0.38 | 6.6e-09 |
| BasenjiΔ-H3K27ac-blood-Max (0.5%) | 1.5 | 0.22 | 1.1e-11 | 4.8 | 0.25 | 1.5e-08 |
| BasenjiΔ-H3K4me1-blood-Avg (0.3%) | 1.2 | 0.2 | 7.7e-09 | 4.6 | 0.28 | 2.1e-09 |
| BasenjiΔ-H3K4me1-blood-Max (0.8%) | 1 | 0.19 | 8.7e-08 | 4 | 0.22 | 3.4e-09 |
| BasenjiΔ-H3K4me3-blood-Avg (0.1%) | 1.9 | 0.36 | 1.3e-07 | 8.6 | 0.68 | 3e-09 |
| BasenjiΔ-H3K4me3-blood-Max (0.4%) | 2.1 | 0.36 | 7.4e-09 | 5.7 | 0.34 | 1.3e-08 |

**Table S19.** **AUROC of various sets of annotations incorporating blood-specific allelic-effect deep learning annotations in predicting 8,741 fine-mapped autoimmune disease SNPs**. We report the AUROC for a gradient boosting model trained using each respective set of annotations, either without including baseline-LD model annotations; including baseline-LD model annotations; or including annotations from the blood-specific variant-level joint model plus BasenjiΔ-H3K4me3-Max (baseline-LD+7+6blood+1).

| Feature | Farh et al |
| --- | --- |
| DeepSEAΔ-blood-Avg/Max | 0.613 |
| BasenjiΔ-blood-Avg/Max | 0.672 |
| DeepSEAΔ-blood-All | 0.633 |
| BasenjiΔ-blood-All | 0.684 |
| baseline-LD | 0.841 |
| baseline-LD + DeepSEAΔ-blood-Avg/Max | 0.847 |
| baseline-LD + BasenjiΔ-blood-Avg/Max | 0.852 |
| baseline-LD + DeepSEAΔ-blood-All | 0.848 |
| baseline-LD + BasenjiΔ-blood-All | 0.851 |
| baseline-LD+7+6blood+1 | 0.848 |
| baseline-LD+7+6blood+1 + DeepSEAΔ-blood-Avg/Max | 0.847 |
| baseline-LD+7+6blood+1 + BasenjiΔ-blood-Avg/Max | 0.851 |
| baseline-LD+7+6blood+1 + DeepSEAΔ-blood-All | 0.849 |
| baseline-LD+7+6blood+1 + BasenjiΔ-blood-All | 0.851 |

**Table S20.** **Enrichment of fine-mapped autoimmune disease SNPs for blood-specific allelic-effect annotations**. We report the enrichment in blood-specific deep learning allelic-effect annotations (along with Jackknife standard error) for 8741 fine-mapped SNPs in immune-related traits^24^.

| Feature | Farh et al |
| --- | --- |
| BasenjiΔ-DNase-blood-Avg | 2.59 (0.05) |
| BasenjiΔ-DNase-blood-Max | 2.41 (0.04) |
| BasenjiΔ-H3K27ac-blood-Avg | 2.44 (0.03) |
| BasenjiΔ-H3K27ac-blood-Max | 2.29 (0.03) |
| BasenjiΔ-H3K4me1-blood-Avg | 2.21 (0.03) |
| BasenjiΔ-H3K4me1-blood-Max | 2.09 (0.02) |
| BasenjiΔ-H3K4me3-blood-Avg | 2.52 (0.03) |
| BasenjiΔ-H3K4me3-blood-Max | 2.40 (0.03) |
| DeepSEAΔ-DNase-blood-Avg | 1.87 (0.03) |
| DeepSEAΔ-DNase-blood-Max | 1.62 (0.02) |
| DeepSEAΔ-H3K27ac-blood-Avg | 1.83 (0.03) |
| DeepSEAΔ-H3K27ac-blood-Max | 1.60 (0.01) |
| DeepSEAΔ-H3K4me1-blood-Avg | 1.58 (0.01) |
| DeepSEAΔ-H3K4me1-blood-Max | 1.45 (0.01) |
| DeepSEAΔ-H3K4me3-blood-Avg | 2.01 (0.04) |
| DeepSEAΔ-H3K4me3-blood-Max | 1.66 (0.02) |

**Table S21.**
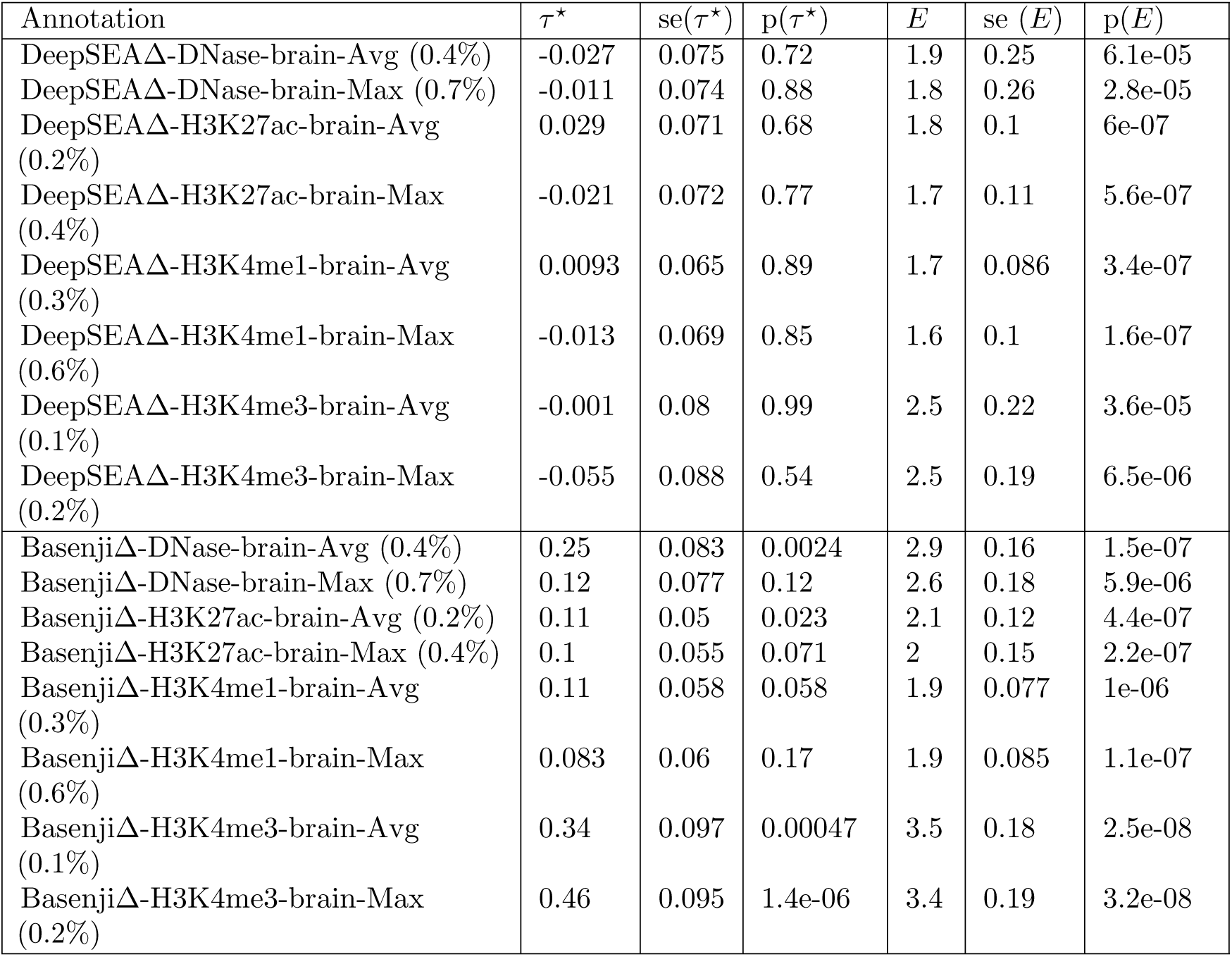
**S-LDSC results for marginal analysis of brain-specific allelic-effect deep learning annotations against brain-specific variant-level joint model and 1 significant non-tissue-specific Basenji**Δ **annotation**. Standardized Effect sizes (*τ\**) and Enrichment (E) of 8 brain-specific DeepSEAΔ and 8 brain-specific BasenjiΔ annotations, when conditioned on brain-specific variant-level joint model (baseline-LD model + 7 non-tissue-specific annotations from Figure S6 + DeepSEAV-H3K4me3-brain-Max and BasenjiV-H3K27ac-brain-Max (the 2 significant brain-specific variant-level annotations; Figure S12) + 4 additional brain-specific annotations from Figure S15) and 1 significant non-tissue-specific annotation from Table S6). Results are meta-analyzed across 8 brain-related traits.

| Annotation | $\tau^*$ | se($\tau^*$) | p($\tau^*$) | $E$ | se ($E$) | p($E$) |
| --- | --- | --- | --- | --- | --- | --- |
| DeepSEAΔ-DNase-brain-Avg (0.4%) | -0.027 | 0.075 | 0.72 | 1.9 | 0.25 | 6.1e-05 |
| DeepSEAΔ-DNase-brain-Max (0.7%) | -0.011 | 0.074 | 0.88 | 1.8 | 0.26 | 2.8e-05 |
| DeepSEAΔ-H3K27ac-brain-Avg (0.2%) | 0.029 | 0.071 | 0.68 | 1.8 | 0.1 | 6e-07 |
| DeepSEAΔ-H3K27ac-brain-Max (0.4%) | -0.021 | 0.072 | 0.77 | 1.7 | 0.11 | 5.6e-07 |
| DeepSEAΔ-H3K4me1-brain-Avg (0.3%) | 0.0093 | 0.065 | 0.89 | 1.7 | 0.086 | 3.4e-07 |
| DeepSEAΔ-H3K4me1-brain-Max (0.6%) | -0.013 | 0.069 | 0.85 | 1.6 | 0.1 | 1.6e-07 |
| DeepSEAΔ-H3K4me3-brain-Avg (0.1%) | -0.001 | 0.08 | 0.99 | 2.5 | 0.22 | 3.6e-05 |
| DeepSEAΔ-H3K4me3-brain-Max (0.2%) | -0.055 | 0.088 | 0.54 | 2.5 | 0.19 | 6.5e-06 |
| BasenjiΔ-DNase-brain-Avg (0.4%) | 0.25 | 0.083 | 0.0024 | 2.9 | 0.16 | 1.5e-07 |
| BasenjiΔ-DNase-brain-Max (0.7%) | 0.12 | 0.077 | 0.12 | 2.6 | 0.18 | 5.9e-06 |
| BasenjiΔ-H3K27ac-brain-Avg (0.2%) | 0.11 | 0.05 | 0.023 | 2.1 | 0.12 | 4.4e-07 |
| BasenjiΔ-H3K27ac-brain-Max (0.4%) | 0.1 | 0.055 | 0.071 | 2 | 0.15 | 2.2e-07 |
| BasenjiΔ-H3K4me1-brain-Avg (0.3%) | 0.11 | 0.058 | 0.058 | 1.9 | 0.077 | 1e-06 |
| BasenjiΔ-H3K4me1-brain-Max (0.6%) | 0.083 | 0.06 | 0.17 | 1.9 | 0.085 | 1.1e-07 |
| BasenjiΔ-H3K4me3-brain-Avg (0.1%) | 0.34 | 0.097 | 0.00047 | 3.5 | 0.18 | 2.5e-08 |
| BasenjiΔ-H3K4me3-brain-Max (0.2%) | 0.46 | 0.095 | 1.4e-06 | 3.4 | 0.19 | 3.2e-08 |

**Table S22.**
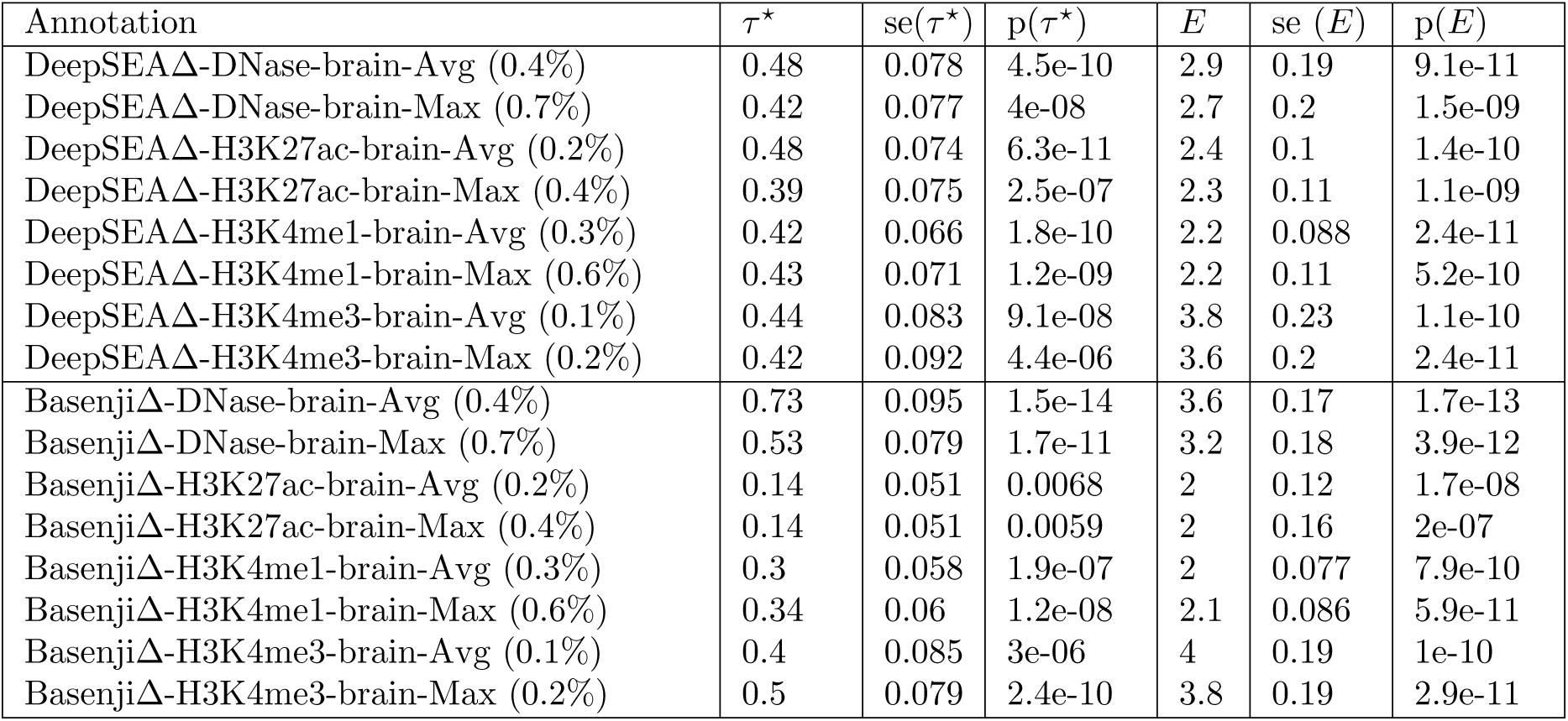
**S-LDSC results for marginal analysis of brain-specific allelic-effect deep learning annotations against brain-specific variant-level joint model and 1 significant non-tissue-specific Basenji**Δ **annotation but without any conservation related annotations**. Standardized Effect sizes (*τ\**) and Enrichment (E) of 8 brain-specific DeepSEAΔ and 8 brain-specific BasenjiΔ annotations, when conditioned on the brain-specific variant-level joint model (baseline-LD model + 7 non-tissue-specific annotations from Figure S6 + DeepSEAV-H3K4me3-brain-Max and BasenjiV-H3K27ac-brain-Max (the 2 significant brain-specific variant-level annotations; Figure S12) + 4 additional brain-specific annotations from Figure S15) and BasenjiΔ-H3K4me3-Max but without the conservation related annotations. Results are meta-analyzed across 8 brain-related traits.

| Annotation | $\tau^*$ | se($\tau^*$) | p($\tau^*$) | $E$ | se ($E$) | p($E$) |
| --- | --- | --- | --- | --- | --- | --- |
| DeepSEAΔ-DNase-brain-Avg (0.4%) | 0.48 | 0.078 | 4.5e-10 | 2.9 | 0.19 | 9.1e-11 |
| DeepSEAΔ-DNase-brain-Max (0.7%) | 0.42 | 0.077 | 4e-08 | 2.7 | 0.2 | 1.5e-09 |
| DeepSEAΔ-H3K27ac-brain-Avg (0.2%) | 0.48 | 0.074 | 6.3e-11 | 2.4 | 0.1 | 1.4e-10 |
| DeepSEAΔ-H3K27ac-brain-Max (0.4%) | 0.39 | 0.075 | 2.5e-07 | 2.3 | 0.11 | 1.1e-09 |
| DeepSEAΔ-H3K4me1-brain-Avg (0.3%) | 0.42 | 0.066 | 1.8e-10 | 2.2 | 0.088 | 2.4e-11 |
| DeepSEAΔ-H3K4me1-brain-Max (0.6%) | 0.43 | 0.071 | 1.2e-09 | 2.2 | 0.11 | 5.2e-10 |
| DeepSEAΔ-H3K4me3-brain-Avg (0.1%) | 0.44 | 0.083 | 9.1e-08 | 3.8 | 0.23 | 1.1e-10 |
| DeepSEAΔ-H3K4me3-brain-Max (0.2%) | 0.42 | 0.092 | 4.4e-06 | 3.6 | 0.2 | 2.4e-11 |
| BasenjiΔ-DNase-brain-Avg (0.4%) | 0.73 | 0.095 | 1.5e-14 | 3.6 | 0.17 | 1.7e-13 |
| BasenjiΔ-DNase-brain-Max (0.7%) | 0.53 | 0.079 | 1.7e-11 | 3.2 | 0.18 | 3.9e-12 |
| BasenjiΔ-H3K27ac-brain-Avg (0.2%) | 0.14 | 0.051 | 0.0068 | 2 | 0.12 | 1.7e-08 |
| BasenjiΔ-H3K27ac-brain-Max (0.4%) | 0.14 | 0.051 | 0.0059 | 2 | 0.16 | 2e-07 |
| BasenjiΔ-H3K4me1-brain-Avg (0.3%) | 0.3 | 0.058 | 1.9e-07 | 2 | 0.077 | 7.9e-10 |
| BasenjiΔ-H3K4me1-brain-Max (0.6%) | 0.34 | 0.06 | 1.2e-08 | 2.1 | 0.086 | 5.9e-11 |
| BasenjiΔ-H3K4me3-brain-Avg (0.1%) | 0.4 | 0.085 | 3e-06 | 4 | 0.19 | 1e-10 |
| BasenjiΔ-H3K4me3-brain-Max (0.2%) | 0.5 | 0.079 | 2.4e-10 | 3.8 | 0.19 | 2.9e-11 |

**Table S23.**
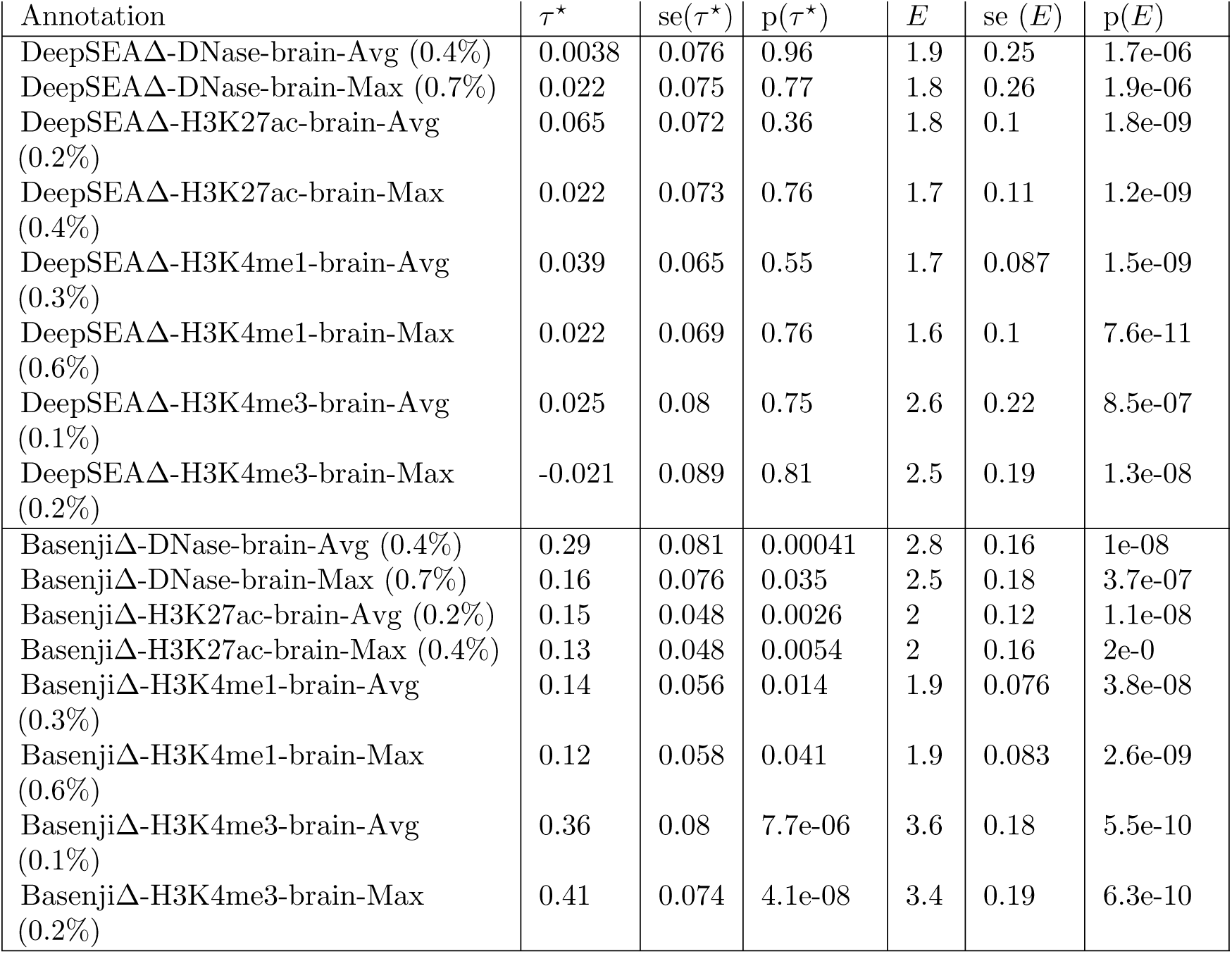
**S-LDSC results for marginal analysis of brain-specific allelic-effect deep learning annotations conditioned on the brain-specific variant-level joint model**. Standardized Effect sizes (*τ\**) and Enrichment (E) of 8 brain-specific DeepSEAΔ and 8 brain-specific BasenjiΔ annotations, when conditioned on the brain-specific variant-level joint model (baseline-LD model + 7 non-tissue-specific annotations from Figure S6 + DeepSEAV-H3K4me3-brain-Max and BasenjiV-H3K27ac-brain-Max (the 2 significant brain-specific variant-level annotations; Figure S12) + 4 additional brain-specific annotations from Figure S15). Results are meta-analyzed across 8 brain-related traits.

**Table S24.**
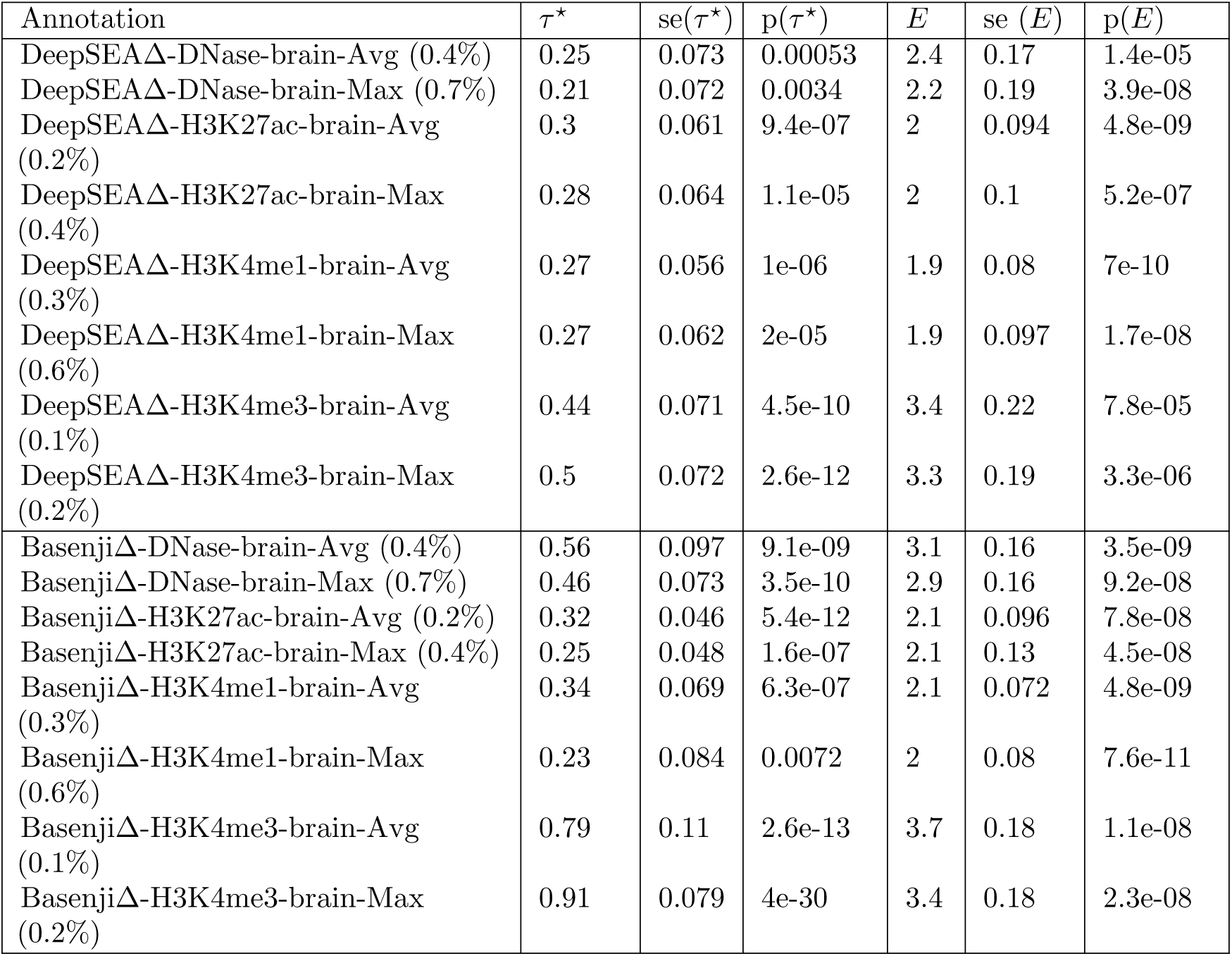
**S-LDSC results for marginal analysis of brain-specific allelic-effect deep learning annotations conditioned on the non-tissue-specific final joint model**. Standardized Effect sizes (*τ\**) and Enrichment (E) of 8 brain-specific DeepSEAΔ and 8 brain-specific BasenjiΔ annotations, when conditioned on the non-tissue-specific final joint model. Results are meta-analyzed across 8 brain-related traits.

**Table S25.** **S-LDSC results for marginal analysis of brain-specific Basenji variant-level annotations and allelic-effect annotations conditioned on either the brain-specific variant-level joint model and BasenjiΔ-H3K4me3-Max or the brain-specific final joint model. We report the standardized effect** sizes (*τ^∗^*) and enrichment (E) of Basenji-H3K4me3-brain-Avg, Basenji-H3K4me3-brain-Max and BasenjiΔ-H3K4me3-brain-Avg, when conditioned on either the brain-specific variant-level joint model (baseline-LD model + 7 non-tissue-specific annotations from Figure S6 + DeepSEAV-H3K4me3-brain-Max and BasenjiV-H3K27ac-brain-Max (the 2 significant brain-specific variant-level annotations; Figure S12) + 4 additional brain-specific annotations from Figure S15) and BasenjiΔ-H3K4me3-Max (top panel) or the brain-specific final joint model (bottom panel). Results are meta-analyzed across 8 brain-related traits.

| Conditional on brain specific variant-level joint model and BasenjiΔ-H3K4me3-Max | | | | | | |
| --- | --- | --- | --- | --- | --- | --- |
| Annotation | $\tau^*$ | se($\tau^*$) | p($\tau^*$) | $E$ | se ($E$) | p($E$) |
| Basenji-H3K4me3-brain-Avg (1.9%) | -0.21 | 0.097 | 0.02 | 3.9 | 0.29 | 4.3e-08 |
| Basenji-H3K4me3-brain-Max (3.4%) | 0.16 | 0.11 | 0.16 | 3.6 | 0.19 | 2.4e-08 |
| BasenjiΔ-H3K4me3-brain-Avg (0.1%) | 0.34 | 0.097 | 0.00047 | 3.5 | 0.18 | 2.5e-08 |
| BasenjiΔ-H3K4me3-brain-Max (0.2%) | 0.46 | 0.095 | 1.4e-06 | 3.4 | 0.19 | 3.2e-08 |
| Conditional on brain-specific final joint model |  |  |  |  |  |  |
| Annotation | $\tau^*$ | se($\tau^*$) | p($\tau^*$) | $E$ | se ($E$) | p($E$) |
| Basenji-H3K4me3-brain-Avg (1.9%) | -0.21 | 0.098 | 0.02 | 3.6 | 0.3 | 3e-10 |
| Basenji-H3K4me3-brain-Max (3.4%) | 0.049 | 0.11 | 0.65 | 3.4 | 0.2 | 1.6e-10 |
| BasenjiΔ-H3K4me3-brain-Avg (0.1%) | -0.62 | 0.33 | 0.061 | 3.3 | 0.18 | 1.3e-08 |
| BasenjiΔ-H3K4me3-brain-Max (0.2%) | 0.45 | 0.13 | 5e-04 | 3.3 | 0.18 | 2.7e-08 |

**Table S26.** **S-LDSC results for marginal analysis of brain-specific allelic-effect deep learning annotations conditioned on the joint model of brain-specific Roadmap and ChromHMM annotations but not the variant-level deep learning annotations**. Standardized Effect sizes (*τ\**) and Enrichment (E) of 8 brain-specific DeepSEAΔ and 8 brain-specific BasenjiΔ annotations, when conditioned on the joint model in Table S47) but without the variant-level deep learning terms. Results are meta-analyzed across 8 brain-related traits.

| Annotation | $\tau^*$ | $se(\tau^*)$ | $p(\tau^*)$ | $E$ | $se(E)$ | $p(E)$ |
| --- | --- | --- | --- | --- | --- | --- |
| DeepSEAΔ-DNase-brain-Avg (0.4%) | 0.22 | 0.074 | 0.0024 | 2.3 | 0.2 | 1.8e-05 |
| DeepSEAΔ-DNase-brain-Max (0.7%) | 0.21 | 0.073 | 0.0032 | 2.2 | 0.22 | 0.00014 |
| DeepSEAΔ-H3K27ac-brain-Avg (0.2%) | 0.26 | 0.066 | 8.5e-05 | 2 | 0.1 | 1.5e-07 |
| DeepSEAΔ-H3K27ac-brain-Max (0.4%) | 0.24 | 0.067 | 0.0003 | 2 | 0.11 | 1.7e-06 |
| DeepSEAΔ-H3K4me1-brain-Avg (0.3%) | 0.21 | 0.061 | 0.00068 | 1.8 | 0.085 | 2.6e-08 |
| DeepSEAΔ-H3K4me1-brain-Max (0.6%) | 0.21 | 0.065 | 0.0011 | 1.8 | 0.1 | 3.3e-06 |
| DeepSEAΔ-H3K4me3-brain-Avg (0.1%) | 0.28 | 0.07 | 4.6e-05 | 3.1 | 0.22 | 4.8e-08 |
| DeepSEAΔ-H3K4me3-brain-Max (0.2%) | 0.31 | 0.072 | 1.4e-05 | 3 | 0.18 | 9.9e-09 |
| BasenjiΔ-DNase-brain-Avg (0.4%) | 0.43 | 0.077 | 2.9e-08 | 3 | 0.16 | 1.5e-10 |
| BasenjiΔ-DNase-brain-Max (0.7%) | 0.32 | 0.074 | 1.8e-05 | 2.7 | 0.17 | 3.6e-08 |
| BasenjiΔ-H3K27ac-brain-Avg (0.2%) | 0.17 | 0.048 | 0.00031 | 2 | 0.12 | 1.2e-08 |
| BasenjiΔ-H3K27ac-brain-Max (0.4%) | 0.14 | 0.048 | 0.0031 | 2 | 0.16 | 1.4e-07 |
| BasenjiΔ-H3K4me1-brain-Avg (0.3%) | 0.23 | 0.056 | 3.6e-05 | 1.9 | 0.076 | 7.1e-09 |
| BasenjiΔ-H3K4me1-brain-Max (0.6%) | 0.17 | 0.058 | 0.0038 | 1.9 | 0.083 | 1.4e-09 |
| BasenjiΔ-H3K4me3-brain-Avg (0.1%) | 0.44 | 0.077 | 1.3e-08 | 3.5 | 0.19 | 4.7e-10 |
| BasenjiΔ-H3K4me3-brain-Max (0.2%) | 0.48 | 0.072 | 4.6e-11 | 3.3 | 0.2 | 1.8e-10 |

**Table S27.** **S-LDSC results for brain-specific final joint model**. The results are conditional on the brain-specific variant-level joint model and 1 significant non-tissue-specific allelic-effect annotation. Results are meta-analyzed across 8 brain-related traits

| Annotation | $\tau^*$ | $se(\tau^*)$ | $p(\tau^*)$ | $E$ | $se(E)$ | $p(E)$ |
| --- | --- | --- | --- | --- | --- | --- |
| BasenjiΔ-H3K4me3-brain-Avg (0.1%) | -0.62 | 0.33 | 0.06 | 3.3 | 0.18 | 1.3e-08 |
| BasenjiΔ-H3K4me3-brain-Max (0.2%) | 0.45 | 0.13 | 5e-04 | 3.3 | 0.18 | 2.7e-08 |

**Table S28.** **Number of significantly disease informative non-tissue-specific variant-level DeepSEAV and BasenjiV annotations**. Number of significantly disease informative non-tissue-specific variant-level annotations across different aggregation strategies (A) different deep learning models (DeepSEA and Basenji), (B) different aggregation strategies (Average and Maximum) and (C) different types of epigenomic marks, in terms of marginal enrichment in heritability, conditional *τ\** and joint *τ^*^*for all traits, blood traits and brain traits respectively.

| **PANEL A** |  |  |
| --- | --- | --- |
| Enrichment |  |  |
| Model | Aggr. Strategy | Number of significant annotations |
| DeepSEAV | All tissues. All traits | 8 |
|  | Brain tissues, Brain traits | 8 |
|  | Blood cell types, Blood traits | 8 |
| BasenjiV | All tissues. All traits | 8 |
|  | Brain tissues, Brain traits | 8 |
|  | Blood cell types, Blood traits | 8 |
| Marginal $\tau^*$ | | |
| Model | Aggregation Strategy | Number of significant annotations |
| DeepSEAΔ | All tissues. All traits | 1 |
|  | Brain tissues, Brain traits | 8 |
|  | Blood cell types, Blood traits | 0 |
| BasenjiΔ | All tissues. All traits | 0 |
|  | Brain tissues, Brain traits | 8 |
|  | Blood cell types, Blood traits | 4 |
| Joint $\tau^*$ | | |
| Model | Aggregation Strategy | Number of significant annotations |
| DeepSEAΔ | All tissues. All traits | 0 |
|  | Brain tissues, Brain traits | 1 |
|  | Blood cell types, Blood traits | 0 |
| BasenjiΔ | All tissues. All traits | 0 |
|  | Brain tissues, Brain traits | 1 |
|  | Blood cell types, Blood traits | 0 |
| **PANEL B** |  |  |
| Enrichment |  |  |
| Model | Aggr. Strategy | Number of significant annotations |
| Avg | All tissues. All traits | 8 |
|  | Brain tissues, Brain traits | 8 |
|  | Blood cell types, Blood traits | 8 |
| Max | All tissues. All traits | 8 |
|  | Brain tissues, Brain traits | 8 |
|  | Blood cell types, Blood traits | 8 |
| Marginal $\tau^*$ | | |

| Model | Aggregation Strategy | Number of significant annotations |
| --- | --- | --- |
| Avg | All tissues. All traits | 0 |
|  | Brain tissues, Brain traits | 8 |
|  | Blood cell types, Blood traits | 2 |
| Max | All tissues. All traits | 1 |
|  | Brain tissues, Brain traits | 8 |
|  | Blood cell types, Blood traits | 2 |

Joint $\tau^*$
| Model | Aggregation Strategy | Number of significant annotations |
| --- | --- | --- |
| Avg | All tissues. All traits | 0 |
|  | Brain tissues, Brain traits | 0 |
|  | Blood cell types, Blood traits | 0 |
| Max | All tissues. All traits | 0 |
|  | Brain tissues, Brain traits | 2 |
|  | Blood cell types, Blood traits | 0 |

Marginal $\tau^*$
| Model | Aggregation Strategy | Number of significant annotations |
| --- | --- | --- |
| DNase | All tissues. All traits | 0 |
|  | Brain tissues, Brain traits | 4 |
|  | Blood cell types, Blood traits | 0 |
| H3K27ac | All tissues. All traits | 0 |
|  | Brain tissues, Brain traits | 4 |
|  | Blood cell types, Blood traits | 1 |
| H3K4me1 | All tissues. All traits | 0 |
|  | Brain tissues, Brain traits | 4 |
|  | Blood cell types, Blood traits | 2 |
| H3K4me3 | All tissues. All traits | 1 |
|  | Brain tissues, Brain traits | 4 |

|  | Blood cell types, Blood traits | 1 |
| --- | --- | --- |
| Joint $\tau^*$ | | |
| Model | Aggregation Strategy | Number of significant annotations |
| DNase | All tissues. All traits | 0 |
|  | Brain tissues, Brain traits | 0 |
|  | Blood cell types, Blood traits | 0 |
| H3K27ac | All tissues. All traits | 0 |
|  | Brain tissues, Brain traits | 1 |
|  | Blood cell types, Blood traits | 0 |
| H3K4me1 | All tissues. All traits | 0 |
|  | Brain tissues, Brain traits | 0 |
|  | Blood cell types, Blood traits | 0 |
| H3K4me3 | All tissues. All traits | 0 |
|  | Brain tissues, Brain traits | 1 |
|  | Blood cell types, Blood traits | 0 |

**Table S29.** **S-LDSC results for marginal analysis of non-tissue-specific variant-level DeepSEAV and BasenjiV annotations**. Standardized Effect sizes (*τ\**) and Enrichment (E) of 8 non-tissue-specific variant-level DeepSEAV and 8 non-tissue-specific variant-level BasenjiV annotations, when conditioned on 86 baseline-LD annotations. Results are meta-analyzed across 41 traits.

| Annotation | $\tau^*$ | $se(\tau^*)$ | $p(\tau^*)$ | $E$ | $se(E)$ | $p(E)$ |
| --- | --- | --- | --- | --- | --- | --- |
| BasenjiV-DNase-Avg (1.7%) | -0.078 | 0.07 | 0.26 | 2.4 | 0.2 | 4.9e-09 |
| BasenjiV-DNase-Max (13.6%) | -0.095 | 0.065 | 0.14 | 1.8 | 0.079 | 1.7e-16 |
| BasenjiV-H3K27ac-Avg (3.4%) | 0.14 | 0.059 | 0.016 | 2.5 | 0.14 | 2.7e-16 |
| BasenjiV-H3K27ac-Max (14.2%) | -0.026 | 0.031 | 0.41 | 1.8 | 0.074 | 9.4e-16 |
| BasenjiV-H3K4me1-Avg (5.6%) | 0.08 | 0.042 | 0.055 | 2 | 0.076 | 2.7e-17 |
| BasenjiV-H3K4me1-Max (23.7%) | -0.057 | 0.027 | 0.032 | 1.6 | 0.046 | 1.4e-17 |
| BasenjiV-H3K4me3-Avg (2.0%) | 0.17 | 0.07 | 0.014 | 3.7 | 0.26 | 1.4e-16 |
| BasenjiV-H3K4me3-Max (9.4%) | 0.079 | 0.036 | 0.029 | 2 | 0.088 | 5.3e-15 |
| DeepSEAV-DNase-Avg (1.7%) | -0.12 | 0.11 | 0.27 | 2.3 | 0.2 | 3e-07 |
| DeepSEAV-DNase-Max (13.6%) | -0.17 | 0.078 | 0.028 | 1.8 | 0.075 | 1.1e-15 |
| DeepSEAV-H3K27ac-Avg (3.4%) | 0.19 | 0.071 | 0.0089 | 2.4 | 0.12 | 1.1e-17 |
| DeepSEAV-H3K27ac-Max (14.2%) | 0.0092 | 0.06 | 0.88 | 1.8 | 0.066 | 2.6e-17 |
| DeepSEAV-H3K4me1-Avg (5.6%) | 0.015 | 0.059 | 0.8 | 1.9 | 0.059 | 2.2e-17 |
| DeepSEAV-H3K4me1-Max (23.7%) | -0.13 | 0.06 | 0.027 | 1.6 | 0.041 | 8.8e-18 |
| DeepSEAV-H3K4me3-Avg (2.0%) | 0.27 | 0.093 | 0.0033 | 4 | 0.25 | 7.1e-19 |
| DeepSEAV-H3K4me3-Max (9.4%) | 0.25 | 0.06 | 4.5e-05 | 2.2 | 0.087 | 2.2e-17 |

**Table S30.**
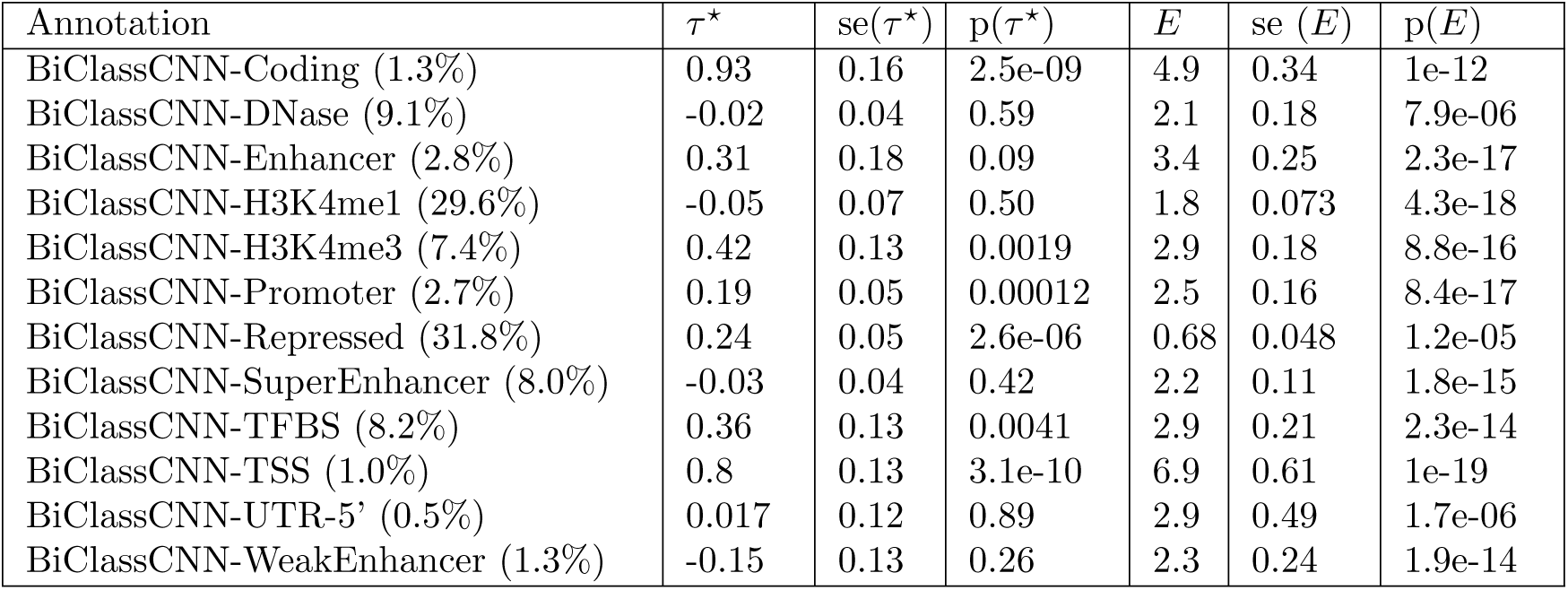
**S-LDSC results for marginal analysis of non-tissue-specific Bi-ClassCNN annotations**. Standardized Effect sizes (*τ\**) and Enrichment (E) of 12 non-tissue-specific BiClassCNN annotations. Results are meta-analyzed across 41 traits.

| Annotation | $\tau^*$ | $se(\tau^*)$ | $p(\tau^*)$ | $E$ | $se(E)$ | $p(E)$ |
| --- | --- | --- | --- | --- | --- | --- |
| BiClassCNN-Coding (1.3%) | 0.93 | 0.16 | 2.5e-09 | 4.9 | 0.34 | 1e-12 |
| BiClassCNN-DNase (9.1%) | -0.02 | 0.04 | 0.59 | 2.1 | 0.18 | 7.9e-06 |
| BiClassCNN-Enhancer (2.8%) | 0.31 | 0.18 | 0.09 | 3.4 | 0.25 | 2.3e-17 |
| BiClassCNN-H3K4me1 (29.6%) | -0.05 | 0.07 | 0.50 | 1.8 | 0.073 | 4.3e-18 |
| BiClassCNN-H3K4me3 (7.4%) | 0.42 | 0.13 | 0.0019 | 2.9 | 0.18 | 8.8e-16 |
| BiClassCNN-Promoter (2.7%) | 0.19 | 0.05 | 0.00012 | 2.5 | 0.16 | 8.4e-17 |
| BiClassCNN-Repressed (31.8%) | 0.24 | 0.05 | 2.6e-06 | 0.68 | 0.048 | 1.2e-05 |
| BiClassCNN-SuperEnhancer (8.0%) | -0.03 | 0.04 | 0.42 | 2.2 | 0.11 | 1.8e-15 |
| BiClassCNN-TFBS (8.2%) | 0.36 | 0.13 | 0.0041 | 2.9 | 0.21 | 2.3e-14 |
| BiClassCNN-TSS (1.0%) | 0.8 | 0.13 | 3.1e-10 | 6.9 | 0.61 | 1e-19 |
| BiClassCNN-UTR-5' (0.5%) | 0.017 | 0.12 | 0.89 | 2.9 | 0.49 | 1.7e-06 |
| BiClassCNN-WeakEnhancer (1.3%) | -0.15 | 0.13 | 0.26 | 2.3 | 0.24 | 1.9e-14 |

**Table S31.** **. BiClassCNN-Coding, BiClassCNN-Repressed and BiClassCNN-TSS produced independent signals.**. We report standardized effect sizes (*τ\**) and enrichment (E) of the 4 significant BiClassCNN annotations from Figure 1, when modeled jointly and passed through the forward stepwise elimination, all conditioned on the baseline-LD model. The three annotations that remain are BiClassCNN-Coding, BiClassCNN-Repressed and BiClassCNN-TSS. Results are meta-analyzed across 41 traits.

| Annotation | $\tau^*$ | $\text{se}(\tau^*)$ | $\text{p}(\tau^*)$ | $E$ | $\text{se}(E)$ | $\text{p}(E)$ |
| --- | --- | --- | --- | --- | --- | --- |
| BiClassCNN-Coding (1.3%) | 0.73 | 0.17 | 2.2e-05 | 4.9 | 0.34 | 1.4e-15 |
| BiClassCNN-Repressed (31.8%) | 0.19 | 0.051 | 0.00018 | 0.68 | 0.048 | 2.9e-07 |
| BiClassCNN-TSS (1.0%) | 0.68 | 0.13 | 9.6e-08 | 6.8 | 0.6 | 0.00023 |

**Table S32.** **S-LDSC results for marginal analysis of non-tissue-specific Roadmap and ChromHMM annotations**. Standardized Effect sizes (*τ\**) and Enrichment (E) of 8 non-tissue-specific Roadmap and 40 non-tissue-specific ChromHMM annotations. Results are meta-analyzed across 41 traits.

| Annotation | $\tau^*$ | $se(\tau^*)$ | $p(\tau^*)$ | $E$ | $se(E)$ | $p(E)$ |
| --- | --- | --- | --- | --- | --- | --- |
| Roadmap-DNase-Avg (2.2%) | 0.39 | 0.11 | 0.00034 | 5.3 | 0.46 | 9.7e-13 |
| Roadmap-DNase-Max (13.4%) | 0.18 | 0.075 | 0.02 | 3 | 0.23 | 3.9e-12 |
| Roadmap-H3K27ac-Avg (3.1%) | 0.37 | 0.084 | 1.1e-05 | 4.2 | 0.27 | 9.4e-17 |
| Roadmap-H3K27ac-Max (21.4%) | 0.16 | 0.079 | 0.046 | 2.5 | 0.13 | 6.4e-17 |
| Roadmap-H3K4me1-Avg (4.4%) | 0.47 | 0.077 | 9.1e-10 | 3.6 | 0.19 | 1.9e-17 |
| Roadmap-H3K4me1-Max (25.4%) | 0.094 | 0.099 | 0.34 | 2.2 | 0.1 | 8.1e-19 |
| Roadmap-H3K4me3-Avg (1.7%) | 0.43 | 0.088 | 7.2e-07 | 5.8 | 0.41 | 2.7e-18 |
| Roadmap-H3K4me3-Max (6.6%) | -0.12 | 0.065 | 0.075 | 3.3 | 0.23 | 5.7e-16 |
| ChromHMM-DNase-Avg (0.7%) | -0.12 | 0.049 | 0.012 | 1.4 | 0.39 | 0.4 |
| ChromHMM-DNase-Max (9.2%) | -0.19 | 0.061 | 0.0021 | 1.5 | 0.24 | 0.71 |
| ChromHMM-Active-Enhancer-1-Avg (0.3%) | -0.15 | 0.03 | 3.2e-07 | 2.7 | 0.3 | 1.2e-08 |
| ChromHMM-Active-Enhancer-1-Max (7.1%) | -0.17 | 0.044 | 8.1e-05 | 2.3 | 0.22 | 0.0025 |
| ChromHMM-Active-Enhancer-2-Avg (0.4%) | -0.073 | 0.052 | 0.16 | 2.9 | 0.34 | 0.0011 |
| ChromHMM-Active-Enhancer-2-Max (9.9%) | -0.13 | 0.057 | 0.027 | 2.2 | 0.2 | 0.011 |
| ChromHMM-Enhancer-acetylation-Avg (0.3%) | -0.19 | 0.039 | 8.5e-07 | 1.4 | 0.25 | 0.79 |
| ChromHMM-Enhancer-acetylation-Max (11.3%) | -0.19 | 0.029 | 1.8e-10 | 1.5 | 0.13 | 0.0066 |
| ChromHMM-Active-Enhancer-Flanking-Avg (0.5%) | -0.12 | 0.035 | 0.00085 | 2.3 | 0.22 | 0.0092 |
| ChromHMM-Active-Enhancer-Flanking-Max (11.1%) | -0.15 | 0.046 | 0.00078 | 1.9 | 0.15 | 0.11 |
| ChromHMM-Weak-Enhancer-1-Avg (0.3%) | -0.14 | 0.035 | 7.1e-05 | 1.8 | 0.31 | 9.9e-08 |
| ChromHMM-Weak-Enhancer-1-Max (7.7%) | -0.21 | 0.034 | 8.8e-10 | 1.8 | 0.19 | 5.1e-06 |
| ChromHMM-Weak-Enhancer-2-Avg (1.2%) | -0.052 | 0.036 | 0.15 | 2.1 | 0.14 | 1.5e-10 |
| ChromHMM-Weak-Enhancer-2-Max (19.9%) | -0.12 | 0.052 | 0.02 | 1.7 | 0.093 | 5.8e-11 |
| ChromHMM-Heterochromatin-Avg (1.4%) | -0.074 | 0.02 | 0.00027 | 0.3 | 0.058 | 2e-18 |
| ChromHMM-Heterochromatin-Max (7.1%) | -0.055 | 0.018 | 0.0022 | 0.39 | 0.038 | 3.5e-21 |
| ChromHMM-Promoter-Bivalent-Avg (0.3%) | 0.1 | 0.042 | 0.017 | 6.1 | 0.48 | 8.2e-14 |
| ChromHMM-Promoter-Bivalent-Max (1.9%) | 0.055 | 0.041 | 0.18 | 4 | 0.27 | 2.1e-14 |
| ChromHMM-Promoter-Downstream-1-Avg (0.4%) | 0.18 | 0.062 | 0.0029 | 7 | 0.89 | 7.8e-12 |
| ChromHMM-Promoter-Downstream-1-Max (1.4%) | 0.24 | 0.063 | 0.0002 | 6.5 | 0.6 | 8.9e-16 |
| ChromHMM-Promoter-Downstream-2-Avg (0.2%) | -0.014 | 0.045 | 0.76 | 2.5 | 0.67 | 0.1 |
| ChromHMM-Promoter-Downstream-2-Max (1.5%) | 0.095 | 0.033 | 0.004 | 3.6 | 0.34 | 2.2e-14 |
| ChromHMM-Promoter-Poised-Avg (0.2%) | -0.14 | 0.046 | 0.0018 | 1.3 | 0.25 | 0.074 |
| ChromHMM-Promoter-Poised-Max (6.8%) | -0.23 | 0.037 | 1.1e-09 | 1.2 | 0.098 | 0.0011 |

Table S32. (continued)
| Annotation | $\tau^*$ | $\text{se}(\tau^*)$ | $p(\tau^*)$ | $E$ | $\text{se}(E)$ | $p(E)$ |
| --- | --- | --- | --- | --- | --- | --- |
| ChromHMM-Promoter_Upstream-Avg (0.4%) | 0.04 | 0.051 | 0.43 | 5.7 | 0.45 | 5.3e-16 |
| ChromHMM-Promoter_Upstream-Max (2.5%) | -0.13 | 0.045 | 0.0052 | 3.8 | 0.34 | 7.5e-14 |
| ChromHMM-Quiescent-Avg (78.4%) | -0.061 | 0.018 | 0.00074 | 0.73 | 0.02 | 3e-18 |
| ChromHMM-Quiescent-Max (91.0%) | -0.086 | 0.014 | 1.4e-09 | 0.86 | 0.01 | 6.6e-20 |
| ChromHMM-Repressed_Polycomb-Avg (1.7%) | -0.052 | 0.016 | 0.0011 | 1.5 | 0.065 | 1.5e-16 |
| ChromHMM-Repressed_Polycomb-Max (9.9%) | -0.072 | 0.016 | 6.5e-06 | 1.3 | 0.037 | 2.9e-20 |
| ChromHMM-TSS-Avg (0.2%) | -0.0011 | 0.075 | 0.99 | 5.1 | 1 | 5.1e-05 |
| ChromHMM-TSS-Max (0.9%) | -0.058 | 0.073 | 0.42 | 4.5 | 0.49 | 1.7e-10 |
| ChromHMM-Transcription_Enhancer_3'-Avg (0.2%) | 0.078 | 0.027 | 0.0038 | 3.4 | 0.29 | 1.7e-16 |
| ChromHMM-Transcription_Enhancer_3'-Max (3.1%) | 0.088 | 0.035 | 0.013 | 2.8 | 0.21 | 6.7e-18 |
| ChromHMM-Transcription_Enhancer-5'-Avg (0.4%) | 0.12 | 0.024 | 9.7e-07 | 3.6 | 0.23 | 5.4e-18 |
| ChromHMM-Transcription_Enhancer-5'-Max (4.8%) | 0.16 | 0.022 | 2.2e-13 | 2.9 | 0.15 | 6.5e-19 |
| ChromHMM-Transcription_Weak_Enhancer-Avg (0.5%) | 0.12 | 0.024 | 1.3e-06 | 2.8 | 0.16 | 7.3e-19 |
| ChromHMM-Transcription_Weak_Enhancer-Max (4.7%) | 0.16 | 0.022 | 3e-12 | 2.6 | 0.12 | 1.7e-21 |
| ChromHMM-Transcriptional_Regulator-Avg (0.3%) | 0.27 | 0.039 | 5.2e-12 | 6.8 | 0.51 | 4.3e-17 |
| ChromHMM-Transcriptional_Regulator-Max (3.3%) | 0.22 | 0.037 | 4.2e-09 | 4 | 0.26 | 1.8e-18 |

**Table S33.** **S-LDSC results for marginal analysis of CpG-island, local CpG-content, local GC-content, pLI, SIFT and Polyphen annotations**. Standardized Effect sizes (*τ\**) and Enrichment (E) of the weighted Coding, TSS, Repressed and Gene ±5*KB* annotations, where the weights may be driven by the underlying sequence context such as CpG-island, LocalCpG-content (±1*kb*), LocalGC-content (±1*kb*) etc as well as pLI, pLI-Coding, pLI-TSS, PolyPhen and SIFT. All analyses are conditioned on baseline-LD annotations. Results are meta-analyzed across 41 traits.

| Annotation | $\tau^*$ | se($\tau^*$) | p($\tau^*$) | $E$ | se ($E$) | p($E$) |
| --- | --- | --- | --- | --- | --- | --- |
| CpGisland (0.8%) | 0.2 | 0.051 | 6.1e-05 | 6.7 | 0.61 | 5e-14 |
| CpGisland-Coding (0.2%) | 0.094 | 0.049 | 0.056 | 7.7 | 1.2 | 1.6e-11 |
| CpGisland-Repressed (0.02%) | -0.074 | 0.023 | 0.001 | 3.9 | 1.4 | 0.19 |
| CpGisland-TSS (0.4%) | 0.23 | 0.074 | 0.0019 | 8.8 | 0.98 | 2.2e-11 |
| LocalCpGcontent (5.5%) | -0.19 | 0.052 | 0.00025 | 1.1 | 0.013 | 1.5e-15 |
| LocalCpGcontent-Coding (0.2%) | 0.44 | 0.075 | 4.9e-09 | 6 | 0.45 | 7.6e-16 |
| LocalCpGcontent-Repressed (2.2%) | -0.34 | 0.058 | 4.9e-09 | 0.59 | 0.055 | 3.7e-10 |
| LocalCpGcontent-TSS (0.2%) | 0.66 | 0.12 | 3.8e-08 | 7.1 | 0.59 | 1.1e-17 |
| LocalGCcontent (41.0%) | -0.15 | 0.03 | 5.7e-07 | 1.1 | 0.0062 | 1.1e-11 |
| LocalGCcontent-Coding (0.8%) | 1.1 | 0.17 | 1.9e-10 | 5 | 0.35 | 2e-16 |
| LocalGCcontent-Repressed (18.0%) | -0.68 | 0.11 | 5.3e-10 | 0.63 | 0.05 | 1.1e-08 |
| LocalGCcontent-TSS (1.0%) | 1.8 | 0.28 | 1.7e-10 | 6.4 | 0.56 | 5.7e-19 |
| pLI (14.0%) | 0.095 | 0.011 | 3.9e-18 | 1.5 | 0.034 | 9.5e-17 |
| pLI-Coding (0.4%) | 0.62 | 0.057 | 2.5e-27 | 13 | 0.78 | 5.6e-17 |
| pLI-TSS (0.3%) | 0.52 | 0.039 | 3.5e-41 | 14 | 1.1 | 6.4e-20 |
| Polyphen (0.1%) | 0.23 | 0.081 | 0.0046 | 13 | 2.2 | 1.8e-10 |
| SIFT (0.1%) | -0.067 | 0.065 | 0.3 | 4.6 | 1.5 | 4e-06 |

**Table S34.** **S-LDSC results for non-tissue-specific variant-level joint model**. Joint S-LDSC model fit of all non-tissue-specific annotations as well as LocalGC-content, LocalCpG-content and CpG-island weighted annotations of TSS, Repressed and Coding regions, and pLI, pLI-TSS, pLI-Coding, SIFT and Polyphen. Only 7 annotations were found to be significant conditioned on all other annotations and the baseline-LD model. Results are meta-analyzed across 41 traits. We also highlight in red annotations from Figure 1 that were not significant under the joint model

| Annotation | $\tau^*$ | $se(\tau^*)$ | $p(\tau^*)$ | $E$ | $se(E)$ | $p(E)$ |
| --- | --- | --- | --- | --- | --- | --- |
| ChromHMM-Active-Enhancer-1-Avg (0.3%) | -0.16 | 0.03 | 6.8e-08 | 2.6 | 0.29 | 1.8e-10 |
| Roadmap-H3K4me1-Avg (4.4%) | 0.46 | 0.079 | 5e-09 | 3.6 | 0.19 | 6.5e-21 |
| pLI-Coding (0.4%) | 0.45 | 0.054 | 4e-17 | 11 | 0.71 | 3e-16 |
| pLI-TSS (0.3%) | 0.35 | 0.039 | 1.4e-19 | 12 | 1 | 8.2e-21 |
| LocalGCcontent (41.0%) | -0.16 | 0.03 | 1.1e-07 | 1.1 | 0.0066 | 4.2e-17 |
| LocalGCcontent-TSS (1.0%) | 1.2 | 0.28 | 1.6e-05 | 5.8 | 0.52 | 5.8e-26 |
| LocalGCcontent-Coding (0.8%) | 0.91 | 0.18 | 7.8e-07 | 5.2 | 0.34 | 6.7e-20 |
| DeepSEAV-H3K4me3-Max (9.4%) | 0.13 | 0.06 | 0.029 | 2.2 | 0.077 | 6.8e-19 |
| BiClassCNN-Coding (1.3%) | 0.43 | 0.28 | 0.13 | 5.1 | 0.34 | 6.5e-19 |
| BiClassCNN-Promoter (2.0%) | 0.14 | 0.048 | 0.0036 | 2.7 | 0.17 | 2.2e-20 |
| BiClassCNN-Repressed (31.8%) | 0.1 | 0.043 | 0.02 | 0.66 | 0.046 | 1.1e-08 |
| BiClassCNN-TSS (1.0%) | 0.19 | 0.25 | 0.44 | 6.2 | 0.55 | 3.9e-18 |

**Table S35.** **The Roadmap-H3K4me1-Avg annotation was sufficient to eliminate the DeepSEAV-H3K4me3-Max annotation that was significant in marginal analysis**. Standardized Effect sizes (*τ\**) and Enrichment (E) of the significant non-tissue-specific variant-level DeepSEAV and Roadmap annotations from marginal analysis, when modeled conditioned on each other and the 86 baseline-LD annotations, and passed through Forward Stepwise Elimination. The results for the penultimate step of the elimination process is shown. DeepSEAV-H3K4me3-Max is not significant after correcting for all tested hypothesis (0.05*/*106) and gets eliminated next. Results are meta-analyzed across 41 traits.

| Annotation | $\tau^*$ | $se(\tau^*)$ | $p(\tau^*)$ | $E$ | $se(E)$ | $p(E)$ |
| --- | --- | --- | --- | --- | --- | --- |
| Roadmap-H3K4me1-Avg (4.4%) | 0.455 | 0.08 | 1e-08 | 3.63 | 0.19 | 3e-19 |
| DeepSEAV-H3K4me3-Max (9.4%) | 0.182 | 0.06 | 0.004 | 2.14 | 0.08 | 1e-16 |

**Table S36.** **The Local GC-content annotations were sufficient to eliminate the 4 BiClassCNN annotations that were significant in our marginal analysis**. Meta-analyzed *τ\** and Enrichment *E* of the Joint S-LDSC model fit of the 4 BiClassCNN annotations from Figure 1 conditioned on the 86 baseline-LD annotations and 3 LocalGCcontent annotations from Table S34. Results are meta-analyzed across 41 traits.

| Annotation | $\tau^*$ | $\text{se}(\tau^*)$ | $\text{p}(\tau^*)$ | $E$ | $\text{se}(E)$ | $\text{p}(E)$ |
| --- | --- | --- | --- | --- | --- | --- |
| BiClassCNN-Coding (1.3%) | 0.69 | 0.3 | 0.023 | 5 | 0.35 | 1e-16 |
| BiClassCNN-Promoter (2.7%) | 0.15 | 0.052 | 0.0039 | 2.7 | 0.17 | 3.1e-20 |
| BiClassCNN-Repressed (31.8%) | 0.1 | 0.045 | 0.022 | 0.69 | 0.047 | 1.9e-07 |
| BiClassCNN-TSS (1.0%) | 0.13 | 0.28 | 0.65 | 6.6 | 0.58 | 3e-19 |

**Table S37.**
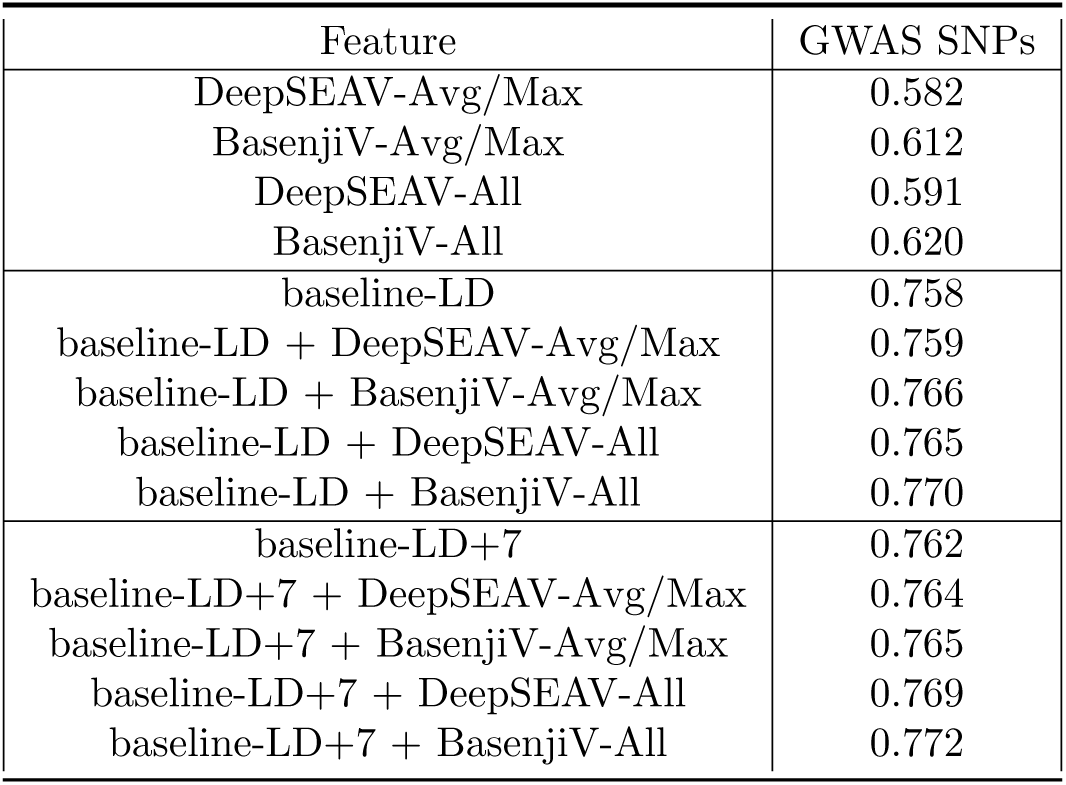
**AUROC of various sets of annotations incorporating non-tissue-specific variant-level deep learning annotations in predicting** 12, 296 **NIH GWAS SNPs**. We report the AUROC for a gradient boosting model trained using each respective set of annotations, either without including baseline-LD model annotations; including baseline-LD model annotations; or including annotations from the non-tissue-specific variant-level joint model (baseline-LD model + 7 annotations; baseline-LD+7).

| Feature | GWAS SNPs |
| --- | --- |
| DeepSEAV-Avg/Max | 0.582 |
| BasenjiV-Avg/Max | 0.612 |
| DeepSEAV-All | 0.591 |
| BasenjiV-All | 0.620 |
| baseline-LD | 0.758 |
| baseline-LD + DeepSEAV-Avg/Max | 0.759 |
| baseline-LD + BasenjiV-Avg/Max | 0.766 |
| baseline-LD + DeepSEAV-All | 0.765 |
| baseline-LD + BasenjiV-All | 0.770 |
| baseline-LD+7 | 0.762 |
| baseline-LD+7 + DeepSEAV-Avg/Max | 0.764 |
| baseline-LD+7 + BasenjiV-Avg/Max | 0.765 |
| baseline-LD+7 + DeepSEAV-All | 0.769 |
| baseline-LD+7 + BasenjiV-All | 0.772 |

**Table S38.** **Enrichment of NIH GWAS SNPs for non-tissue-specific variant-level annotations**. We report the enrichment in deep learning annotations (along with Jackknife standard error) at top 12, 296 known disease-associated SNPs from NIH GWAS catalog^23^.

| Feature | GWAS SNPs |
| --- | --- |
| BasenjiV-DNase-Avg | 1.61 (0.010) |
| BasenjiV-DNase-Max | 1.35 (0.005) |
| BasenjiV-H3K27ac-Avg | 1.62 (0.009) |
| BasenjiV-H3K27ac-Max | 1.42 (0.005) |
| BasenjiV-H3K4me1-Avg | 1.48 (0.005) |
| BasenjiV-H3K4me1-Max | 1.31 (0.003) |
| BasenjiV-H3K4me3-Avg | 1.81 (0.016) |
| BasenjiV-H3K4me3-Max | 1.30 (0.006) |
| DeepSEAV-DNase-Avg | 1.58 (0.009) |
| DeepSEAV-DNase-Max | 1.39 (0.004) |
| DeepSEAV-H3K27ac-Avg | 1.58 (0.008) |
| DeepSEAV-H3K27ac-Max | 1.39 (0.004) |
| DeepSEAV-H3K4me1-Avg | 1.46 (0.001) |
| DeepSEAV-H3K4me1-Max | 1.31 (0.003) |
| DeepSEAV-H3K4me3-Avg | 1.81 (0.013) |
| DeepSEAV-H3K4me3-Max | 1.43 (0.005) |

**Table S39.** **S-LDSC results for marginal analysis of blood-specific variant-level DeepSEAV and BasenjiV annotations**. Standardized Effect sizes (*τ\**) and Enrichment (E) of 8 blood-specific variant-level DeepSEAV and 8 blood-specific variant-level BasenjiV annotations, when conditioned on non-tissue-specific variant-level joint model (baseline-LD and annotations from Figure S6). Results are meta-analyzed across 11 blood-related traits.

| Annotation | $\tau^*$ | $se(\tau^*)$ | $p(\tau^*)$ | $E$ | $se(E)$ | $p(E)$ |
| --- | --- | --- | --- | --- | --- | --- |
| BasenjiV-DNase-blood-Avg (1.9%) | 0.06 | 0.11 | 0.6 | 5 | 0.37 | 1.2e-12 |
| BasenjiV-DNase-blood-Max (5.4%) | 0.22 | 0.1 | 0.028 | 3.5 | 0.2 | 8.8e-06 |
| BasenjiV-H3K27ac-blood-Avg (2.5%) | 0.59 | 0.13 | 4.2e-06 | 4.6 | 0.25 | 3.2e-05 |
| BasenjiV-H3K27ac-blood-Max (8.5%) | 0.38 | 0.13 | 0.0045 | 3.2 | 0.13 | 4.5e-05 |
| BasenjiV-H3K4me1-blood-Avg (6.2%) | 0.77 | 0.13 | 4.4e-09 | 3.2 | 0.15 | 3.7e-05 |
| BasenjiV-H3K4me1-blood-Max (13.7%) | 0.58 | 0.13 | 9.8e-06 | 2.6 | 0.095 | 6.4e-05 |
| BasenjiV-H3K4me3-blood-Avg (2.0%) | 0.34 | 0.13 | 0.011 | 6.9 | 0.41 | 5.8e-05 |
| BasenjiV-H3K4me3-blood-Max (5.5%) | 0.34 | 0.098 | 0.00065 | 4.2 | 0.18 | 6.7e-05 |
| DeepSEAV-DNase-blood-Avg (1.9%) | -0.12 | 0.13 | 0.35 | 4.9 | 0.36 | 6e-05 |
| DeepSEAV-DNase-blood-Max (5.4%) | 0.055 | 0.11 | 0.6 | 3.4 | 0.18 | 6.9e-05 |
| DeepSEAV-H3K27ac-blood-Avg (2.5%) | 0.12 | 0.11 | 0.29 | 4 | 0.21 | 9.1e-05 |
| DeepSEAV-H3K27ac-blood-Max (8.5%) | 0.14 | 0.11 | 0.2 | 2.8 | 0.1 | 9e-05 |
| DeepSEAV-H3K4me1-blood-Avg (6.2%) | 0.14 | 0.086 | 0.093 | 2.8 | 0.11 | 0.0001 |
| DeepSEAV-H3K4me1-blood-Max (13.7%) | 0.14 | 0.073 | 0.048 | 2.4 | 0.074 | 0.0001 |
| DeepSEAV-H3K4me3-blood-Avg (2.0%) | -0.25 | 0.13 | 0.052 | 5.8 | 0.42 | 4.1e-05 |
| DeepSEAV-H3K4me3-blood-Max (5.5%) | 0.064 | 0.096 | 0.5 | 3.9 | 0.19 | 8.3e-05 |

**Table S40.** **S-LDSC results for marginal analysis of blood-specific Roadmap and ChromHMM annotations**. Standardized Effect sizes (*τ\**) and Enrichment (E) of 8 blood-specific Roadmap and 40 blood-specific ChromHMM annotations, when conditioned on non-tissue-specific variant-level joint model (baseline-LD and annotations from Figure S6). Results are meta-analyzed across 11 blood-related traits.

| Annotation | $\tau^*$ | $se(\tau^*)$ | $p(\tau^*)$ | $E$ | $se(E)$ | $p(E)$ |
| --- | --- | --- | --- | --- | --- | --- |
| Roadmap-DNase-blood-Avg (2.1%) | 1.8 | 0.29 | 1.6e-09 | 14 | 1.2 | 7.1e-06 |
| Roadmap-DNase-blood-Max (6.2%) | 1.4 | 0.22 | 9.8e-10 | 8.7 | 0.68 | 1e-05 |
| Roadmap-H3K27ac-blood-Avg (3.2%) | 1.5 | 0.32 | 2e-06 | 9.5 | 0.77 | 4.9e-06 |
| Roadmap-H3K27ac-blood-Max (10.2%) | 1.3 | 0.21 | 1.7e-10 | 6.4 | 0.43 | 9.1e-06 |
| Roadmap-H3K4me1-blood-Avg (4.9%) | 1.5 | 0.33 | 7.3e-06 | 7.1 | 0.54 | 6.4e-06 |
| Roadmap-H3K4me1-blood-Max (11.2%) | 1.4 | 0.24 | 5.7e-09 | 5.8 | 0.42 | 2.8e-06 |
| Roadmap-H3K4me3-blood-Avg (1.8%) | 1.3 | 0.3 | 1.5e-05 | 13 | 1.1 | 1e-05 |
| Roadmap-H3K4me3-blood-Max (3.6%) | 1.4 | 0.23 | 3.6e-09 | 11 | 0.86 | 2.9e-06 |
| ChromHMM-DNase-blood-Avg (0.4%) | -0.017 | 0.11 | 0.88 | 2.9 | 0.91 | 0.0015 |
| ChromHMM-DNase-blood-Max (2.2%) | 0.37 | 0.099 | 0.00019 | 5.2 | 0.59 | 3e-07 |
| ChromHMM-Active_Enhancer_1-blood-Avg (0.3%) | 1 | 0.25 | 3.4e-05 | 12 | 2.2 | 2.1e-05 |
| ChromHMM-Active_Enhancer_1-blood-Max (2.6%) | 1 | 0.17 | 7.7e-09 | 9.3 | 1.1 | 1.7e-07 |
| ChromHMM-Active_Enhancer_2-blood-Avg (0.4%) | 1.1 | 0.26 | 2.1e-05 | 12 | 1.8 | 6.4e-07 |
| ChromHMM-Active_Enhancer_2-blood-Max (3.5%) | 1.2 | 0.22 | 3.3e-08 | 9 | 1.1 | 3.3e-08 |
| ChromHMM-Enhancer_acetylation-blood-Avg (0.2%) | 0.24 | 0.13 | 0.059 | 5 | 0.9 | 4.9e-06 |
| ChromHMM-Enhancer_acetylation-blood-Max (2.8%) | 0.54 | 0.14 | 8.3e-05 | 5.6 | 0.72 | 9.5e-06 |
| ChromHMM-Active_Enhancer_Flanking-blood-Avg (0.4%) | 0.96 | 0.24 | 4.6e-05 | 10 | 1.6 | 1.9e-06 |
| ChromHMM-Active_Enhancer_Flanking-blood-Max (3.3%) | 1.1 | 0.22 | 1.1e-06 | 8.2 | 1.1 | 2.3e-07 |
| ChromHMM-Weak_Enhancer_1-blood-Avg (0.3%) | 0.43 | 0.11 | 5.3e-05 | 8.8 | 1 | 7.4e-08 |
| ChromHMM-Weak_Enhancer_1-blood-Max (2.2%) | 0.78 | 0.14 | 1.4e-08 | 8.6 | 0.89 | 9.7e-06 |
| ChromHMM-Weak_Enhancer_2-blood-Avg (1.1%) | 0.81 | 0.19 | 1.3e-05 | 6.6 | 0.88 | 9e-07 |
| ChromHMM-Weak_Enhancer_2-blood-Max (6.5%) | 1 | 0.19 | 4.1e-08 | 5.8 | 0.62 | 1.6e-07 |
| ChromHMM-Heterochromatin-blood-Avg (1.2%) | -0.13 | 0.031 | 3.2e-05 | 0.12 | 0.16 | 0.00025 |
| ChromHMM-Heterochromatin-blood-Max (3.2%) | -0.097 | 0.031 | 0.002 | 0.01 | 0.13 | 0.00057 |
| ChromHMM-Promoter_Bivalent-blood-Avg (0.3%) | -0.34 | 0.12 | 0.0053 | 2.1 | 1.5 | 0.08 |
| ChromHMM-Promoter_Bivalent-blood-Max (0.8%) | -0.078 | 0.13 | 0.54 | 4.7 | 0.96 | 1.3e-05 |
| ChromHMM-Promoter_Downstream_1-blood-Avg (0.4%) | 0.66 | 0.18 | 0.00034 | 19 | 2.1 | 1.4e-05 |

Table S40. (continued)
| Annotation | $\tau^*$ | $\text{se}(\tau^*)$ | $\text{p}(\tau^*)$ | $E$ | $\text{se}(E)$ | $\text{p}(E)$ |
| --- | --- | --- | --- | --- | --- | --- |
| ChromHMM-Promoter_Downstream_1-blood-Max (0.9%) | 1.1 | 0.18 | 3.2e-10 | 19 | 1.5 | 8.7e-06 |
| ChromHMM-Promoter_Downstream_2-blood-Avg (0.1%) | 0.51 | 0.13 | 0.0001 | 14 | 2.2 | 7.7e-08 |
| ChromHMM-Promoter_Downstream_2-blood-Max (0.7%) | 0.65 | 0.11 | 1.4e-08 | 12 | 1.5 | 3.8e-11 |
| ChromHMM-Promoter_Poised-blood-Avg (0.2%) | -0.12 | 0.12 | 0.32 | 2.8 | 1.4 | 0.0029 |
| ChromHMM-Promoter_Poised-blood-Max (1.6%) | 0.065 | 0.075 | 0.38 | 3.6 | 0.53 | 2.5e-05 |
| ChromHMM-Promoter_Upstream-blood-Avg (0.5%) | -0.15 | 0.12 | 0.22 | 8.7 | 1.2 | 6.9e-08 |
| ChromHMM-Promoter_Upstream-blood-Max (1.3%) | 0.59 | 0.13 | 1e-05 | 12 | 0.86 | 2.1e-05 |
| ChromHMM-Quiescent-blood-Avg (80.0%) | -0.21 | 0.082 | 0.012 | 0.52 | 0.022 | 4.9e-05 |
| ChromHMM-Quiescent-blood-Max (86.0%) | -0.092 | 0.049 | 0.06 | 0.64 | 0.017 | 6e-05 |
| ChromHMM-Repressed_Polycomb-blood-Avg (1.4%) | -0.17 | 0.047 | 0.00037 | 1.1 | 0.36 | 0.089 |
| ChromHMM-Repressed_Polycomb-blood-Max (0.04%) | -0.12 | 0.04 | 0.0038 | 1.3 | 0.22 | 0.0036 |
| ChromHMM-TSS-blood-Avg (0.1%) | -0.35 | 0.15 | 0.022 | 8.9 | 3.9 | 0.0041 |
| ChromHMM-TSS-blood-Max (0.3%) | -0.34 | 0.12 | 0.0051 | 7.5 | 2 | 0.00049 |
| ChromHMM-Transcription_Enhancer_3'-blood-Avg (0.2%) | 0.29 | 0.11 | 0.0067 | 7.7 | 1.4 | 1.4e-06 |
| ChromHMM-Transcription_Enhancer_3'-blood-Max (1.1%) | 0.45 | 0.11 | 2.9e-05 | 7.6 | 0.95 | 4.2e-06 |
| ChromHMM-Transcription_Enhancer_5'-blood-Avg (0.5%) | 0.28 | 0.11 | 0.0085 | 6.2 | 0.96 | 2.6e-10 |
| ChromHMM-Transcription_Enhancer_5'-blood-Max (2.5%) | 0.45 | 0.11 | 5.4e-05 | 5.8 | 0.68 | 1.4e-06 |
| ChromHMM-Transcription.Weak_Enhancer-blood-Avg (0.6%) | -0.035 | 0.1 | 0.72 | 3.1 | 0.87 | 6.9e-07 |
| ChromHMM-Transcription.Weak_Enhancer-blood-Max (2.6%) | 0.19 | 0.09 | 0.035 | 3.9 | 0.64 | 6.8e-10 |
| ChromHMM-Transcriptional_Regulator-blood-Avg (0.4%) | 0.76 | 0.11 | 2.8e-12 | 14 | 1.3 | 1.3e-10 |
| ChromHMM-Transcriptional_Regulator-blood-Max (1.7%) | 0.74 | 0.16 | 2.6e-06 | 9.6 | 1.1 | 2.4e-11 |

**Table S41.** **S-LDSC results for blood-specific variant-level joint model**. Standardized Effect sizes (*τ\**) and Enrichment (E) of the significant blood-specific variant-level DeepSEAV, BasenjiV, Roadmap and ChromHMM annotations from Table S39 and Table S40, conditioned on each other and the non-tissue-specific variant-level joint model (baseline-LD and annotations from Figure S6). Results are meta-analyzed across 11 blood-related traits.

| Annotation | $\tau^*$ | se($\tau^*$) | p($\tau^*$) | $E$ | se ($E$) | p($E$) |
| --- | --- | --- | --- | --- | --- | --- |
| ChromHMM-Active_Enhancer_2-blood-Max (3.5%) | 0.73 | 0.18 | 6.9e-05 | 7.5 | 0.89 | 1.7e-06 |
| ChromHMM-Promoter_Downstream_1-blood-Max (1.1%) | 1 | 0.21 | 1.4e-06 | 15 | 1.5 | 3.5e-05 |
| ChromHMM-Transcriptional_Regulator-blood-Avg (0.4%) | 0.84 | 0.13 | 1.6e-11 | 12 | 1.1 | 7e-09 |
| ChromHMM-Repressed_Polycomb-blood-Avg (1.4%) | -0.11 | 0.032 | 5.8e-04 | 1.5 | 0.35 | 0.0026 |
| Roadmap-H3K4me1-blood-Max (11.2%) | 0.68 | 0.18 | 0.00023 | 5.4 | 0.39 | 6.9e-06 |
| Roadmap-H3K4me3-blood-Avg (1.8%) | -0.77 | 0.19 | 7.4e-05 | 9.4 | 0.71 | 3.8e-05 |

**Table S42.** **The 2 surviving blood-specific Roadmap annotations were sufficient to eliminate 3 of the 4 blood-specific variant-level BasenjiV annotations that were significant in marginal analysis**. Joint S-LDSC model fit of the blood-specific variant-level BasenjiV annotations from Table S39 conditioned on the non-tissue-specific variant-level joint model plus the two Roadmap blood annotations from Table S41. Results are meta-analyzed across 11 blood-related traits.

| Annotation | $\tau^*$ | $\text{se}(\tau^*)$ | $\text{p}(\tau^*)$ | $E$ | $\text{se}(E)$ | $\text{p}(E)$ |
| --- | --- | --- | --- | --- | --- | --- |
| BasenjiV-H3K27ac-blood-Avg (2.5%) | 0.16 | 0.12 | 0.17 | 4.4 | 0.23 | 2e-05 |
| BasenjiV-H3K4me1-blood-Avg (6.2%) | 0.34 | 0.094 | 0.00031 | 3.1 | 0.12 | 2.5e-05 |
| BasenjiV-H3K4me1-blood-Max (13.7%) | 0.18 | 0.11 | 0.096 | 2.5 | 0.086 | 3.9e-05 |
| BasenjiV-H3K4me3-blood-Max (5.5%) | -0.052 | 0.1 | 0.61 | 4.1 | 0.19 | 3.1e-05 |

**Table S43.** **AUROC of various sets of annotations incorporating blood-specific variant-level deep learning annotations in predicting 8,741 fine-mapped autoimmune diseease SNPs**.We report the AUROC for a gradient boosting model trained using each respective set of annotations, either without including baseline-LD model annotations; including baseline-LD model annotations; including annotations from the non-tissue-specific variant-level joint model (baseline-LD model + 7 annotations; baseline-LD+7); or including annotations from the blood-specific joint model (non-tissue-specific joint variant-level joint model + 6 blood annotations; baseline-LD+7+6Blood).

| Feature | finemapped SNPs |
| --- | --- |
| DeepSEAV-blood-Avg/Max | 0.621 |
| BasenjiV-blood-Avg/Max | 0.664 |
| DeepSEAV-blood-All | 0.631 |
| BasenjiV-blood-All | 0.671 |
| baseline-LD | 0.841 |
| baseline-LD + DeepSEAV-blood-Avg/Max | 0.846 |
| baseline-LD + BasenjiV-blood-Avg/Max | 0.852 |
| baseline-LD + DeepSEAV-blood-All | 0.849 |
| baseline-LD + BasenjiV-blood-All | 0.854 |
| baseline-LD+7 | 0.845 |
| baseline-LD+7 + DeepSEAV-blood-Avg/Max | 0.849 |
| baseline-LD+7 + BasenjiV-blood-Avg/Max | 0.853 |
| baseline-LD+7 + DeepSEAV-blood-All | 0.848 |
| baseline-LD+7 + BasenjiV-blood-All | 0.853 |
| baseline-LD+7+6Blood | 0.851 |
| baseline-LD+7+6Blood + DeepSEAV-blood-Avg/Max | 0.853 |
| baseline-LD+7+6Blood + BasenjiV-blood-Avg/Max | 0.854 |
| baseline-LD+7+6Blood + DeepSEAV-blood-All | 0.853 |
| baseline-LD+7+6Blood + BasenjiV-blood-All | 0.853 |

**Table S44.** **Enrichment of fine-mapped autoimmune disease SNPs for blood-specific variant-level annotations**. We report the enrichment in blood-specific variant-level deep learning annotations (along with Jackknife standard error) for 8741 fine-mapped SNPs in immune-related traits^24^.

| Feature | finemapped SNPs |
| --- | --- |
| BasenjiV-DNase-blood-Avg | 2.02 (0.03) |
| BasenjiV-DNase-blood-Max | 1.80 (0.03) |
| BasenjiV-H3K27ac-blood-Avg | 2.14 (0.02) |
| BasenjiV-H3K27ac-blood-Max | 1.85 (0.02) |
| BasenjiV-H3K4me1-blood-Avg | 2.04 (0.03) |
| BasenjiV-H3K4me1-blood-Max | 1.81 (0.02) |
| BasenjiV-H3K4me3-blood-Avg | 2.17 (0.02) |
| BasenjiV-H3K4me3-blood-Max | 1.95 (0.03) |
| DeepSEAV-DNase-blood-Avg | 1.88 (0.03) |
| DeepSEAV-DNase-blood-Max | 1.68 (0.02) |
| DeepSEAV-H3K27ac-blood-Avg | 1.81 (0.02) |
| DeepSEAV-H3K27ac-blood-Max | 1.67 (0.02) |
| DeepSEAV-H3K4me1-blood-Avg | 1.70 (0.01) |
| DeepSEAV-H3K4me1-blood-Max | 1.55 (0.02) |
| DeepSEAV-H3K4me3-blood-Avg | 1.95 (0.02) |
| DeepSEAV-H3K4me3-blood-Max | 1.76 (0.02) |

**Table S45.** **S-LDSC results for marginal analysis of brain-specific variant-level DeepSEAV and BasenjiV annotations**. Standardized Effect sizes (*τ\**) and Enrichment (E) of 8 brain-specific variant-level DeepSEAV and 8 brain-specific variant-level BasenjiV annotations corresponding to 4 chromatin marks, DNase-1 Hypersensitivity Sites (DHS), H3K27ac, H3K4me1 and H3K4me3, when conditioned on the non-tissue-specific variant-level joint model (baseline-LD and annotations from Figure S6). Results are meta-analyzed across 8 brain-related traits..

| Annotation | $\tau^*$ | se($\tau^*$) | p($\tau^*$) | $E$ | se ($E$) | p($E$) |
| --- | --- | --- | --- | --- | --- | --- |
| DeepSEAV-DNase-brain-Avg (2.9%) | 0.5 | 0.081 | 8.6e-10 | 2.9 | 0.23 | 5e-05 |
| DeepSEAV-DNase-brain-Max (4.5%) | 0.41 | 0.067 | 1.7e-09 | 2.5 | 0.19 | 3.4e-05 |
| DeepSEAV-H3K27ac-brain-Avg (3.2%) | 0.6 | 0.066 | 5.4e-20 | 2.3 | 0.072 | 1.1e-08 |
| DeepSEAV-H3K27ac-brain-Max (7.6%) | 0.55 | 0.062 | 1.5e-18 | 2.1 | 0.076 | 8.8e-09 |
| DeepSEAV-H3K4me1-brain-Avg (4.7%) | 0.37 | 0.06 | 5.8e-10 | 2.1 | 0.06 | 3.7e-11 |
| DeepSEAV-H3K4me1-brain-Max (10.5%) | 0.36 | 0.053 | 5.3e-12 | 1.9 | 0.07 | 2.6e-11 |
| DeepSEAV-H3K4me3-brain-Avg (1.9%) | 0.77 | 0.095 | 4.6e-16 | 4.4 | 0.26 | 1.7e-08 |
| DeepSEAV-H3K4me3-brain-Max (3.4%) | 0.77 | 0.085 | 1e-19 | 3.8 | 0.19 | 2.2e-08 |
| BasenjiV-DNase-brain-Avg (2.9%) | 0.47 | 0.078 | 1.3e-09 | 2.7 | 0.16 | 8.8e-08 |
| BasenjiV-DNase-brain-Max (4.5%) | 0.49 | 0.076 | 9.3e-11 | 2.8 | 0.18 | 1.2e-07 |
| BasenjiV-H3K27ac-brain-Avg (3.2%) | 0.34 | 0.049 | 3.1e-12 | 2.3 | 0.079 | 1.6e-08 |
| BasenjiV-H3K27ac-brain-Max (7.6%) | 0.4 | 0.058 | 3.6e-12 | 2.3 | 0.12 | 1.7e-06 |
| BasenjiV-H3K4me1-brain-Avg (4.7%) | 0.37 | 0.059 | 4.1e-10 | 2.1 | 0.059 | 7.6e-11 |
| BasenjiV-H3K4me1-brain-Max (10.5%) | 0.33 | 0.09 | 0.00022 | 2 | 0.062 | 2.6e-10 |
| BasenjiV-H3K4me3-brain-Avg (1.9%) | 0.55 | 0.074 | 1.1e-13 | 3.6 | 0.18 | 5.2e-07 |
| BasenjiV-H3K4me3-brain-Max (3.4%) | 0.77 | 0.089 | 8.2e-18 | 4 | 0.3 | 8.8e-06 |

**Table S46.** **S-LDSC for marginal analysis of brain-specific Roadmap and ChromHMM annotations**.Standardized Effect sizes (*τ\**) and Enrichment (E) of 8 brain-specific Roadmap and 40 brain-specific ChromHMM annotations, when conditioned on the non-tissue-specific variant-level joint model annotations (baseline-LD and annotations from Figure S6). Results are meta-analyzed across 8 brain-related traits.

| Annotation | $\tau^*$ | se($\tau^*$) | p($\tau^*$) | $E$ | se ($E$) | p($E$) |
| --- | --- | --- | --- | --- | --- | --- |
| Roadmap-DNase-brain-Avg (2.4%) | 0.55 | 0.084 | 5.5e-11 | 4.9 | 0.41 | 5.4e-06 |
| Roadmap-DNase-brain-Max (5.1%) | 0.39 | 0.079 | 7.1e-07 | 3.5 | 0.21 | 2.4e-06 |
| Roadmap-H3K27ac-brain-Avg (3.2%) | 0.34 | 0.09 | 0.00019 | 3.5 | 0.15 | 1.1e-10 |
| Roadmap-H3K27ac-brain-Max (8.2%) | 0.37 | 0.095 | 0.00012 | 2.9 | 0.11 | 7.6e-11 |
| Roadmap-H3K4me1-brain-Avg (4.4%) | 0.48 | 0.12 | 5e-05 | 3.2 | 0.12 | 4.4e-11 |
| Roadmap-H3K4me1-brain-Max (11.2%) | 0.46 | 0.12 | 0.00016 | 2.7 | 0.09 | 3.6e-11 |
| Roadmap-H3K4me3-brain-Avg (1.5%) | 0.86 | 0.15 | 7.7e-09 | 6.8 | 0.37 | 1.1e-12 |
| Roadmap-H3K4me3-brain-Max (2.9%) | 0.72 | 0.12 | 1.8e-09 | 5.7 | 0.28 | 4e-12 |
| ChromHMM-DNase-brain-Avg (0.8%) | -0.12 | 0.096 | 0.21 | 0.86 | 0.8 | 0.45 |
| ChromHMM-DNase-brain-Max (3.0%) | 0.062 | 0.072 | 0.39 | 1.8 | 0.5 | 0.0031 |
| ChromHMM-Active-Enhancer-1-brain-Avg (0.4%) | 0.24 | 0.086 | 0.0061 | 3.6 | 0.5 | 0.06 |
| ChromHMM-Active-Enhancer-1-brain-Max (1.5%) | 0.32 | 0.1 | 0.002 | 3.5 | 0.45 | 0.056 |
| ChromHMM-Active_Enhancer_2-brain-Avg (0.4%) | 0.39 | 0.083 | 3.4e-06 | 4.8 | 0.48 | 0.00043 |
| ChromHMM-Active_Enhancer_2-brain-Max (2.2%) | 0.36 | 0.082 | 8e-06 | 4 | 0.3 | 1e-05 |
| ChromHMM-Enhancer_acetylation-brain-Avg (0.3%) | 0.13 | 0.054 | 0.02 | 2.3 | 0.29 | 0.077 |
| ChromHMM-Enhancer_acetylation-brain-Max (2.2%) | 0.19 | 0.068 | 0.0047 | 2.5 | 0.29 | 0.016 |
| ChromHMM-Active_Enhancer_Flanking-brain-Avg (0.8%) | 0.13 | 0.057 | 0.024 | 2.5 | 0.21 | 0.0031 |
| ChromHMM-Active_Enhancer_Flanking-brain-Max (3.1%) | 0.16 | 0.062 | 0.011 | 2.3 | 0.17 | 0.002 |

**Table S46.** *(continued)*
| Annotation | $\tau^*$ | $\text{se}(\tau^*)$ | $\text{p}(\tau^*)$ | $E$ | $\text{se}(E)$ | $\text{p}(E)$ |
| --- | --- | --- | --- | --- | --- | --- |
| ChromHMM-Weak_Enhancer_1-brain-Avg (0.4%) | 0.19 | 0.073 | 0.0087 | 3.9 | 0.56 | 0.03 |
| ChromHMM-Weak_Enhancer_1-brain-Max (1.7%) | 0.29 | 0.091 | 0.0012 | 4.1 | 0.51 | 0.014 |
| ChromHMM-Weak_Enhancer_2-brain-Avg (1.3%) | 0.25 | 0.074 | 0.00065 | 2.8 | 0.27 | 3.5e-10 |
| ChromHMM-Weak_Enhancer_2-brain-Max (5.6%) | 0.27 | 0.085 | 0.0015 | 2.5 | 0.23 | 3.6e-10 |
| ChromHMM-Heterochromatin-brain-Avg (1.2%) | -<br>0.00042 | 0.021 | 0.98 | 0.37 | 0.12 | 0.016 |
| ChromHMM-Heterochromatin-brain-Max (2.3%) | 0.0051 | 0.021 | 0.81 | 0.42 | 0.1 | 0.0058 |
| ChromHMM-Promoter_Bivalent-brain-Avg (0.2%) | 0.15 | 0.06 | 0.01 | 7.4 | 0.97 | 0.00052 |
| ChromHMM-Promoter_Bivalent-brain-Max (0.6%) | 0.29 | 0.058 | 3e-07 | 7.2 | 0.57 | 2.7e-05 |
| ChromHMM-Promoter_Downstream_1-brain-Avg (0.4%) | 0.29 | 0.072 | 7e-05 | 7.7 | 0.9 | 1.5e-05 |
| ChromHMM-Promoter_Downstream_1-brain-Max (0.6%) | 0.47 | 0.077 | 1.2e-09 | 9.1 | 0.83 | 1.9e-08 |
| ChromHMM-Promoter_Downstream_2-brain-Avg (0.2%) | 0.12 | 0.069 | 0.075 | 4.4 | 1.2 | 0.095 |
| ChromHMM-Promoter_Downstream_2-brain-Max (0.4%) | 0.22 | 0.068 | 0.0011 | 5.5 | 0.98 | 0.00026 |
| ChromHMM-Promoter_Poised-brain-Avg (0.2%) | 0.23 | 0.083 | 0.0064 | 4.3 | 0.81 | 0.059 |
| ChromHMM-Promoter_Poised-brain-Max (1.1%) | 0.24 | 0.1 | 0.018 | 3.7 | 0.75 | 0.11 |
| ChromHMM-Promoter_Upstream-brain-Avg (0.5%) | 0.46 | 0.11 | 2.7e-05 | 9.6 | 0.82 | 1e-10 |
| ChromHMM-Promoter_Upstream-brain-Max (1.0%) | 0.5 | 0.11 | 5.9e-06 | 8.1 | 0.6 | 1.3e-11 |

**Table S46.** *(continued)*
| Annotation | $\tau^*$ | $se(\tau^*)$ | $p(\tau^*)$ | $E$ | $se(E)$ | $p(E)$ |
| --- | --- | --- | --- | --- | --- | --- |
| ChromHMM-Quiescent-brain-Avg (77.6%) | -0.22 | 0.054 | 5.1e-05 | 0.81 | 0.013 | 8.1e-07 |
| ChromHMM-Quiescent-brain-Max (86.0%) | -0.088 | 0.029 | 0.0025 | 0.87 | 0.0098 | 3.4e-06 |
| ChromHMM-Repressed_Polycomb-brain-Avg (1.4%) | -0.069 | 0.025 | 0.005 | 1 | 0.16 | 0.31 |
| ChromHMM-Repressed_Polycomb-brain-Max (4.3%) | -0.046 | 0.024 | 0.054 | 1.2 | 0.094 | 0.012 |
| ChromHMM-TSS-brain-Avg (0.2%) | 0.32 | 0.092 | 0.00048 | 7.9 | 1.6 | 0.00061 |
| ChromHMM-TSS-brain-Max (0.4%) | 0.44 | 0.097 | 5e-06 | 9 | 1.3 | 0.0017 |
| ChromHMM-Transcription_Enhancer_3'-brain-Avg (0.3%) | 0.12 | 0.038 | 0.0027 | 3.6 | 0.39 | 0.039 |
| ChromHMM-Transcription_Enhancer_3'-brain-Max (1.0%) | 0.092 | 0.039 | 0.017 | 2.9 | 0.31 | 0.0022 |
| ChromHMM-Transcription_Enhancer_5'-brain-Avg (0.5%) | 0.027 | 0.032 | 0.4 | 2.8 | 0.24 | 1.7e-08 |
| ChromHMM-Transcription_Enhancer_5'-brain-Max (1.6%) | 0.083 | 0.034 | 0.016 | 2.8 | 0.2 | 5.4e-10 |
| ChromHMM-Transcription_Weak_Enhancer-brain-Avg (0.6%) | 0.21 | 0.049 | 2.8e-05 | 3.3 | 0.33 | 0.00026 |
| ChromHMM-Transcription_Weak_Enhancer-brain-Max (1.8%) | 0.22 | 0.05 | 1.7e-05 | 3 | 0.25 | 2.2e-11 |
| ChromHMM-Transcriptional_Regulator-brain-Avg (0.3%) | 0.11 | 0.045 | 0.019 | 5 | 0.52 | 1.8e-06 |
| ChromHMM-Transcriptional_Regulator-brain-Max (0.9%) | 0.056 | 0.047 | 0.24 | 3.7 | 0.35 | 3.3e-06 |

**Table S47.** **S-LDSC results for brain-specific variant-level joint model**. Standardized Effect sizes (*τ\**) and Enrichment (E) of the significant brain-specific variant-level DeepSEAV, BasenjiV, Roadmap and ChromHMM annotations from Table S45 and Table S46, conditioned on each other and the non-tissue-specific variant-level joint model annotations (baseline-LD and annotations from Figure S6). Results are meta-analyzed across 8 brain-related traits.

| Annotation | $\tau^*$ | se($\tau^*$) | p($\tau^*$) | $E$ | se ($E$) | p($E$) |
| --- | --- | --- | --- | --- | --- | --- |
| BasenjiV-H3K27ac-brain-Max (7.7%) | 0.27 | 0.059 | 5.4e-06 | 2.3 | 0.077 | 1.3e-08 |
| ChromHMM-Promoter_Bivalent-brain-Max (0.6%) | 0.21 | 0.056 | 0.00014 | 6.9 | 0.56 | 3.3e-06 |
| ChromHMM-Transcriptional-Weak-Enhancer-brain-Max (1.8%) | 0.26 | 0.06 | 1.2e-05 | 3 | 0.25 | 4.8e-16 |
| ChromHMM-Weak_Enhancer_2-brain-Avg (1.3%) | 0.27 | 0.078 | 0.00067 | 2.7 | 0.25 | 3.4e-12 |
| DeepSEAV-H3K4me3-brain-Max (3.4%) | 0.35 | 0.09 | 0.0001 | 3.6 | 0.2 | 5.8e-08 |
| Roadmap-H3K4me3-brain-Max (2.9%) | 0.4 | 0.091 | 1.3e-05 | 5.2 | 0.28 | 1.5e-10 |

**Table S48.** **The surviving brain-specific Roadmap annotations do not eliminate any brain specific variant-level DeepSEAV and brain-specific variant-level BasenjiV annotations that were significant in marginal analysis**. Standardized Effect sizes (*τ\**) and Enrichment (E) of brain-specific DeepSEAV and brain-specific BasenjiV annotations, conditioned on the non-tissue-specific variant-level joint model annotations and 1 significant Roadmap brain annotation (Roadmap-H3K4me3-brain-Max) that is significant in the joint model in Table S47. Results are meta-analyzed across 8 brain-related traits.

| Annotation | $\tau^*$ | $se(\tau^*)$ | $p(\tau^*)$ | $E$ | $se(E)$ | $p(E)$ |
| --- | --- | --- | --- | --- | --- | --- |
| DeepSEAV-DNase-brain-Avg (2.9%) | 0.31 | 0.081 | 0.00011 | 2.8 | 0.24 | 3.8e-08 |
| DeepSEAV-DNase-brain-Max (4.5%) | 0.25 | 0.069 | 0.00022 | 2.5 | 0.2 | 2.3e-08 |
| DeepSEAV-H3K27ac-brain-Avg (3.2%) | 0.47 | 0.068 | 2.4e-12 | 2.4 | 0.072 | 9e-12 |
| DeepSEAV-H3K27ac-brain-Max (7.7%) | 0.43 | 0.064 | 1.7e-11 | 2.2 | 0.072 | 5.6e-12 |
| DeepSEAV-H3K4me1-brain-Avg (4.7%) | 0.3 | 0.054 | 1.6e-08 | 2.1 | 0.057 | 4.1e-13 |
| DeepSEAV-H3K4me1-brain-Max (10.5%) | 0.29 | 0.053 | 4.9e-08 | 1.9 | 0.068 | 5e-13 |
| DeepSEAV-H3K4me3-brain-Avg (1.9%) | 0.5 | 0.092 | 5.7e-08 | 4.4 | 0.26 | 1.3e-11 |
| DeepSEAV-H3K4me3-brain-Max (3.4%) | 0.54 | 0.087 | 6.4e-10 | 3.8 | 0.19 | 6.7e-12 |
| BasenjiV-DNase-brain-Avg (2.9%) | 0.34 | 0.064 | 8.7e-08 | 2.8 | 0.17 | 1.4e-10 |
| BasenjiV-DNase-brain-Max (4.5%) | 0.36 | 0.068 | 1.4e-07 | 2.7 | 0.15 | 1.5e-10 |
| BasenjiV-H3K27ac-brain-Avg (3.2%) | 0.25 | 0.055 | 4e-06 | 2.4 | 0.11 | 1.3e-10 |
| BasenjiV-H3K27ac-brain-Max (7.7%) | 0.34 | 0.056 | 1.1e-09 | 2.3 | 0.073 | 1.3e-10 |
| BasenjiV-H3K4me1-brain-Avg (4.7%) | 0.3 | 0.053 | 2.2e-08 | 2.1 | 0.059 | 7e-12 |
| BasenjiV-H3K4me1-brain-Max (10.5%) | 0.29 | 0.087 | 0.00077 | 2.1 | 0.061 | 1.6e-12 |
| BasenjiV-H3K4me3-brain-Avg (1.9%) | 0.31 | 0.081 | 0.00014 | 4.1 | 0.28 | 4.8e-11 |
| BasenjiV-H3K4me3-brain-Max (3.4%) | 0.61 | 0.08 | 2.3e-14 | 3.8 | 0.17 | 3.5e-11 |

**Table S49.** **The surviving brain-specific ChromHMM annotations do not eliminate any brain specific variant-level DeepSEAV and brain-specific variant-level BasenjiV annotations that were significant in marginal analysis**. Standardized Effect sizes (*τ\**) and Enrichment (E) of brain-specific variant-level DeepSEAV and brain-specific variant-level BasenjiV annotations, conditioned on the non-tissue-specific variant-level joint model annotations and 3 significant ChromHMM brain annotations that are significant in the joint model in Table S47. Results are meta-analyzed across 8 brain-related traits.

| Annotation | $\tau^*$ | $se(\tau^*)$ | $p(\tau^*)$ | $E$ | $se(E)$ | $p(E)$ |
| --- | --- | --- | --- | --- | --- | --- |
| DeepSEAV-DNase-brain-Avg (2.9%) | 0.44 | 0.079 | 2e-08 | 2.8 | 0.23 | 3.5e-06 |
| DeepSEAV-DNase-brain-Max (4.5%) | 0.36 | 0.067 | 5.9e-08 | 2.5 | 0.2 | 1.3e-06 |
| DeepSEAV-H3K27ac-brain-Avg (3.2%) | 0.55 | 0.065 | 2.6e-17 | 2.2 | 0.085 | 7.1e-08 |
| DeepSEAV-H3K27ac-brain-Max (7.6%) | 0.5 | 0.062 | 5.3e-16 | 2.1 | 0.086 | 5.4e-08 |
| DeepSEAV-H3K4me1-brain-Avg (4.7%) | 0.28 | 0.053 | 9.9e-08 | 1.9 | 0.07 | 4.4e-08 |
| DeepSEAV-H3K4me1-brain-Max (10.5%) | 0.28 | 0.052 | 5.5e-08 | 1.8 | 0.08 | 2.3e-08 |
| DeepSEAV-H3K4me3-brain-Avg (1.9%) | 0.73 | 0.094 | 8.4e-15 | 4.2 | 0.26 | 5.4e-08 |
| DeepSEAV-H3K4me3-brain-Max (3.4%) | 0.72 | 0.086 | 3.9e-17 | 3.6 | 0.2 | 6.3e-08 |
| BasenjiV-DNase-brain-Avg (2.9%) | 0.4 | 0.071 | 1.7e-08 | 2.7 | 0.18 | 7.2e-08 |
| BasenjiV-DNase-brain-Max (4.5%) | 0.42 | 0.073 | 5.4e-09 | 2.7 | 0.16 | 5.1e-08 |
| BasenjiV-H3K27ac-brain-Avg (3.2%) | 0.30 | 0.05 | 1.6e-09 | 2.2 | 0.13 | 1.1e-07 |
| BasenjiV-H3K27ac-brain-Max (7.6%) | 0.36 | 0.057 | 2.3e-10 | 2.2 | 0.084 | 5.7e-08 |
| BasenjiV-H3K4me1-brain-Avg (4.7%) | 0.31 | 0.057 | 6.1e-08 | 2 | 0.072 | 6.9e-08 |
| BasenjiV-H3K4me1-brain-Max (10.5%) | 0.30 | 0.089 | 0.00061 | 2 | 0.054 | 4e-09 |
| BasenjiV-H3K4me3-brain-Avg (1.9%) | 0.52 | 0.073 | 1e-12 | 3.8 | 0.31 | 2.4e-07 |
| BasenjiV-H3K4me3-brain-Max (3.4%) | 0.68 | 0.096 | 1.5e-12 | 3.5 | 0.2 | 1.4e-07 |

**Table S50.**
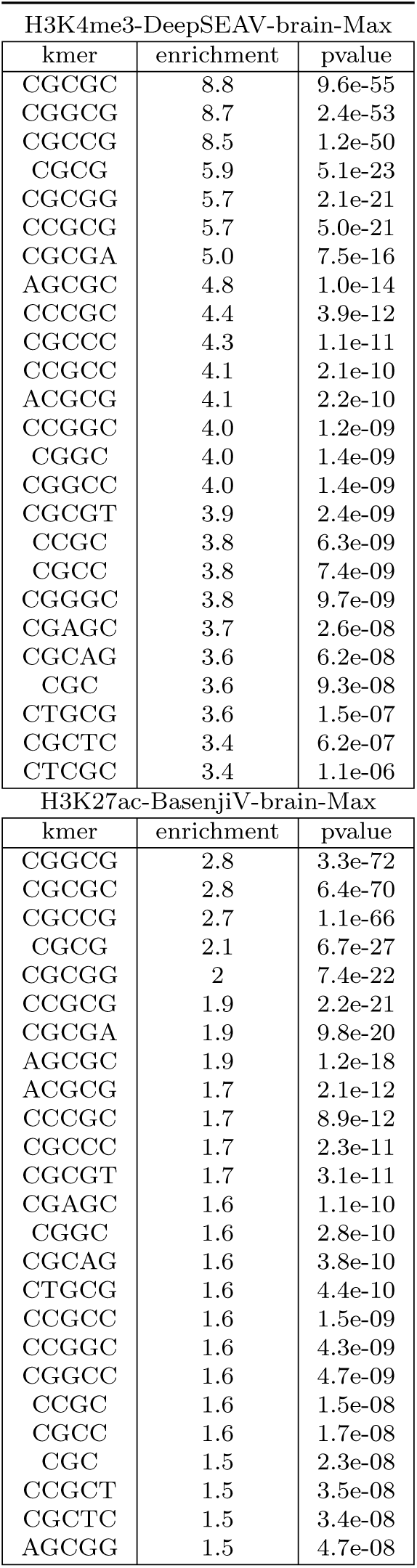
**Weighted** *k***-mer enrichments for significant variant level brain-specific variant-level DeepSEAV and BasenjiV annotations**. We report the weighted *k*-mer enrichment and enrichment p-value for the top significant enriched *k*-mers (1 *k* 5) for H3K4me3-DeepSEAV-brain-Max and H3K27ac-BasenjiV-brain-Max which were jointly significant in the brain-specific variant-level joint model. The Bonferonni corrcection threshold is 0.05*/*(4 682) where 682 is the number of *k*-mers analyzed for the 2 significant variant-level annotations analyzed here and the 2 significant allelic-effect annotations analyzed in Table S11.

## Figures

**Figure S1.**
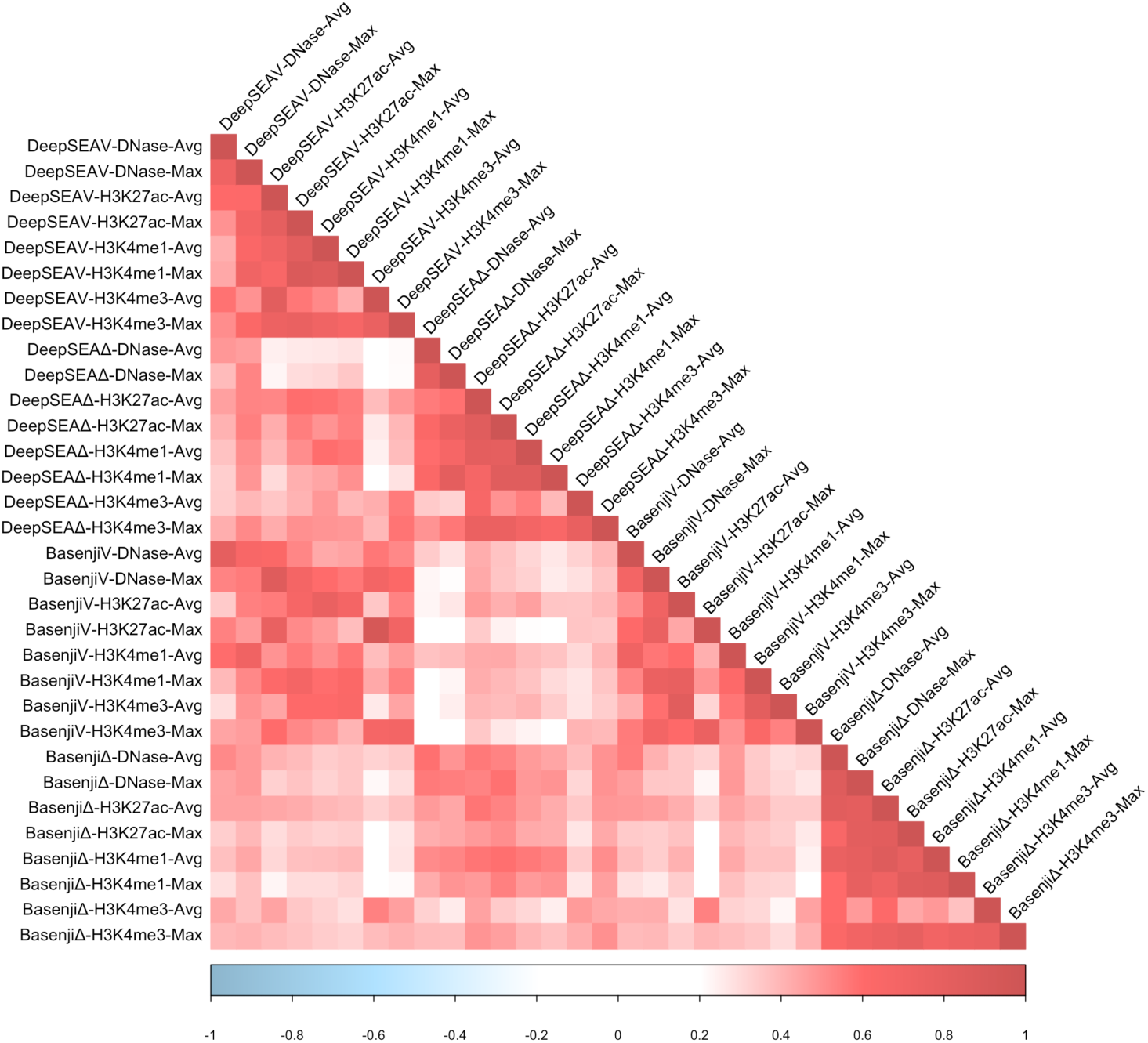
**Correlations between deep learning annotations**. We report correlations between each pair of variant-level (DeepSEA and Basenji) and/or allelic-effect (DeepSEAΔ and BasenjiΔ) annotations. When comparing pairs of annotations that differed only in their aggregation strategy (Avg/Max), chromatin mark (DNase/H3K27ac/H3K4me1/H3K4me3), model (DeepSEA/Basenji) or type (variant-level/allelic-effect), respectively, we observed large correlations across aggregation strategies (average *r* = 0.71), chromatin marks (average *r* = 0.58), models (average *r* = 0.54) and types (average *r* = 0.48).

**Figure S2.**
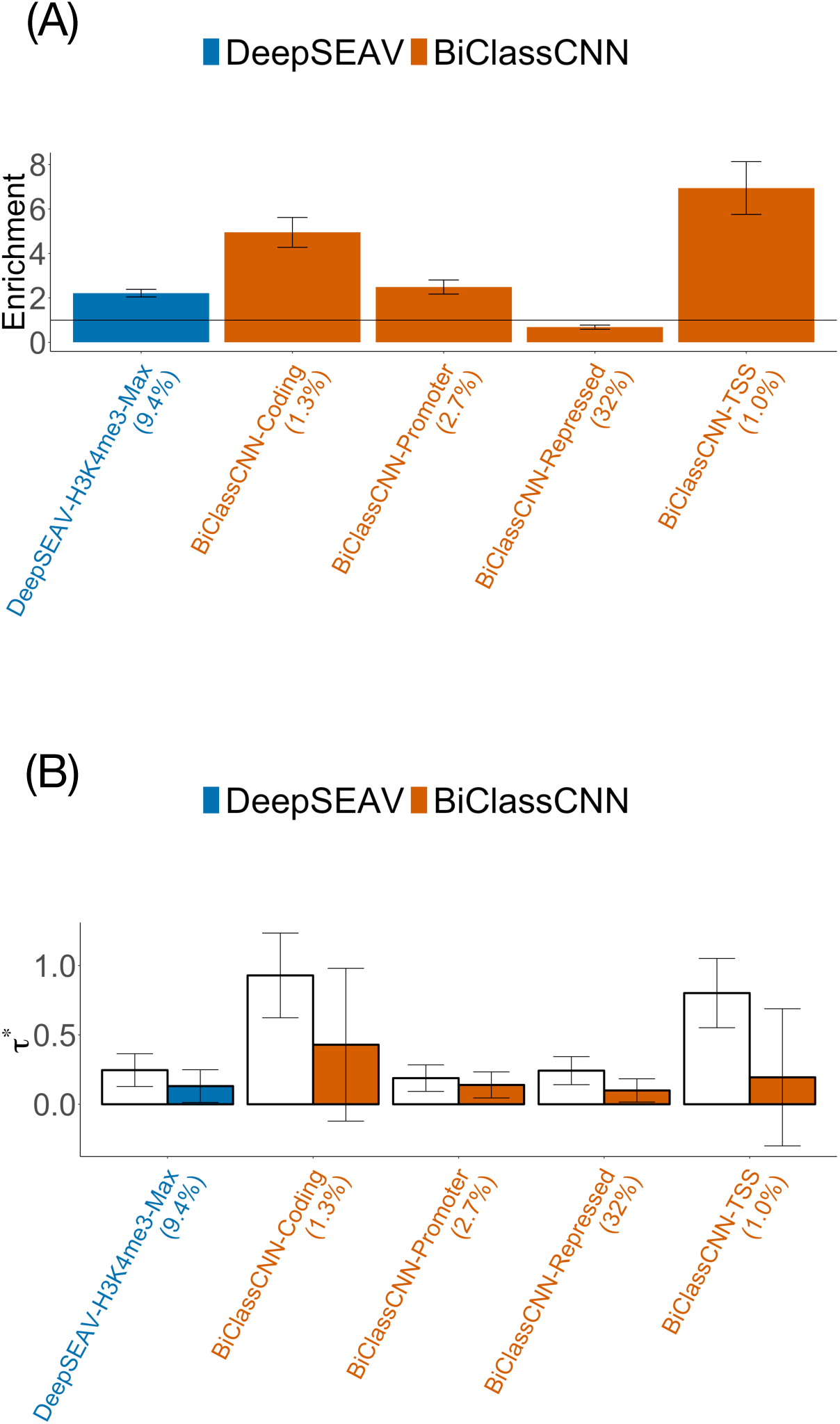
**Disease informativeness of non-tissue-specific variant-level deep learning annotations**: (A) Heritability enrichment, conditional on the baseline-LD model. Horizontal line denotes no enrichment. (B) Standardized effect size (*τ\**) conditional on either the baseline-LD model (marginal analyses: left column, white) or the baseline-LD model plus 1 Roadmap, 1 ChromHMM, 3 LocalGCcontent and 2 pLI annotations (non-tissue-specific variant-level joint model: right column, shaded). Results are meta-analyzed across 41 traits. Results are displayed only for annotations with significant *τ\** in marginal analyses after correcting for 106 (variant-level + allelic-effect) non-tissue-specific annotations tested (*P <* 0.05*/*106). None of these annotations were significant after conditioning on the non-tissue-specific variant-level joint model (right column). Error bars denote 95% confidence intervals. Numerical results are reported in Table S29, Table S30 and Table S34.

**Figure S3.**
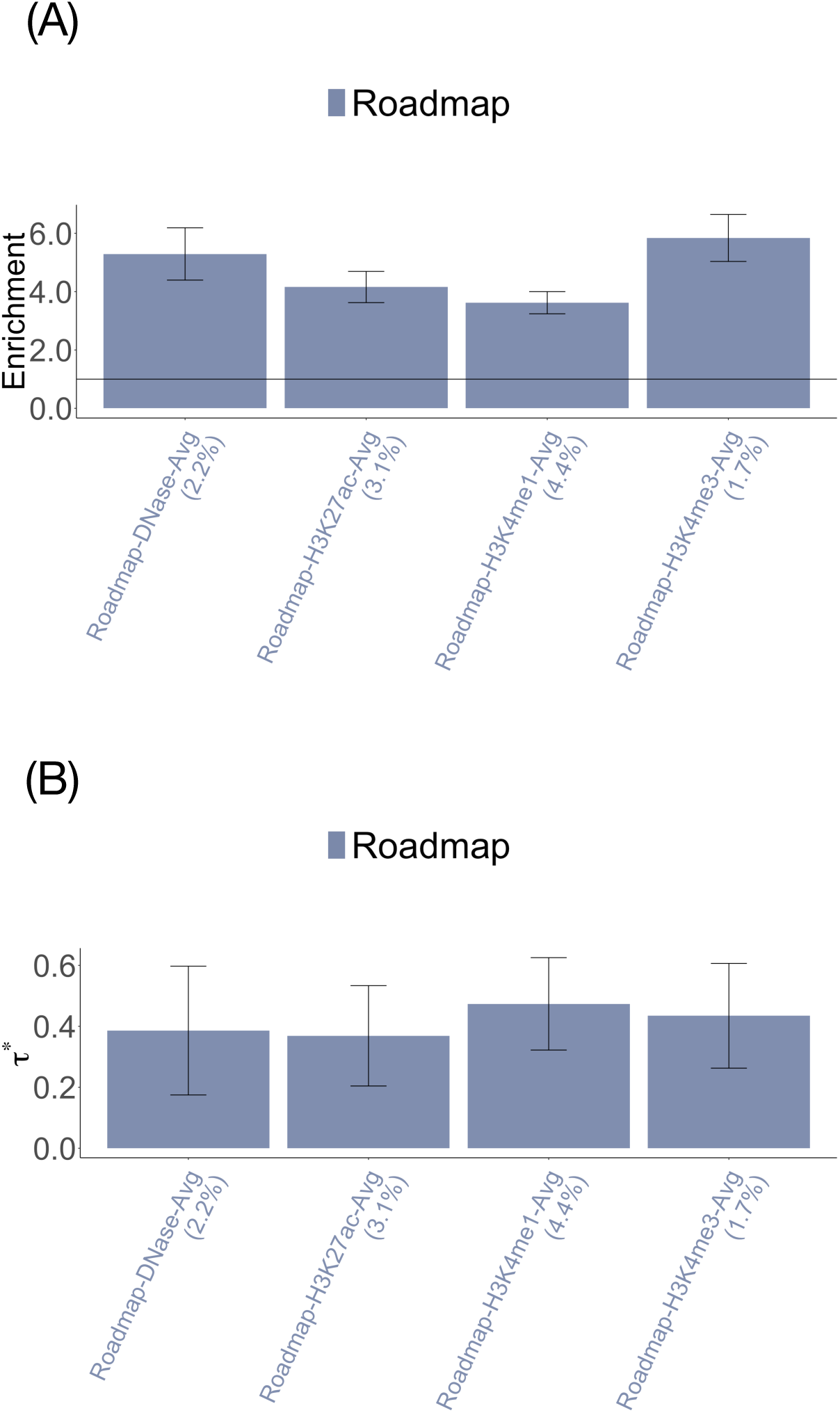
**Disease informativeness of non-tissue-specific Roadmap annotations in marginal analysis**. (A) Heritability enrichment, conditioned on the baseline-LD model. Horizontal line denotes no enrichment. (B) Standardized effect size (*τ\**) conditioned on the baseline-LD model. Results are meta-analyzed across 41 traits. Results are displayed only for annotations with significant *τ\** in marginal analyses after correcting for 106 (variant-level + allelic-effect) non-tissue-specific annotations tested (*P <* 0.05*/*106). Error bars denote 95% confidence intervals. Numerical results are reported in Table S32.

**Figure S4.**
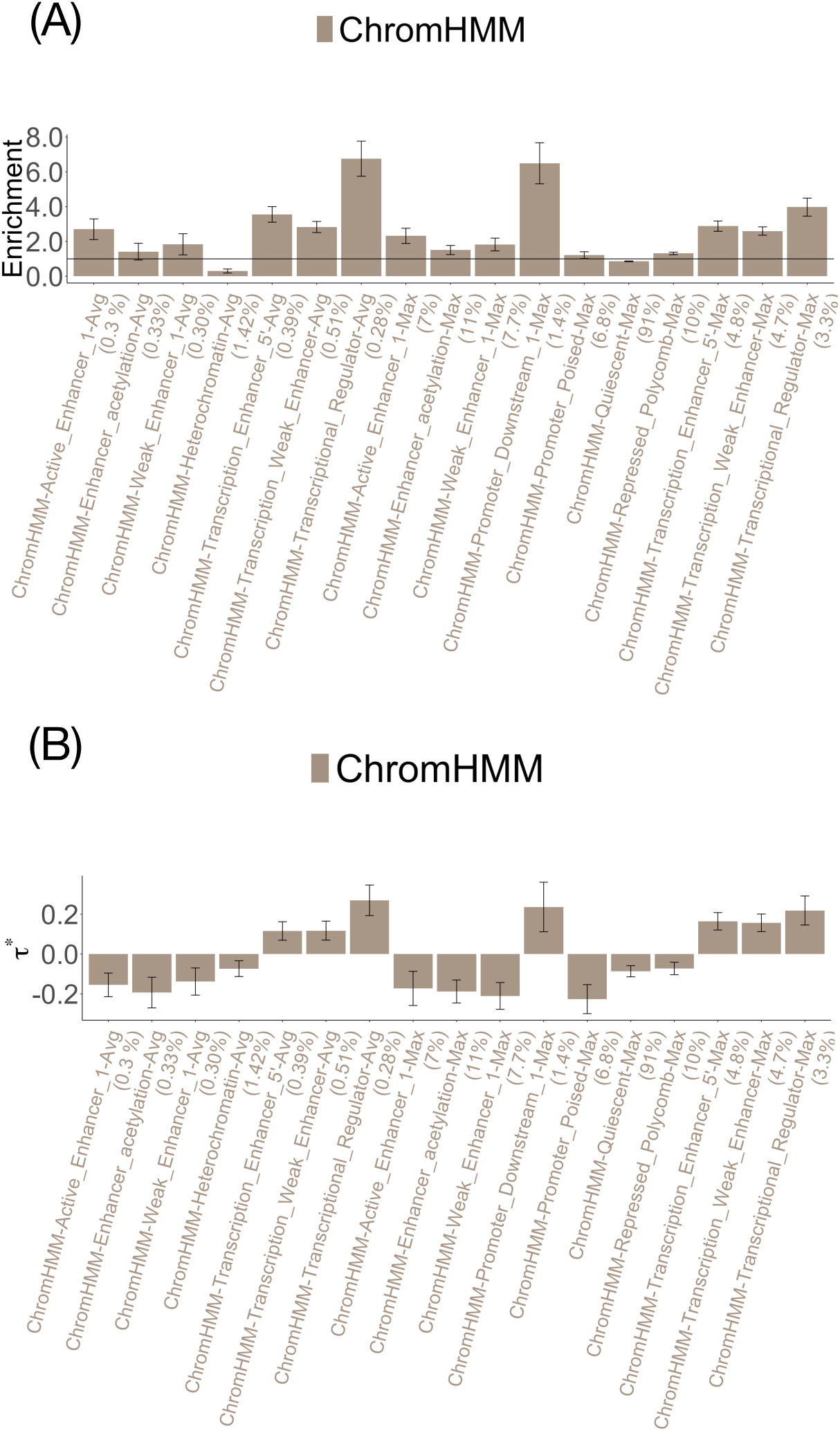
**Disease informativeness of non-tissue-specific ChromHMM annotations in marginal analysis**. (A) Heritability enrichment, conditioned on the baseline-LD model. Horizontal line denotes no enrichment. (B) Standardized effect size (*τ\**) conditioned on the baseline-LD model. Results are meta-analyzed across 41 traits. Results are displayed only for annotations with significant *τ\** in marginal analyses after correcting for 106 (variant-level + allelic-effect) non-tissue-specific annotations tested (*P <* 0.05*/*106). Error bars denote 95% confidence intervals. Numerical results are reported in Table S32.

**Figure S5.**
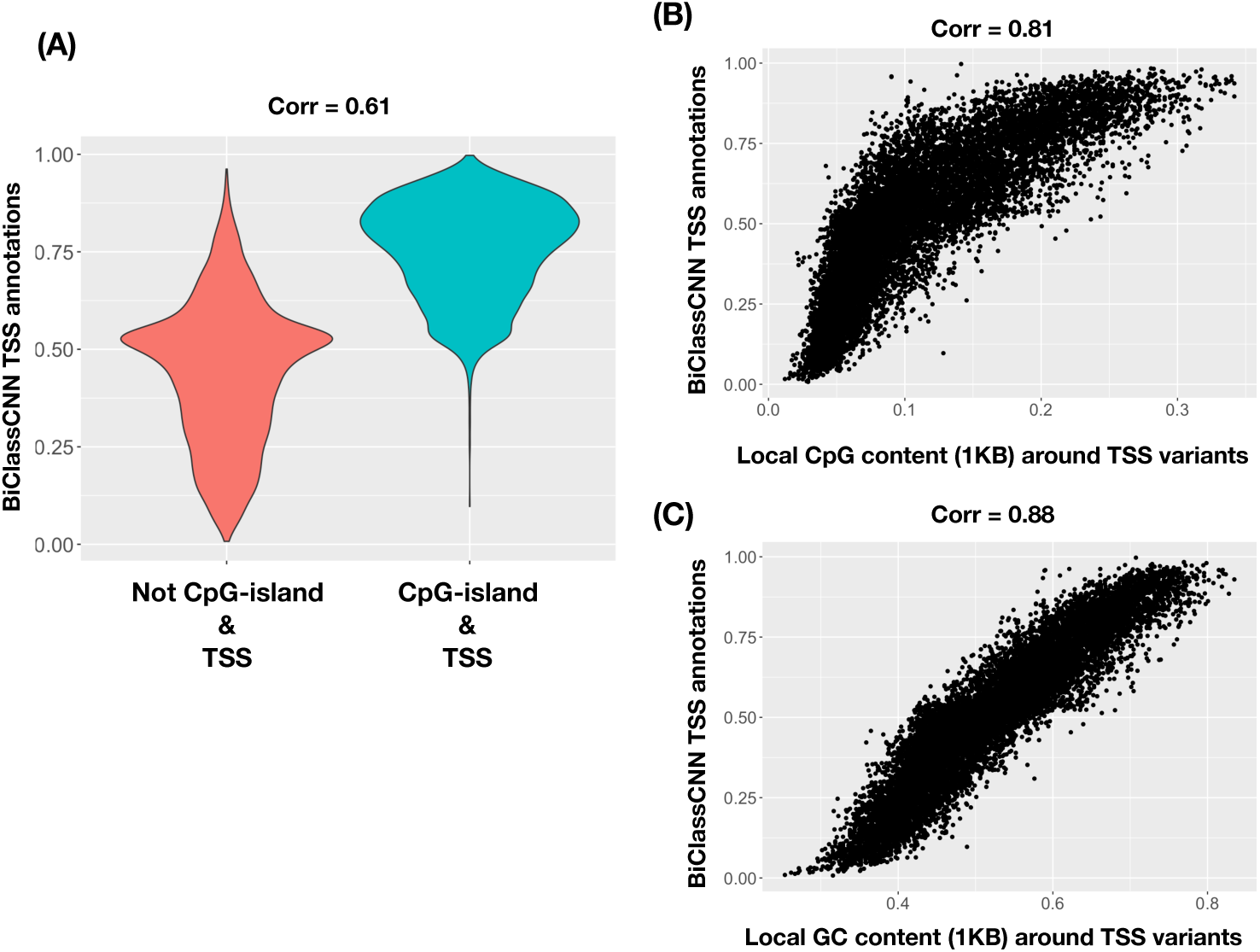
**BiClassCNN-TSS is highly correlated with CpG-island, local CpG-content(***±*1**kb) and local GC-content(***±*1**kb) annotations**. (A) Violin plot of the Bi-ClassCNN TSS annotation for variants that are TSS and fall in a CpG island and variants that are TSS but do not fall in a CpG island. (B) Scatter plot of the BiClassCNN TSS annotation with the local GC content (1KB) around the variant for all TSS variants. (C) Scatter plot of the BiClassCNN TSS annotation with the local CpG content (1KB) around the variant for all TSS variants. Here, by TSS variants, we imply variants that are annotated to fall in a TSS region as predicted by Segway^77, 78^. The BiClassCNN TSS annotations showed strong correlation with all three features - CpG-island, Local GC and Local CpG content at the TSS variants with the correlation with the Local GC content at TSS variants being the strongest.

**Figure S6.**
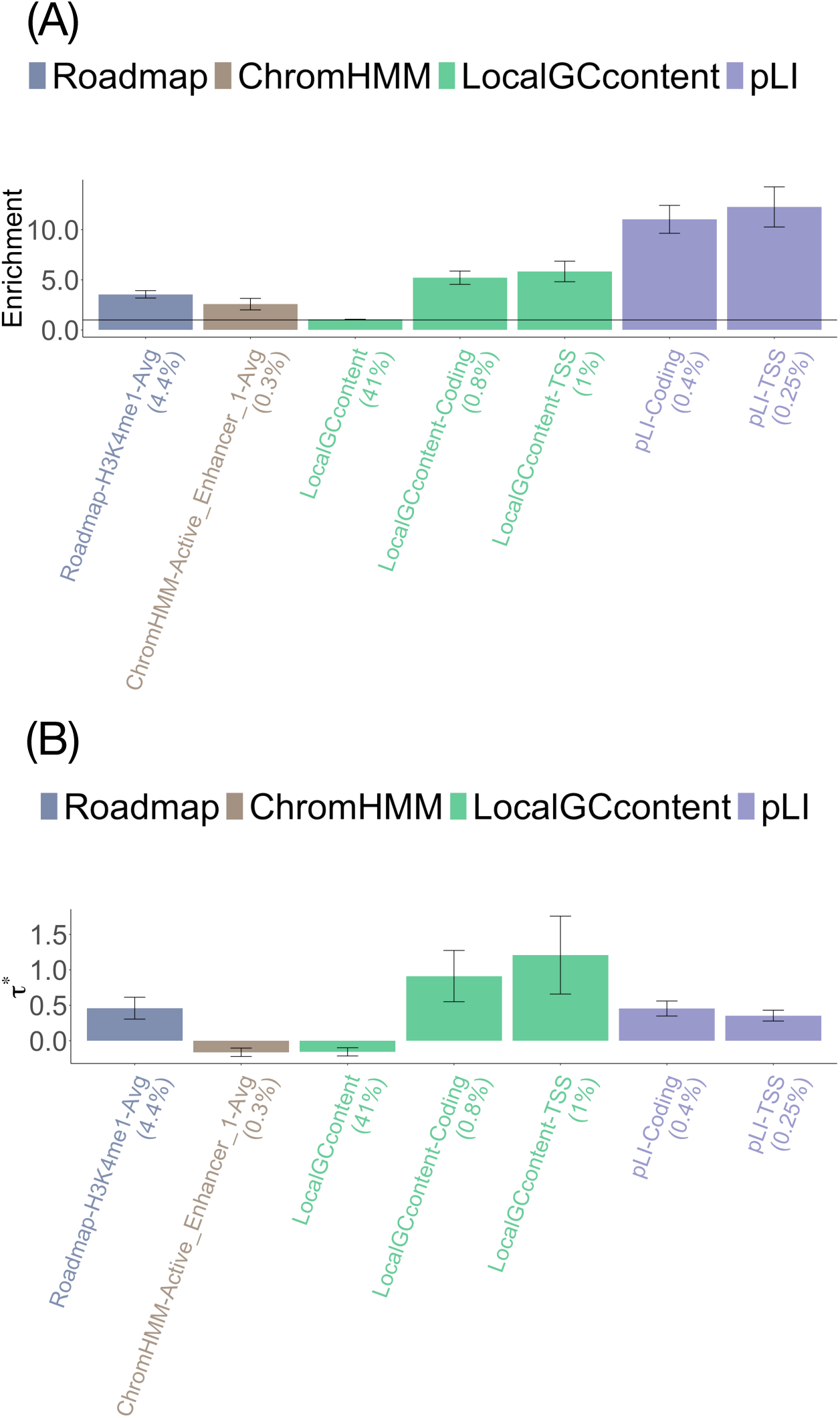
**Disease informativeness of non-tissue-specific Roadmap, ChromHMM, LocalGCcontent and pLI annotations in non-tissue-specific variant-level joint model**. (A) Heritability enrichment, conditioned on the baseline-LD model. Horizontal line denotes no enrichment. (B) Standardized effect size (*τ\**) conditioned on each other as well as the baseline-LD model.Results are meta-analyzed across 41 traits. Results are displayed only for annotations with significant *τ^*^*in marginal analyses after correcting for 106 (variant-level + allelic-effect) non-tissue-specific annotations tested (*P <* 0.05*/*106). Error bars denote 95% confidence intervals. Numerical results are reported in Table S34.

**Figure S7.**
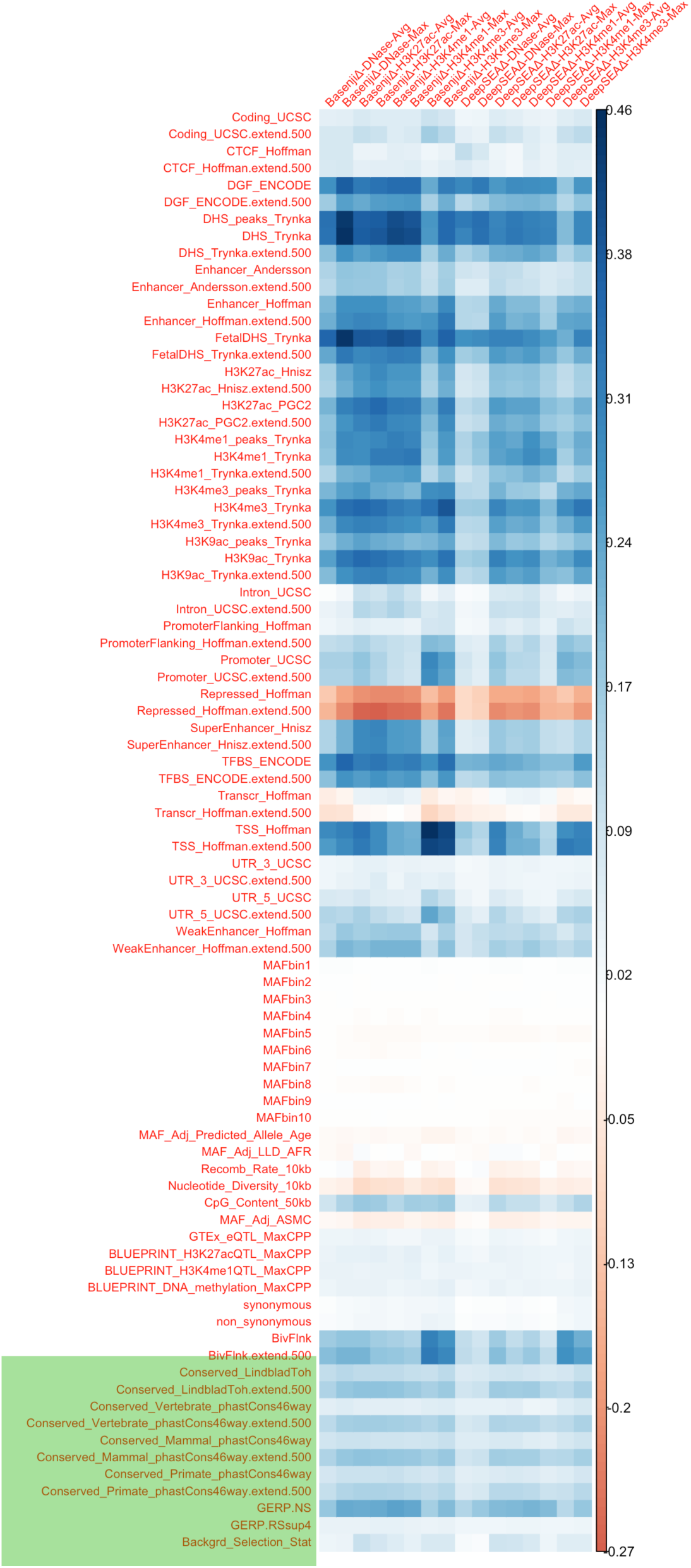
**Correlations between non-tissue-specific deep learning annotations and the baselineLD v2.1 annotations**. We report correlations for 16 non-tissue-specific allelic-effect deep learning annotations and the 85 baseline-LD annotations (excluding the annotation with all 1s). The 11 conservation related annotations are reported at the bottom of the table and shaded green.

**Figure S8.**
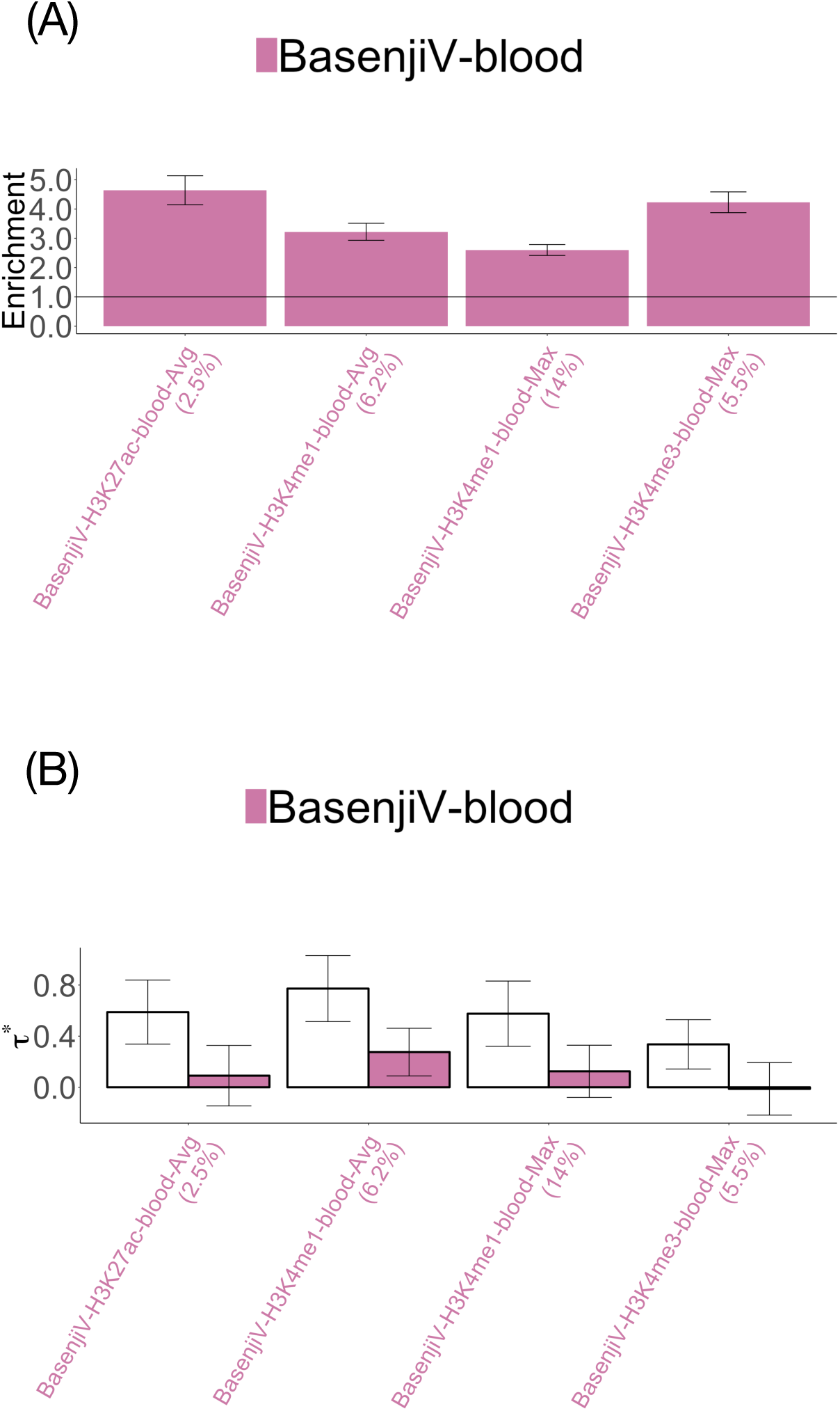
**Disease informativeness of blood-specific variant-level deep learning annotations**: (A) Heritability enrichment, conditioned on the non-tissue-specific variant-level joint model. Horizontal line denotes no enrichment. (B) Standardized effect size *τ\** conditioned on either the non-tissue-specific variant-level joint model (marginal analysis: left column, white) or the non-tissue-specific variant-level joint model plus 2 Roadmap and 4 ChromHMM blood-specific annotations (blood-specific variant-level joint model: right column, dark shading). Results are meta-analyzed across 11 blood-related traits. Results are displayed only for annotations with significant *τ\** in marginal analyses after correcting for 80 (variant-level + allelic-effect) blood-specific annotations tested (*P <* 0.05*/*80). None of these annotations were significant after conditioning on the blood-specific variant-level joint model. Error bars denote 95% confidence intervals. Numerical results are reported in Table S39 and Table S41.

**Figure S9.**
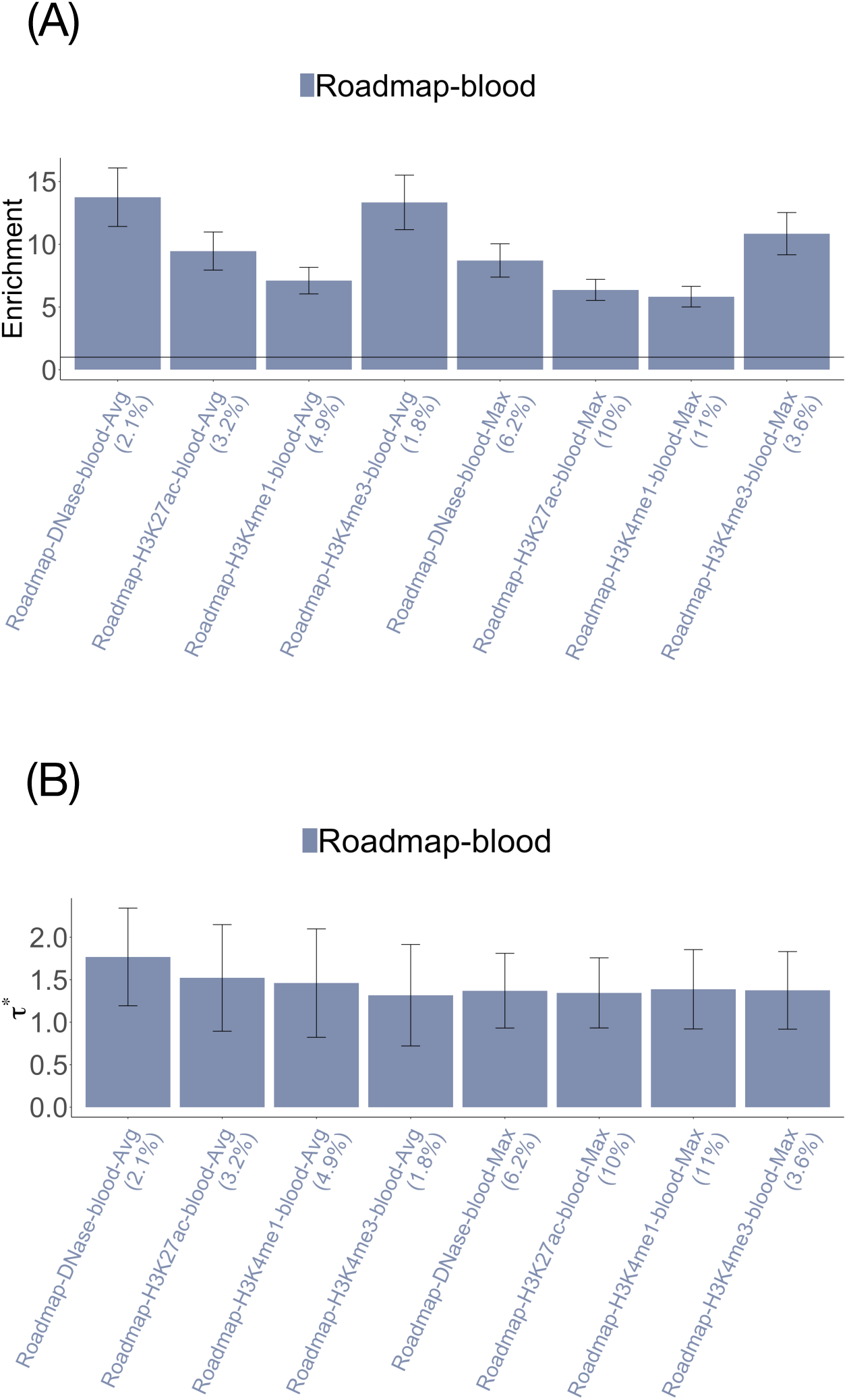
**Disease informativeness of blood-specific Roadmap annotations in marginal analysis conditional on the non-tissue-specific variant-level joint model**. (A) Heritability enrichment, conditioned on the non-tissue-specific variant-level joint model (original baseline-LD annotations and the annotations from Figure S6). Horizontal line denotes no enrichment. (B) Standardized effect size (*τ\**) conditioned on the non-tissue-specific variant-level joint model. Results are meta-analyzed across 11 blood-related traits. Results are displayed only for annotations with significant *τ^*^*in marginal analyses after correcting for 80 (variant-level + allelic-effect) blood-specific annotations tested (*P <* 0.05*/*80). Error bars denote 95% confidence intervals. Numerical results are reported in Table S40.

**Figure S10.**
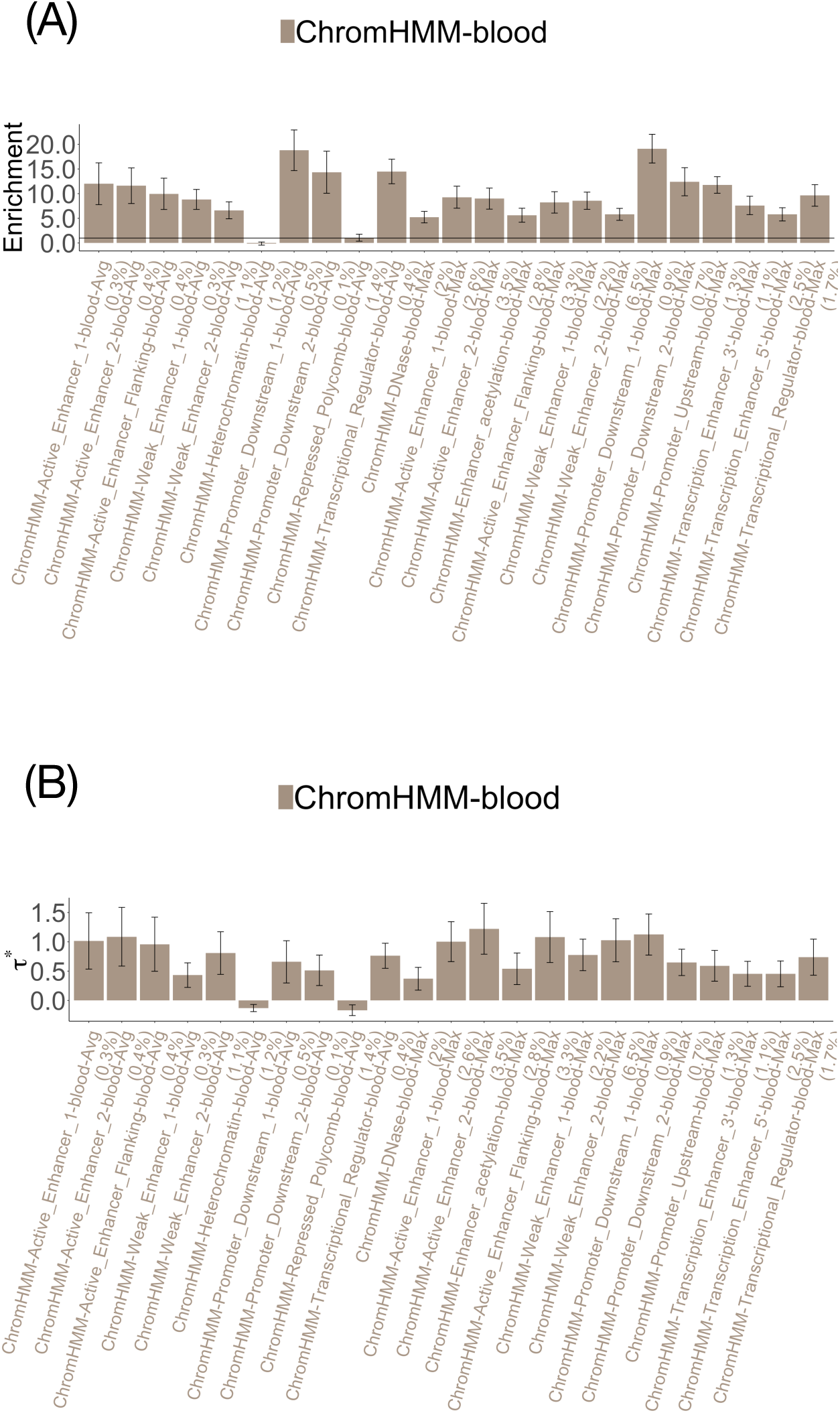
**Disease informativeness of blood-specific ChromHMM annotations in marginal analysis conditional on the non-tissue-specific variant-level joint model**. (A) Heritability enrichment, conditioned on the non-tissue-specific variant-level joint model (original baseline-LD annotations and the annotations from Figure S6). Horizontal line denotes no enrichment. (B) Standardized effect size (*τ\**)conditioned on the non-tissue-specific variant-level joint model. Results are meta-analyzed across 11 blood-related traits. Results are displayed only for annotations with significant *τ^*^*in marginal analyses after correcting for 80 (variant-level + allelic-effect) blood-specific annotations tested (*P <* 0.05*/*80). Error bars denote 95% confidence intervals. Numerical results are reported in Table S40.

**Figure S11.**
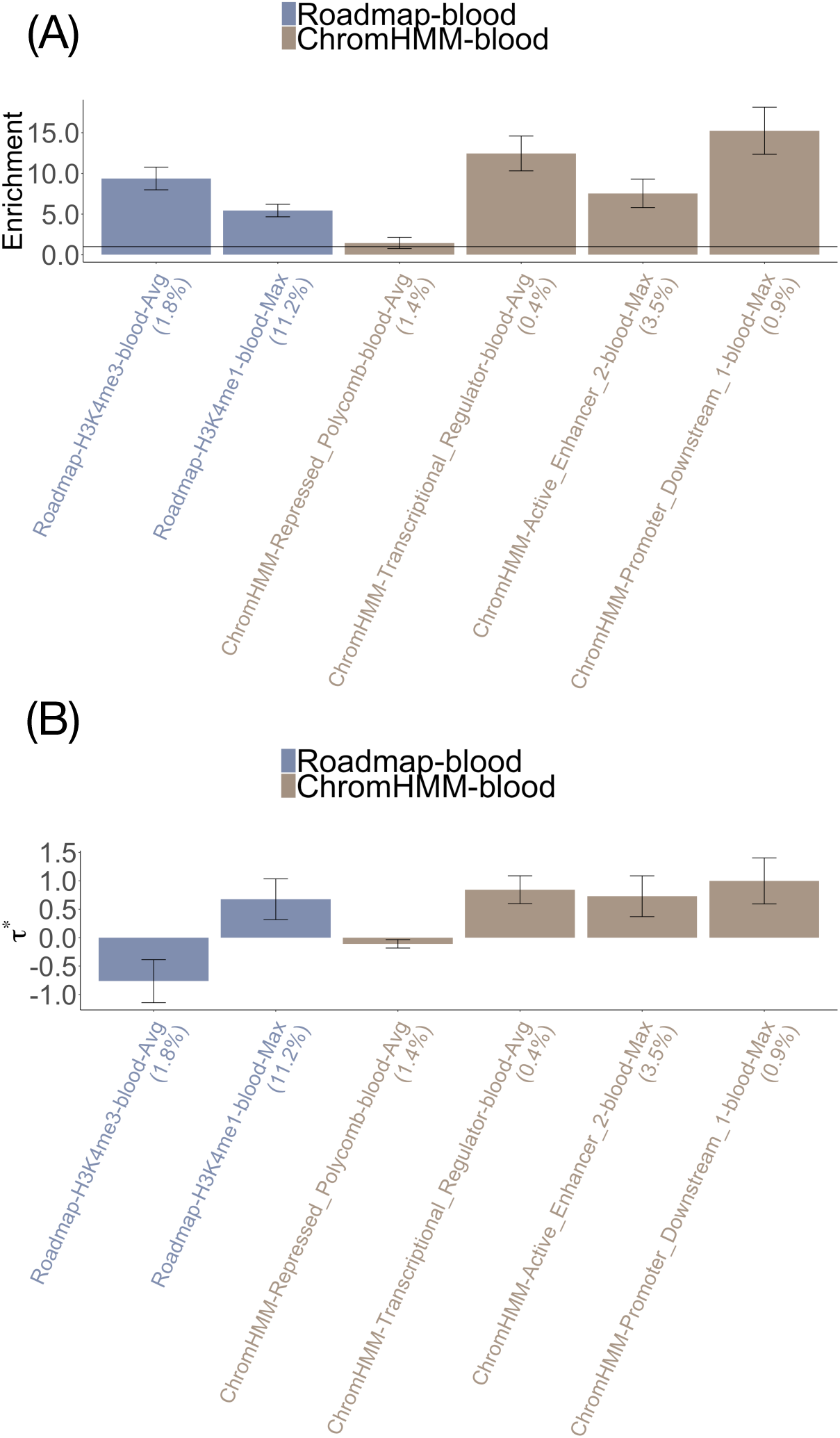
**Disease informativeness of blood-specific Roadmap and ChromHMM annotations in blood-specific final joint model**. (A) Heritability enrichment, conditioned on each other and non-tissue-specific variant-level joint model (original baseline-LD annotations and the annotations from Figure S6). Horizontal line denotes no enrichment. (B) Standardized effect size (*τ\**) conditioned on each other and the non-tissue-specific variant-level joint model. Results are meta-analyzed across 11 blood-related traits. Results are displayed only for annotations with significant *τ\** in marginal analyses after correcting for 80 (variant-level + allelic-effect) blood-specific annotations tested (*P <* 0.05*/*80). Error bars denote 95% confidence intervals. Numerical results are reported in Table S41.

**Figure S12.**
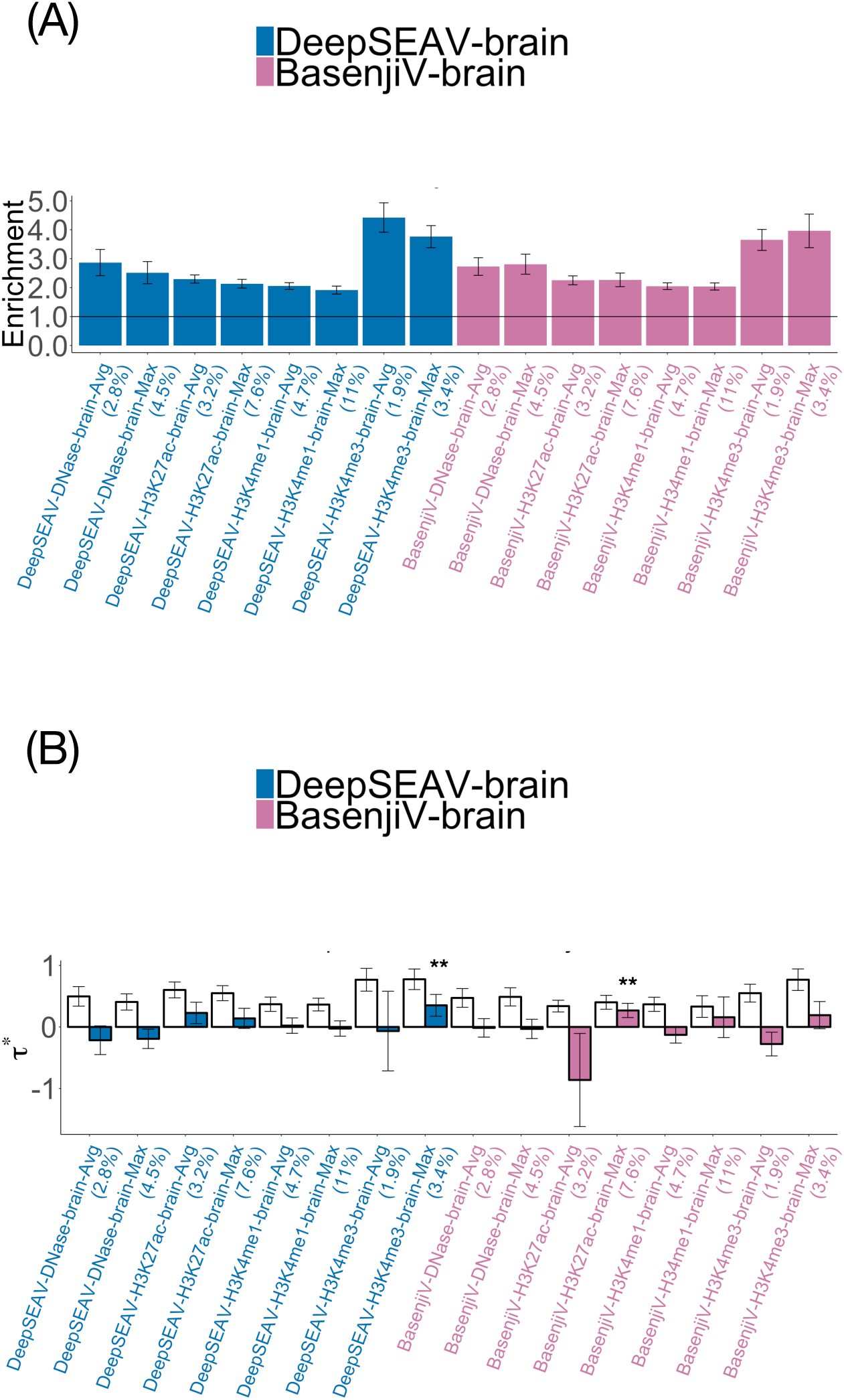
**Disease informativeness of brain-specific variant-level deep learning annotations**: (A) Heritability enrichment, conditioned on the non-tissue-specific variant-level joint model. Horizontal line denotes no enrichment. (B) Standardized effect size *τ\** conditioned on either the non-tissue-specific variant-level joint model (marginal analysis: left column, white) or the non-tissue-specific variant-level joint model plus 1 DeepSEA, 1 Basenji, 1 Roadmap and 3 ChromHMM brain-specific annotations (brain-specific variant-level joint model: right column, dark shading). Results are meta-analyzed across 8 brain-related traits. Results are displayed only for annotations with significant *τ\** in marginal analyses after correcting for 80 (variant-level + allelic-effect) brain-specific annotations tested (*P <* 0.05*/*80). For brain-specific variant-level joint model, ** denotes *P <* 0.05*/*80. Error bars denote 95% confidence intervals. Numerical results are reported in Table S45 and Table S47.

**Figure S13.**
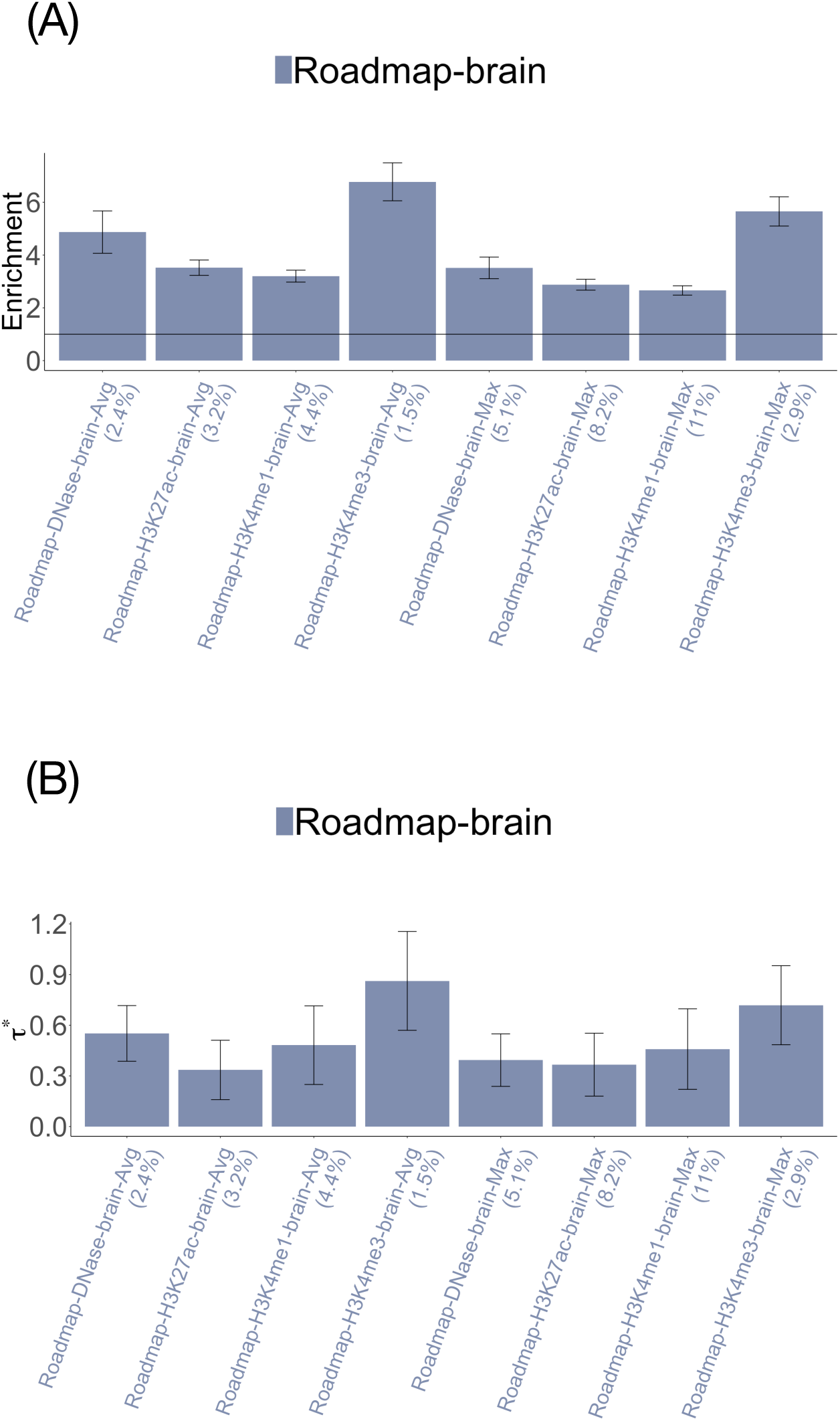
**Disease informativeness of brain-specific Roadmap annotations in marginal analysis conditional on the non-tissue-specific variant-level joint model**. (A) Heritability enrichment, conditioned on the non-tissue-specific variant-level joint model (original baseline-LD annotations and the annotations from Figure S6). Horizontal line denotes no enrichment. (B) Standardized effect size (*τ\**) conditioned on the non-tissue-specific variant-level joint model. Results are meta-analyzed across 8 brain-related traits. Results are displayed only for annotations with significant *τ\** in marginal analyses after correcting for 80 (variant-level + allelic-effect) blood-specific annotations tested (*P <* 0.05*/*80). Error bars denote 95% confidence intervals. Numerical results are reported in Table S46.

**Figure S14.**
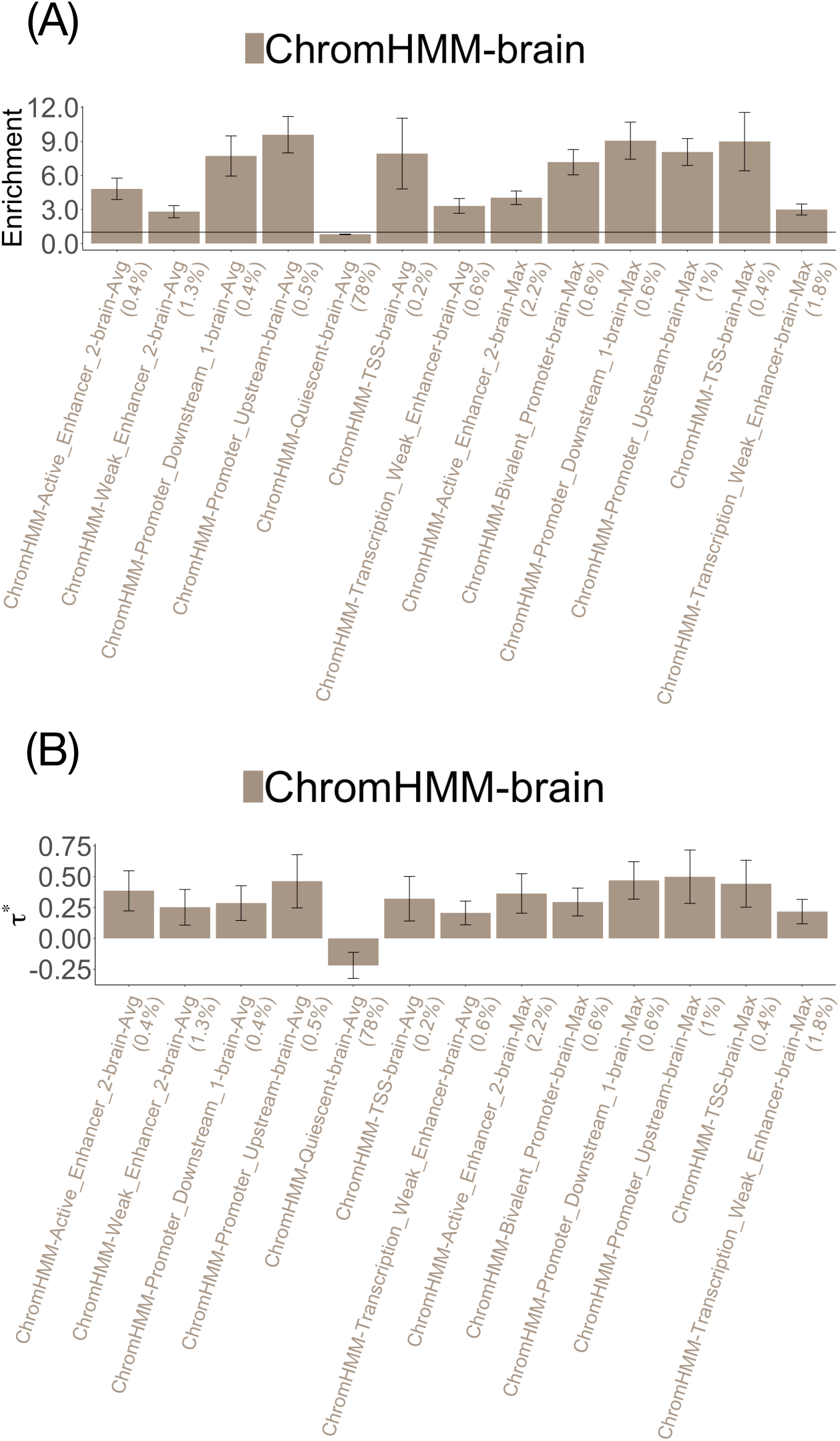
**Disease informativeness of brain-specific ChromHMM annotations in marginal analysis conditional on the non-tissue-specific variant-level joint model**. (A) Heritability enrichment, conditioned on the non-tissue-specific variant-level joint model (original baseline-LD annotations and the annotations from Figure S6). Horizontal line denotes no enrichment. (B) Standardized effect size (*τ\**) conditioned on the non-tissue-specific variant-level joint model. Results are meta-analyzed across 8 brain-related traits. Results are displayed only for annotations with significant *τ\** in marginal analyses after correcting for 80 (variant-level + allelic-effect) blood-specific annotations tested (*P <* 0.05*/*80). Error bars denote 95% confidence intervals. Numerical results are reported in Table S46.

**Figure S15.**
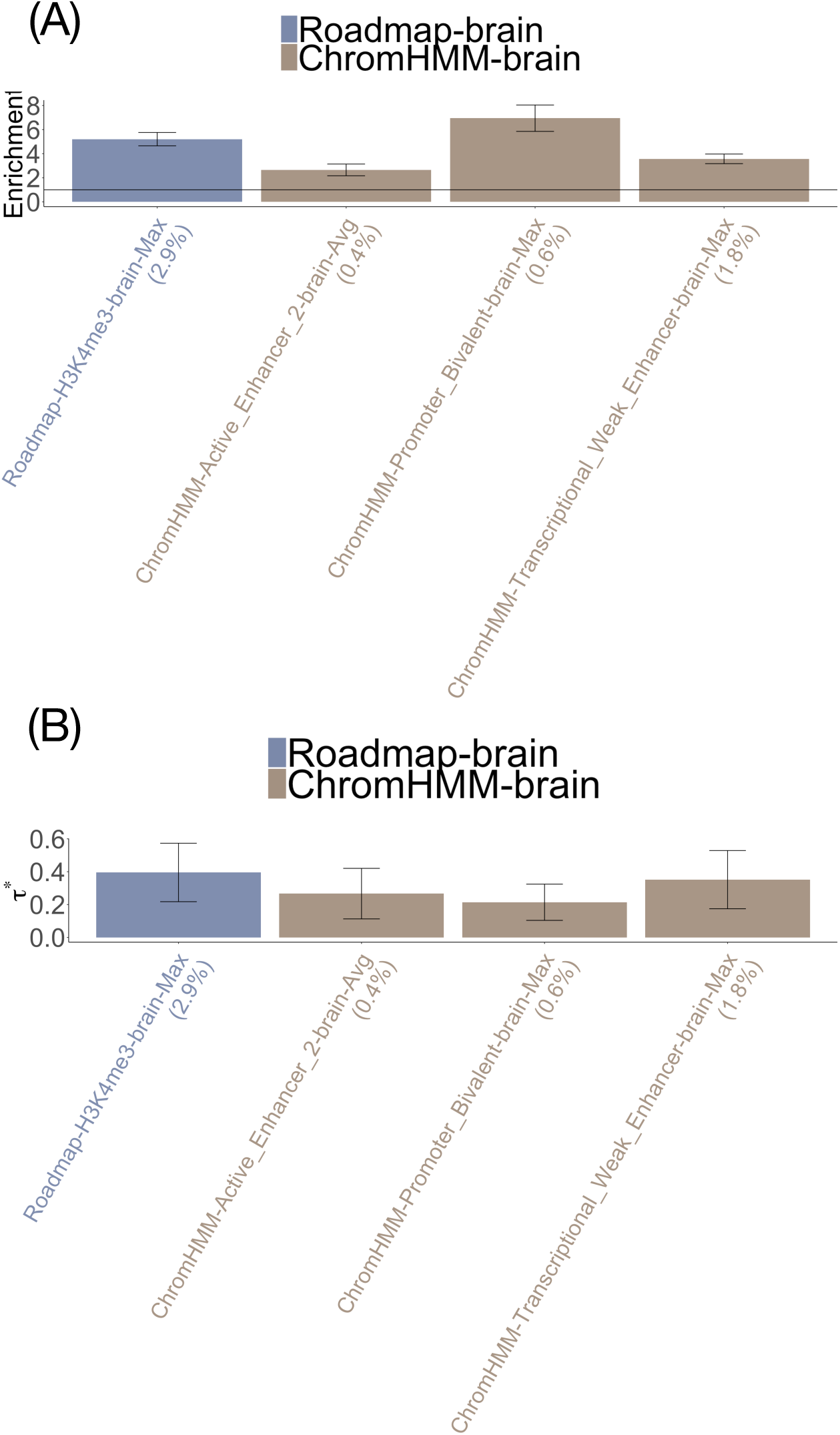
**Disease informativeness of brain-specific Roadmap and ChromHMM annotations in brain-specific final joint model**. (A) Heritability enrichment, conditioned on the non-tissue-specific variant-level joint model (original baseline-LD annotations and the annotations from Figure S6), and all brain-specific variant-level DeepSEA and Basenji, and Roadmap and ChromHMM annotations. Horizontal line denotes no enrichment. (B) Standardized effect size (*τ\**) conditioned on the non-tissue-specific variant-level joint model and all brain-specific variant-level DeepSEA and Basenji, and Roadmap and ChromHMM annotations. Results are meta-analyzed across 8 brain-related traits. Results are displayed only for annotations with significant *τ^*^*in marginal analyses after correcting for 80 (variant-level + allelic-effect) blood-specific annotations tested (*P <* 0.05*/*80). Error bars denote 95% confidence intervals. Numerical results are reported in Table S47.

**Figure S16.**
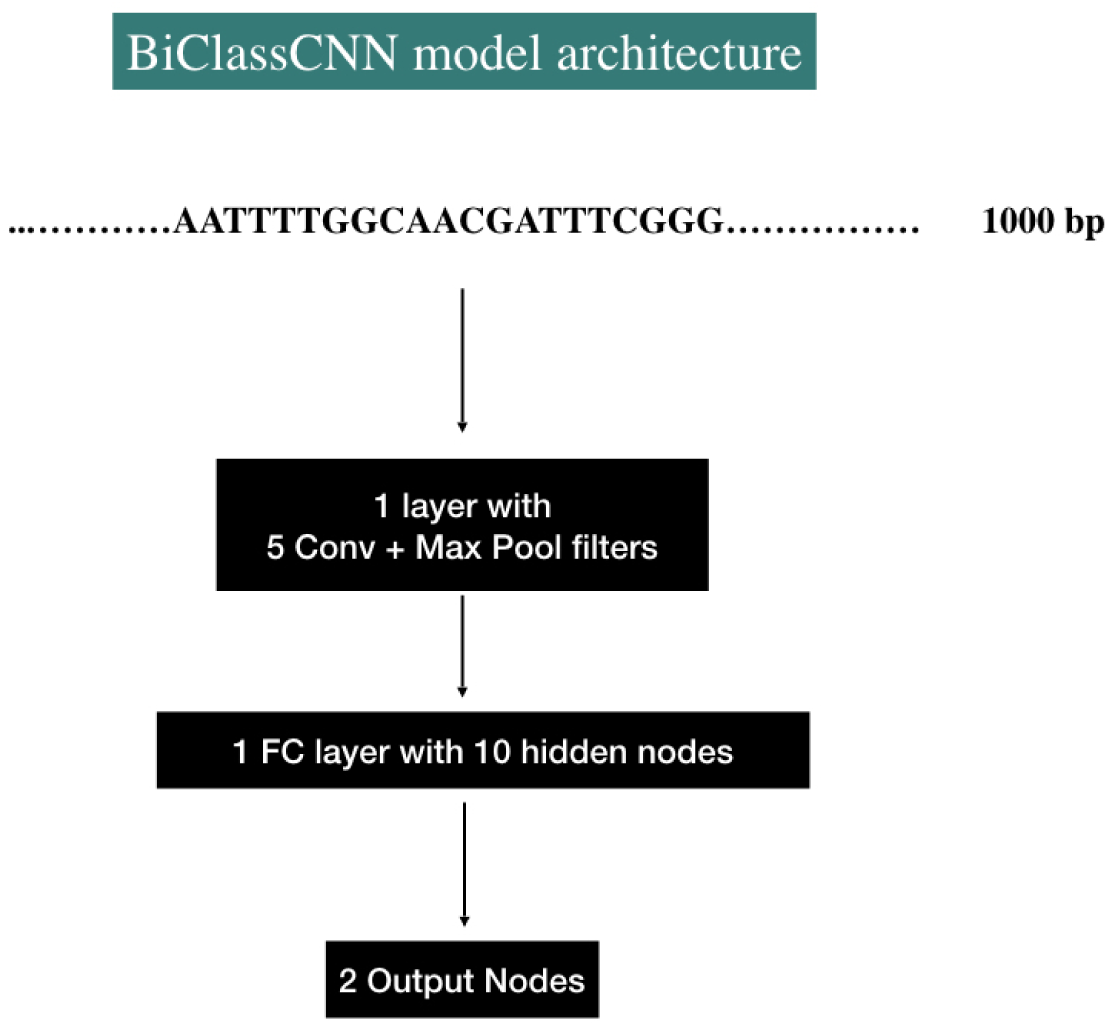
**The Neural net architecture for BiClassCNN**. The architecture of the Covolutional Neural Net (CNN) model used for BiClassCNN training. It comprioses of two layers - 1 layer being Convolutional+MaxPool with 5 filters and 1 Fully Connected hidden layer with 10 nodes leading to the final output layer with 2 nodes distinguishing between the positive and negative sets. We chose the convolutional filter to be of size 4 and stride length 1 and the MaxPool filter to pool every 4 bases at a time. RELU activation was used.

**Figure S17.**
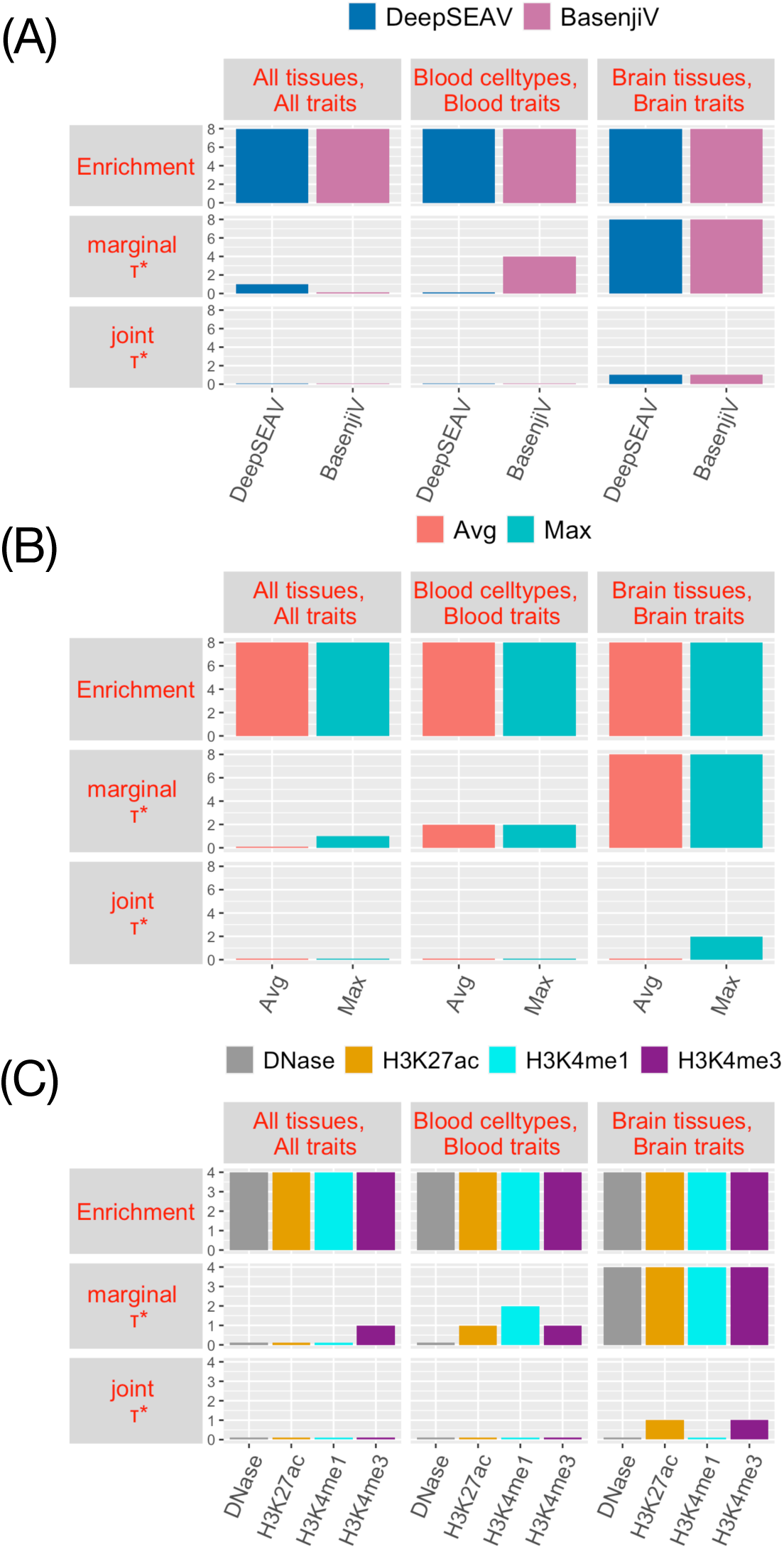
**Number of significantly disease informative variant-level deep learning annotations**: Number of significant non-tissue-specific, blood-specific and brain-specific variant-level annotations across (A) different deep learning models (DeepSEA and Basenji), (C) different aggregation strategies (Average and Maximum) and (C) different types of epigenomic marks, in terms of marginal enrichment in heritability, conditional *τ^*^*and joint *τ\** for all traits, blood traits and brain traits respectively. For all traits analysis, the analysis is conditioned on the baseline-LD model. For blood traits analysis, the analysis is conditioned on non-tissue-specific variant-level joint model. For brain traits analysis, the analysis is conditioned on non-tissue-specific variant-level joint model. Numerical results are reported in Table S28 (numerical summary of results), Table S29 (enrichment and marginal *τ\** for all tissues, all traits analysis), Table S34 (joint *τ\** of all tissues, all traits analysis), Table S39 (enrichment and marginal *τ\** for blood cell ty1p7es, blood traits analysis), Table S41 (joint *τ\** of blood cell types, blood traits analysis), Table S45 (enrichment and marginal *τ\** for brain tissues, brain traits analysis) and Table S47 (joint *τ^*^*of brain tissues, brain traits analysis).

## Notes

https://data.broadinstitute.org/alkesgroup/LDSCORE/DeepLearning/

### Competing Interest Statement

The authors have declared no competing interest.

### Summary of Updates

We have moved the variant-level annotation study to the Supplementary Note upon reviewer request. We have performed some additional comparison of the allelic-effect annotations with conservation-related terms. We have expanded on the downstream motif based analysis. We have toned down the results section considerably.

